# Dimerization and Threonine-Dependent Stabilization Govern Human YRDC Catalysis

**DOI:** 10.64898/2025.12.03.692196

**Authors:** Thư Ngọc-Anh Trịnh, Greice M. Zickuhr, Alison L. Dickson, Silvia Synowksy, Sally L. Shirran, Carlin J. Hamill, Ramasubramanian Sundaramoorthy, Tomas Lebl, David J. Harrison, Clarissa M. Czekster

## Abstract

Addition of *N*^6^-threonylcarbamoyladenosine occurs at position 37 (t⁶A_37_) of five mitochondrial tRNA species, influencing translation fidelity and efficiency. Mutations in YRDC, the enzyme catalyzing the first step in t⁶A_37_ synthesis, cause severe neurological and renal diseases in humans, including in Galloway-Mowatt disease (GAMOS). YRDC generates threonylcarbamoyl-AMP (TC-AMP) using ATP, threonine and bicarbonate as substrates, yet their binding order, residues involved in amino-acid selectivity, and precisely how mutations contribute to disease remain unclear. Here, we combine protein biophysics, mass spectrometry, NMR, mutagenesis and kinetics to define how the human enzyme operates. Differential scanning fluorimetry and circular dichroism identify L-threonine as the gatekeeper ligand. It binds with millimolar affinity, stabilizing YRDC and enhancing ATP binding nearly fortyfold, enabling bicarbonate association. Several amino acids can form aminoacylcarbamoyl-AMP adducts in vitro, but threonine yields superior protein stabilization and product formation. Native mass spectrometry and single-molecule mass photometry corroborate YRDC’s dimeric state. GAMOS-linked mutations destabilize folding, disrupting dimerization and uncoupling substrate binding from catalysis, explaining functional losses. Supported by kinetic data and NMR, we propose a chemical mechanism for the reaction and identify dimerization and conformational stability as control points for activity. This work provides a foundation for designing selective YRDC inhibitors or enhancers.

## Introduction

Post-transcriptional modification of RNA is a fundamental regulatory mechanism that controls its structure, stability, and function. Transfer RNAs (tRNAs) show the greatest diversity, carrying over 90 distinct chemical modifications across phylogenetic domains^1,2^. These are enriched in functionally critical regions of tRNAs, playing essential roles in maintaining the fidelity and efficiency of translation^3–7^. Mammalian mitochondrial DNA (mt-DNA) encodes 22 tRNA species that are required for the translation of 13 polypeptides which are the core subunits of the oxidative phosphorylation complexes^8^. mt-tRNAs undergo extensive post-transcriptional modification, with > 70% of tRNA species modified at position 37 to enhance codon–anticodon pairing, stabilize the anticodon stem–loop (ASL) domain structure, prevent translational frameshifting, and promote decoding efficiency^9^. Disruption of these modification pathways, especially in the mitochondria, is a central mechanism underlying translational failure and human disease^10–13^.

The N⁶-threonylcarbamoyladenosine (t⁶A_37_) modification at position 37 of tRNAs is biologically essential and shared across all domains of life^14–19^. In mammalian mitochondria, t⁶A_37_ is found in tRNAs that correspond to serine, threonine, asparagine, isoleucine, and lysine^20^. Modification loss is lethal in *Escherichia coli* and leads to elevated translational errors and significantly impaired growth in *Saccharomyces cerevisiae*^5,20–25^.

Biosynthesis of t⁶A_37_ proceeds via two consecutive enzymatic steps. First, threonylcarbamoyl-adenylate (TC-AMP) is generated from L-threonine, ATP, and a single-carbon donor proposed to be dissolved bicarbonate. In the second step, the threonylcarbamoyl group is transferred from TC-AMP to the N⁶ position of adenosine at position 37 of the tRNA^24,26–31^. Recent studies have identified YRDC (the yeast *Sua5* homolog), which is distributed in both the cytoplasm and mitochondria, and OSGEPL1 (the yeast *Kae1/Qri7* homolog) as the two enzymes responsible for inserting the mitochondrial t⁶A_37_ modification in humans^32,33^ (Figure 1). Cancer Dependency Map analyses reveal it is essential for the survival of most cancer cell lines, with minimal impact on non-malignant cells^34^. YRDC inhibition lowers t⁶A_37_ levels, suppresses global translation, and impedes tumor growth in vitro and in vivo^35,36^. Threonine restriction, targeting the t⁶A_37_ pathway, similarly reduces tumor size, slows xenograft progression, and enhances chemotherapeutic efficacy^34^. Pathogenic YRDC variants have recently been associated with Galloway–Mowat syndrome (GAMOS)^12^. The clinical spectrum of GAMOS is heterogeneous, but the prognosis is poor, with most affected individuals dying before six years of age. To date, ten GAMOS subtypes (GAMOS 1–10) have been defined, each linked to mutations in distinct genes. A subset of patients with YRDC mutations present with GAMOS10, a particularly severe subtype of GAMOS ^12,37^.

**Figure 1.**
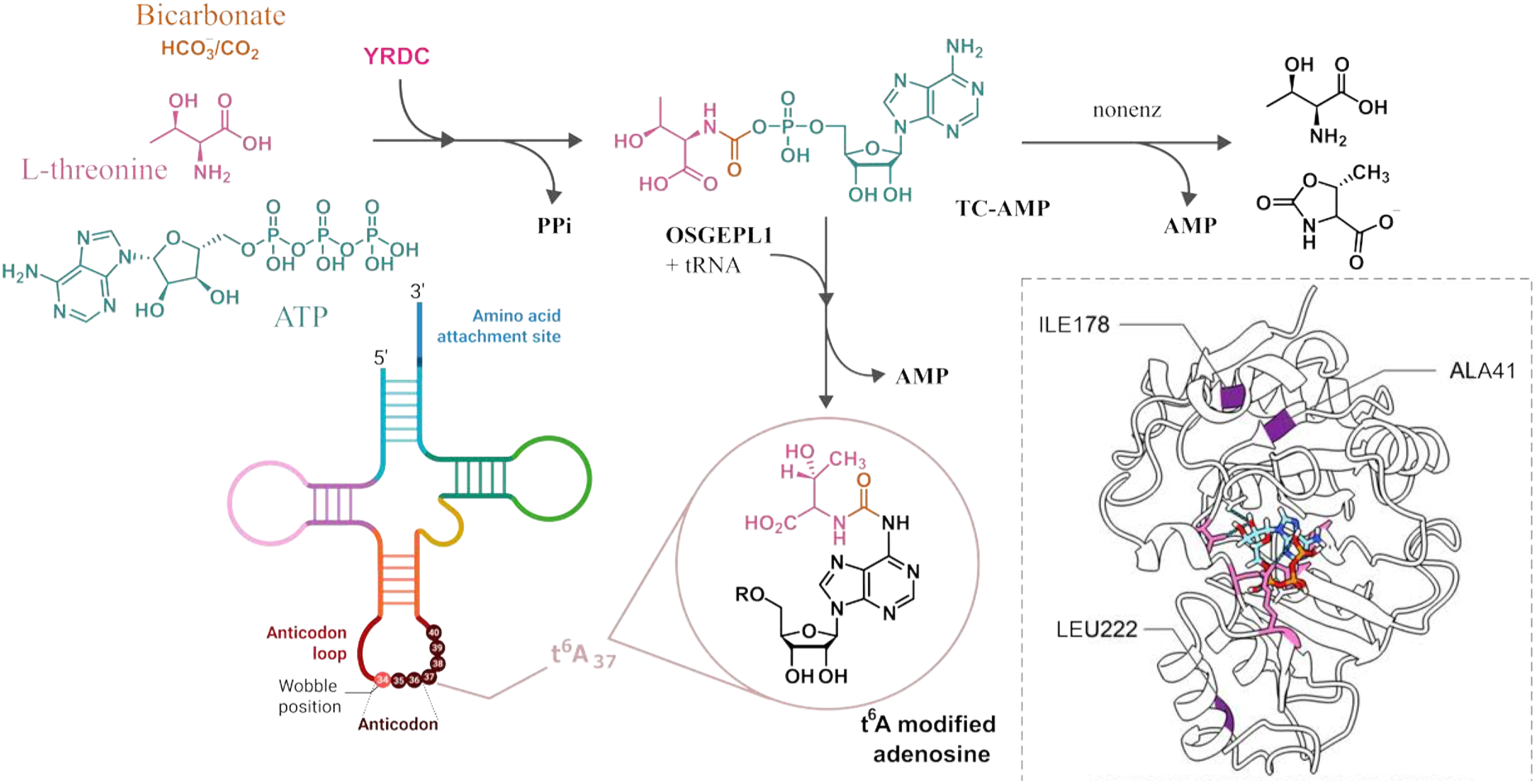
Threonylcarbamoyl adenosine (t⁶A) formation in humans. **Top**: L-threonine and bicarbonate, together with ATP, are converted by YRDC into the activated intermediate threonylcarbamoyl-AMP (TC-AMP) with release of one pyrophosphate (PPi) molecule. OSGEPL1 then transfers the threonylcarbamoyl group from TC-AMP to A37 in the tRNA anticodon loop, yielding t⁶A and AMP; a competing non-enzymatic decay of TC-AMP to AMP and threonine (or threonine-oxazolone) is indicated. **Bottom right**: Cartoon representation of a monomer of human YRDC lacking the N-terminal 43 residues (ΔN43) as predicted by AlphaFold3. Residues implicated in Galloway–Mowat syndrome (GAMOS) are highlighted in purple, and residues selected for binding-site mutagenesis are highlighted in pink. Residue numbering corresponds to the ΔN43 construct used in this study.

Despite its critical role in t⁶A_37_ biosynthesis, association with multiple cancers and with a lethal neuro-renal syndrome, prior to this work the catalytic and kinetic mechanisms of YRDC and precisely how mutations lead to loss in function remained undetermined. Here, informed by comparative analysis with its closest homolog SUA5, we performed site-directed mutagenesis of YRDC and identified key amino acids in L-threonine recognition. In addition, differential scanning fluorimetry (DSF) and circular dichroism (CD) defined a stabilising role for L-threonine and the order of substrate binding. We further showed that other amino acids such as L-Cys, L-Ser, L-Ala, and L-Val can be substrates for YRDC, albeit with lower product formation. Kinetics of YRDC comparing active-site and GAMOS-related mutants describe little catalytic efficiency loss, but larger losses in L-threonine binding. These findings reveal critical active-site interactions, define the effects of GAMOS-related mutations, and quantify the impact of individual substrates on protein stability. Combining structural predictions and our biophysical data employing native mass spectrometry, size exclusion chromatography and mass photometry (MP) we conclude that wild type YRDC exists in a dimeric form and that GAMOS-related mutations interfere with its ability to dimerise, thereby affecting enzymatic activity.

## Results

YRDC possesses a weak mitochondrial targeting sequence (MTS), which results in a considerable proportion of the protein remaining in the cytoplasm, where it participates in the formation of t^6^A_37_ in cytoplasmic tRNAs. In contrast, a smaller fraction of YRDC is imported into mitochondria, where it carries out the same function in mitochondrial tRNAs. Previous work has shown that truncation of up to 52 amino acids from the N-terminus does not significantly impair the enzymatic activity of YRDC^32^. Therefore, in all experiments carried out here the protein was purified in a truncated form lacking the first 43 amino acids (ΔN43 construct, referred to subsequently as YRDC_WT_). Supplementary Figure 1 shows the purified proteins utilised here.

### L-threonine increases T_m_ and enhances ATP binding

In DSF assays (Figure 2, Supplementary Figure 2, Supplementary Table 2), YRDC_WT_ displayed weak affinity for ATP, with an equilibrium dissociation constant (*K*_D_) of approximately 2 mM. The presence of L-threonine enhanced ATP binding more than ten-fold, reducing the *K*_D_ to 77.3 ± 29.4 µM (Figure 2e, Supplementary Table 2). A comparable effect was observed with bicarbonate (HCO₃⁻): no binding of HCO*₃*⁻ was observed for YRDC_WT_ by DSF, but the inclusion of L-threonine yielded a *K*_D_ of 38.3 ± 12.9 mM (Figure 2g). Consistent with this, YRDC_WT_ can bind L-threonine alone, with a *K*_D_ of 31.1 ± 12.1 mM (Figure 2f), corroborated by NMR (below). These suggest that binding of L-threonine may facilitate access of ATP and bicarbonate to the protein’s binding site. In addition, the presence of ATP did not appreciably affect the binding affinity for L-threonine (*K*_D_ = 28.4 ± 12.3 mM). ATP had little or slight effect on the stability of the L-threonine–YRDC complex, as evidenced by an increase in the T_m_ value from 55 °C with 50 mM L-threonine alone to 59 °C with 25 mM L-threonine in an ATP-saturated medium. The combined presence of ATP and L-threonine led to a marked enhancement in bicarbonate binding, reducing the *K*_D_ to 14.6 ± 5.6 mM (Figure 2g). Together, these data indicate that L-threonine acts as a molecular gatekeeper, substantially increasing YRDC’s affinity for its other ligands. Additionally, DSF analyses revealed that YRDC_WT_ binds to pyrophosphate and AMP, both metabolic by-products of the reaction and potential inhibitors, with *K_D_* values of 7.1 ± 0.9 mM and 6.1 ± 1.8 mM, respectively (Supplementary Figure 3).

**Figure 2.**
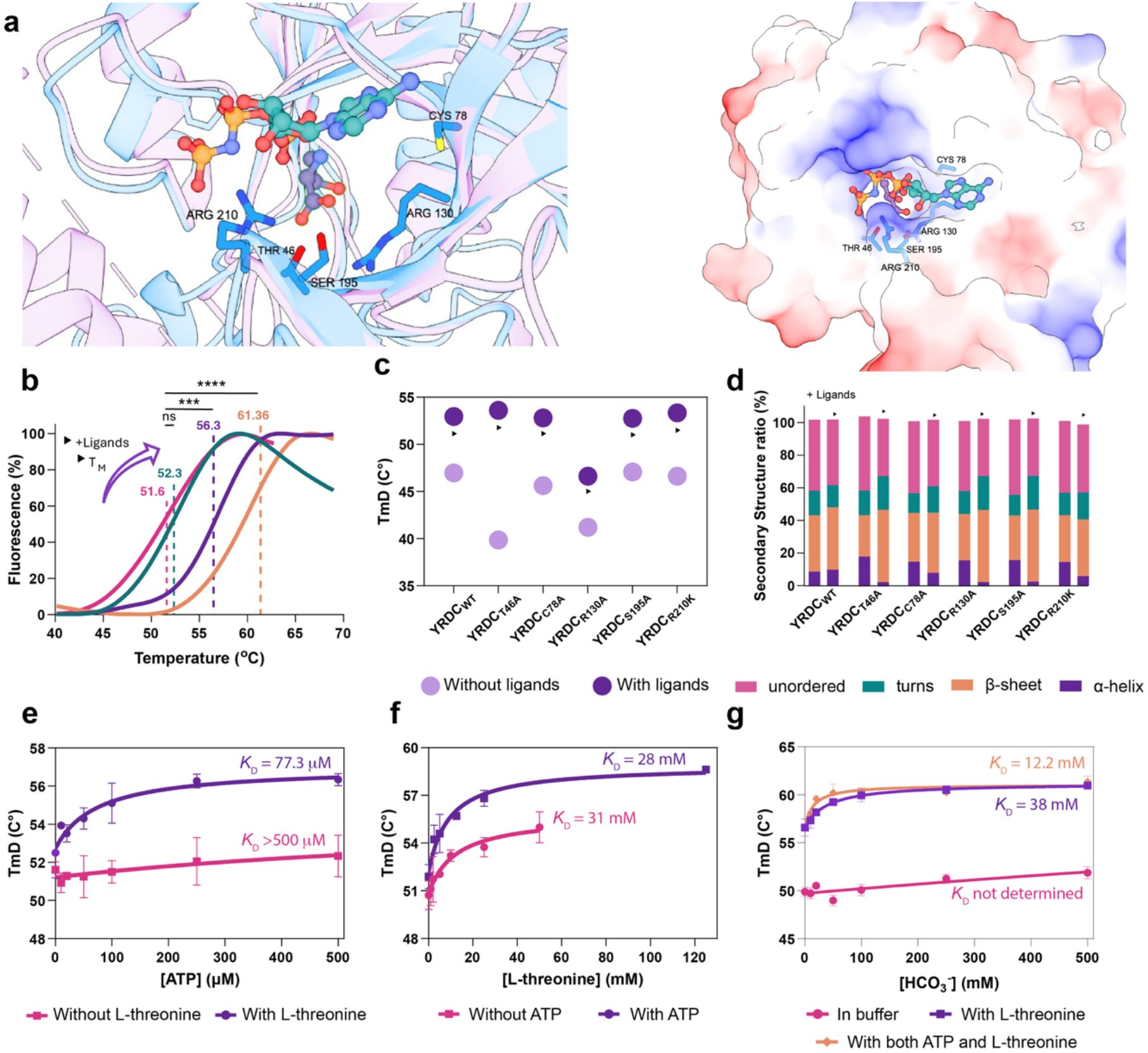
Structural model of YRDC (ΔN43), biophysical characterization of binding-site mutants and ligand effects. **a** Structural overlay of the AlphaFold3 prediction for human YRDC lacking the N-terminal 43 residues (ΔN43; Q86U90) with the homolog SUA5 from *Sulfurisphaera tokodaii* (PDB 4E1B) to orient the binding pocket. Human YRDC is shown in blue, ligands AMP-PNP (teal) and threonine (purple) from SUA5 are shown as ball stick. The five YRDC residues mutated here are shown as sticks (Thr46, Cys78, Arg130, Ser195, and Arg210). Right: based on SUA5, threonine is predicted to be buried deep in its binding site, with AMP-PNP closer to the surface, mutated residues are in proximity to threonine. **b** DSF melting curves and fitted melting temperatures (Tₘ) obtained by a derivative method fit. From left to right, the traces correspond to apo protein (purple) and to protein in the presence of ATP (500 µM; green), L-threonine (50 mM; yellow), and NaHCO₃ (50 mM; pink). Tₘ values are displayed as means (n = 3). Statistical comparisons were performed by one-way ANOVA with a 95% confidence interval; significance is annotated as P < 0.001 (***), P < 0.0001 (****), and not significant (P > 0.05; ns). **c** Melting temperatures of YRDC_WT_ and binding-site mutants measured by DSF with (pink) and without (green) ligands (ATP 500 µM and L-threonine 50 mM). Measurements were acquired over 20–95 °C under identical buffer conditions. Arrows indicate the direction and magnitude of the ΔTₘ shift between ligand-free and ligand-bound states. Data are mean (n = 3). **d** Secondary-structure composition (α-helix, β-sheet, turns, unordered) for YRDC_WT_ and the binding-site mutants derived from CD. Fractional contents were estimated using the SELCON3 algorithm implemented on the ChiraKit^38^ platform. Bars report percentages; bars on the left are without ligand and the bar next to its right with ligands (ATP 500 µM *and L-t*hreonine 50 mM). **e–g**, Concentration–stability relationships for YRDC_WT_ by DSF in the presence of increasing concentrations of ATP (e), L-threonine (f), and bicarbonate (g). Values are presented as mean ± s.e.m (n = 3).

For the majority of YRDC mutants examined here (Figure 2a), including those with alterations at the ligand-binding site, DSF traces revealed either unstable profiles or characteristic of unfolded conformations. Remarkably, the addition of 50 mM L-threonine substantially improved both the thermal stability (Figures 2b and 2c) and folding profiles of these mutants, with the YRDC_C78A_ variant showing particularly enhanced stabilization. Additionally, consistent with the behaviour of the WT protein, none of the mutants demonstrated detectable ATP binding in the absence of L-threonine. Variants YRDC_T46A_ and YRDC_R130A_ showed little or no ligand binding compared to wild type. In contrast, the YRDC_S195A_ and YRDC_R210K_ mutants retained the ability to bind ATP in the presence of L-threonine, with *K*_D_ values of 48.9 ± 37.8 µM and 147.5 ± 111.8 µM, respectively. Both mutants were able to bind L-threonine in the presence of 500 µM ATP, with *K*_D_ values of 17.2 ± 9.8 mM for YRDC_S195A_ and 3.4 ± 1.8 mM for YRDC_R210K_.

### Ligands influence YRDC secondary structure and stability

To determine changes in secondary structure upon ligand binding and confirm whether protein variants were folded, CD spectra were recorded for YRDC_WT_ and mutants (Figure 2d, Supplementary Figure 4) in the absence and presence of ligands (500 µM ATP and 5 mM L-threonine). Ligand binding reduced the proportion of disordered regions and increased β-sheet content (Supplementary Table 3). YRDC_WT_ showed the highest T_m_ of 47.0 ± 0.3 degrees, consistent with DSF values (comparison on Supplementary Table 1). Despite ligand binding reducing the total random coil content (comprising turns and disordered regions) in all cases, the mutants retained a higher disordered fraction and exhibited significantly lower β-sheet content compared to YRDC_WT_. The α-helical content in the apo form of binding-site mutants was nearly double that of the WT (8.5%). However, upon ligand binding, the α-helical fraction was markedly reduced, falling to as low as 2.6% in some, resulting in values significantly below those of the WT in the ligand-bound state (9.6%) (Supplementary Table 3, Supplementary Figure 4).

Consistent with the CD data, thermal stability measurements from DSF assays showed a clear ligand-induced increase in melting temperature (T_m_) of WT (Figure 2e-g), rising from 47.0 ± 0.3 °C to 53.0 ± 0.2 °C, and from 49.9 ± 0.5°C to 59.4 ± 0.5°C in the absence and presence of ATP and L-threonine, respectively. Similar trends were observed for binding-site mutants (Supplementary Table 1).

#### Single point mutations in YRDC severely impact catalytic activity

Using an LC-MS assay quantifying AMP formation as a proxy for TC-AMP formation due to its instability and decay into AMP (see Supplementary methods “Establishment of an LC-MS assay to determine the activity of YRDC”), kinetic parameters for each substrate involved in TC-AMP formation were determined (Figure 3, Supplementary Table 4). Negative control reactions lacking either L-threonine or bicarbonate had no AMP signal detected, showing that ATP was not converted to AMP in the absence of these substrates (confirmed by NMR, Supplementary Figure 5). Furthermore, negative controls showed no TC-AMP or AMP were detected unless all three substrates were present, confirming that AMP quantification reflects the formation of the intermediate TC-AMP. Michaelis–Menten analysis revealed that ATP exhibited a *K*_m_ of 73.6 µM (identical within experimental error to the YRDC_WT_ *K*_D-ATP_ = 77.3 µM) while L-threonine and bicarbonate showed *K*_m_ values of 7.8 mM and 3.3 mM, respectively, suggesting that elevated cellular concentrations of the latter two substrates are required to drive TC-AMP formation in vivo. The catalytic efficiencies (*k_cat_*/*K*_m_) of YRDC_WT_ for ATP, L-threonine, and bicarbonate were determined to be 0.3 mM^−1^s^−1^, 0.003 mM^−1^s^−1^, and 0.005 mM^−1^s^−1^ respectively (Supplementary Table 4). All data for enzymatic reactions are shown on Supplementary Figure 6.

**Figure 3.**
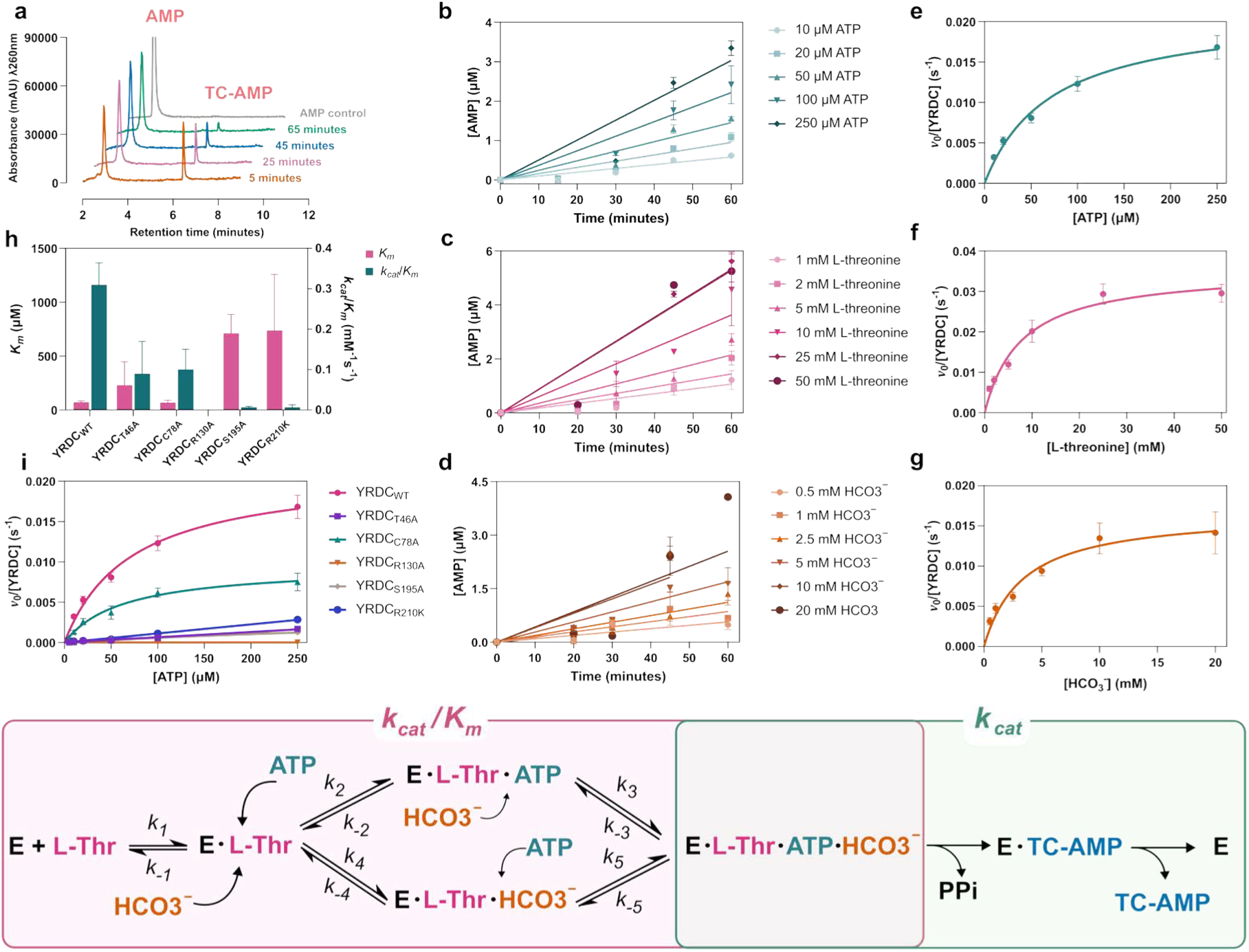
Steady-state kinetics of YRDC WT and binding-site mutants. **a** TC–AMP decay to AMP monitored by LC–MS. The appearance of AMP (control was highlighted in purple) was tracked beginning 5 min after initiating the reaction and then every 20 min for 1 h under identical buffer conditions. Experiments were performed in triplicate, and the curve shown represents the mean of three independent measurements (n = 3) at each time point. **b–d** Steady-state kinetics of YRDC_WT_ quantified from AMP formation resulting from TC–AMP turnover while varying one substrate at a time: ATP (b), *L*-threonine (c), and NaHCO₃ (d). Values reported as mean ± s.d. (n = 3). **e-g** Michaelis–Menten curves corresponding to the initial-rate data in b–d. Data shown as mean ± s.e.m. (n = 3) from independent experiments; solid lines show a fit to a Michaelis–Menten model. **h** Summary of kinetic parameters for YRDC_WT_ and five binding-site point mutants obtained by varying ATP from 10 µM to 250 µM. The Michaelis–Menten constant (*K*_m_) and catalytic efficiency (*k_cat_/K*_m_) for each protein are highlighted in green and pink, respectively. Plots show mean ± s.d. (n = 3). **i** Michaelis–Menten plots comparing the overall catalytic efficiency of the mutants with YRDC_WT_. Data were fitted to a Michaelis–Menten equation (three independent replicates, data as mean ± s.d.). n.d = parameters not determined within the tested ATP range. **Bottom**: Reaction scheme for the reaction catalysed by YRDC. Steps included in *k_cat_*/*K*_m_ are highlighted by a pink box, and those contributing to *k_cat_* highlighted by a green box.

By comparison to the closest structurally characterised homologue of human mitochondrial YRDC - SUA5 from *Sulfurisphaera tokodaii* (PDB: 2EQA), potential residues critical for substrate binding were mutated to generate variants YRDC_T46A_, YRDC_C78A_, YRDC_R130A_, YRDC_S195A_, and YRDC_R210K_.^25^ With the exception of YRDC_C78A_ all mutants exhibited a near-complete loss of catalytic activity, highlighting their essential role (Supplementary Table 5). Interestingly, the YRDC_C78A_ mutant displayed a marginally lower *K*_m_ for ATP (70 µM), yet its overall catalytic efficiency (*k_cat_*/ *K*_m_) was reduced by approximately three-fold, indicating impaired turnover rather than altered substrate affinity. For the other binding-site substitutions, substrate affinity and efficiency deteriorated markedly. The apparent *K*_m_ increased by three-fold for YRDC_T46A_ and by about ten-fold for YRDC_S195A_ and YRDC_R210K_ relative to wild type. In parallel, the *k_cat_*/*K*_m_ values fell by about three-fold for YRDC_T46A_ and by over forty-fold for YRDC_S195K_ and YRDC_R210K_. In the R130 mutant, neither the by-product AMP nor the intermediate TC-AMP were detected, suggesting a complete loss of catalytic activity and underscoring the essential role of this residue in substrate recognition and/or catalysis.

#### YRDC can accept other amino acids as substrates

To evaluate the substrate specificity of human YRDC and determine whether L-threonine is uniquely recognised, a panel of structurally and electrostatically similar amino acids, including L-cysteine, L-serine, L-alanine, and L-valine, was selected for comparative analysis by DSF (Figure 4a). YRDC_WT_ displayed a slightly higher binding affinity for *L*-cysteine than for of L-threonine, with *K*_D_ value of 7.1 mM and 31.1 mM, respectively (Figure 4b). Although at saturating concentrations L-threonine and L-cysteine induced comparable stabilisation of the protein, as evidenced by similar increases in T_m_ (to roughly 55 °C upon incubation with 50 mM of L-threonine, compared to around 54 °C with 125 mM *L*-cysteine), full stabilisation required higher concentrations of L-cysteine. In contrast, L-serine, L-alanine, and L-valine showed minimal and poorly defined binding, with *K*_D_ values of 12.5 ± 12.3 mM, 18 ± 19 mM, and 14.8 ±13.7 mM, respectively (Figure 4b, Table S2). Nevertheless, a gradual increase in T_m_ was observed with increasing ligand concentrations across all tested amino acids, suggesting that even weak binders may contribute to marginal protein stabilisation, likely through a combination of specific low-affinity interactions and non-specific effects. These findings collectively indicate that while YRDC shows some promiscuity in amino acid binding, L-threonine remains functionally distinct in its ability to induce conformational stabilisation. We also considered whether D-amino acids could bind to human YRDC, and tested D-valine, D-alanine, and D-threonine (Supplementary Figure 7). A trend was observed for D-valine, D-alanine yielding *K*_D_ values >25 mM, and although stabilization occurred with D-threonine, determination of an approximate *K*_D_ value was not possible, as it has a *K*_D_ values >100 mM. This shows that YRDC favours L-amino acid binding.

**Figure 4.**
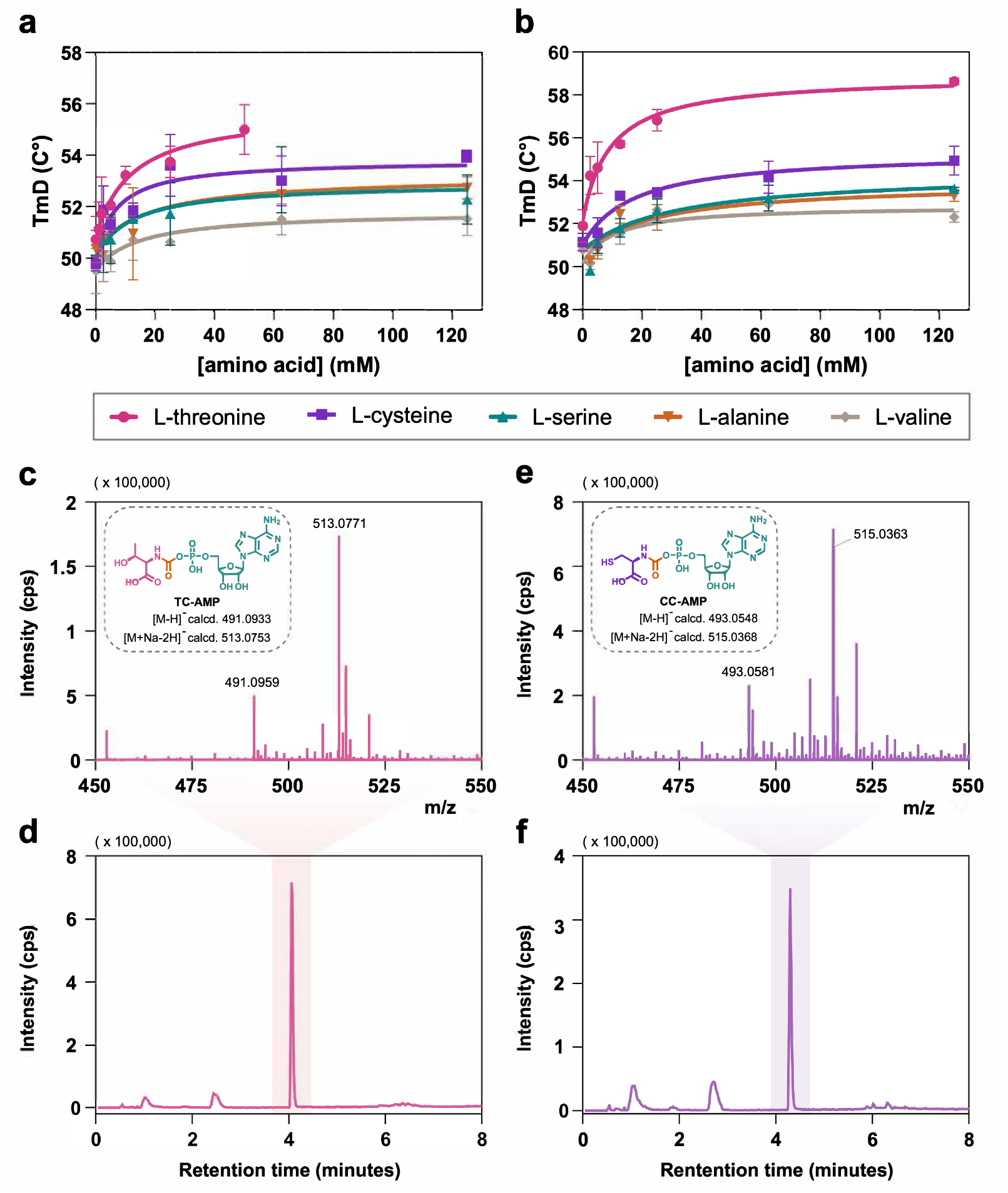
Amino-acid binding to YRDC and LC–ESI–MS detection of threonyl- and cysteinyl-carbamoyl adenylates. **a, b** Binding of human YRDC_WT_ to L-cysteine, L-serine, L-alanine, and L-valine was assessed by DSF across increasing concentrations of each amino acid and compared with binding to the canonical ligand L-threonine. Experiments were performed in two conditions: without ATP (a) and with 500 µM ATP (b). Each condition was measured in three independent experiments (n = 3), and the derivative melting temperature (TₘD) is plotted as mean ± s.d. Concentration–TₘD curves were fit using the equation specified in the Methods to obtain K_D_ values for each amino acid under both ATP conditions. **c, d** High-resolution LC–ESI–MS (negative ion mode) of the YRDC_WT_ reaction with L-threonine, ATP, and bicarbonate at 37 °C, acquired with target enhancement at *m/z* 513. Peaks consistent with threonylcarbamoyl-adenylate (TC-AMP) are observed as the monoanion [M–H]⁻ at *m/z* 491 and the monosodium dianion [M+Na–2H]⁻ at *m/z* 513. The experiment was conducted three times independently, yielding similar spectra in all replicates. **e, f** Under analogous conditions, YRDC_WT_ reacts with L-cysteine to generate a cysteine-derived carbamoyl adenylate detectable by LC–ESI–MS in negative ion mode with target enhancement at *m/z* 515. Signals assigned to the cysteine adduct are observed as the monoanion [M–H]⁻ at *m/z* 493 and the monosodium dianion [M+Na–2H]⁻ at *m/z* 515. This experiment was performed twice independently (n = 2) and produced consistent results. Abbreviations: aa, amino acid; TC-AMP, threonylcarbamoyl-AMP; CC-AMP, cysteinylcarbamoyl-AMP. For c-f CPS stands for counts per second. Raw data for all is shown on Supplementary Figure 8.

Given the earlier observation that ATP enhances the stabilisation of YRDC after L-threonine binding, we next examined how ATP influences the interaction between YRDC and these amino acids. Under these conditions, L-threonine exhibited the strongest stabilising effect on the YRDC – ATP complex, increasing its T_m_ to approximately 58.6 °C at a concentration of 125 mM. This stabilisation was greater than that conferred by the next most effective ligand, L-cysteine, which raised the T_m_ to approximately 55 °C under identical conditions. Interestingly, the presence of ATP also enhanced the stabilising effects of the other tested amino acids (L-serine, L-alanine, and L-valine) on YRDC, as reflected by increased T_m_ values with rising ligand concentrations. However, the *K*_D_ values for L-cysteine, L-serine, and L-alanine slightly increased in the presence of ATP, suggesting reduced binding affinities under these conditions.

To evaluate whether human YRDC can catalyse reactions involving alternative substrates, the activity with different amino acids was evaluated in the presence of ATP and bicarbonate by LC–ESI–MS. The results revealed that YRDC can catalyze the formation of carbamoyl–AMP products from all tested amino acids. As with the reaction involving L-threonine, negatively charged ions [M–H]⁻ were detected, with measured masses matching the theoretical molecular weights of the expected final products corresponding to each amino acid (Supplementary Figure 8). In all cases, the monosodium dianion species [M+Na–2H]⁻ produced a stronger signal than the [M–H]⁻ form, suggesting that the carbamoylated products predominantly exist in the monosodium dianionic state under the experimental conditions. For instance, in the reaction using *L*-cysteine, YRDC catalysed the formation of cysteinylcarbamoyl-AMP, with [M–H]⁻ detected at *m/z* = 493.0548 (theoretical exact mass = 493.05), and [M+Na–2H]⁻ at *m/z* = 515.0368. In addition, the [M–H]⁻ signal of the reaction by-product AMP (*m/z* = 346.0558) was consistently observed across all reactions, further confirming enzymatic activity. These findings demonstrate that YRDC can utilize amino acids beyond L-threonine as substrates when combined with ATP and bicarbonate. However, the consistently lower signal intensities of these alternative products relative to TC-AMP suggest that L-threonine remains the preferred and most efficient substrate for human YRDC.

#### YRDC can bind free L-threonine, with lack of evidence for threonine–carbamate adduct binding

A threonine–carbamate adduct can be generated non-enzymatically from L-threonine and bicarbonate, especially at higher pH values.^39^ Using [^15^N] L-threonine and [^13^C] NaHCO₃ to acquire ^1^H, ^13^C, ^1^H–^13^C HMBC, and ^1^H–^15^N HMBC NMR spectra enabled the detection of this adduct regardless of the presence of YRDC (Supplementary Figure 9). Integration of ^1^H peaks relative to threonine place threonine-carbamate abundance at approximately 9 % (or 4.5 mM, considering 50 mM [^15^N] L-threonine as starting material). No chemical shift perturbations were observed by the addition of 50 µM YRDC and ATP. We cannot discard that the concentration of threonine-carbamate-YRDC complex is too small to be detected under these conditions, but increasing the time of incubation did not result in changes in the amount of threonine– carbamate, in the presence or absence of enzyme.

The stability of the threonine–carbamate adduct is limited, and we could not under the current experimental conditions increase its concentration in solution. We therefore employed saturation transfer difference (STD) NMR to confirm that YRDC recognizes L-threonine directly (Figure 5a). The STD spectra clearly shows a week effect for a L-Thr β-H (4.27 ppm) resonance, indicative of binding to YRDC under these conditions. Figure 5b depicts two possible mechanisms for this reaction, one that does not include the formation of a threonine-carbamate intermediate (Mechanism 1), and one that does follow this path (Mechanism 2). Our data shows that while both mechanisms are possible *on enzyme*, YRDC can directly bind threonine, which lowers the *K*_D_ for ATP, and in turn also increases the affinity for bicarbonate (Supplementary Table 2). These observations support a mechanism in which YRDC forms a quaternary complex with L-threonine, ATP, and bicarbonate to form TC-AMP, rather than requiring pre-formed threonine– carbamate.

**Figure 5.**
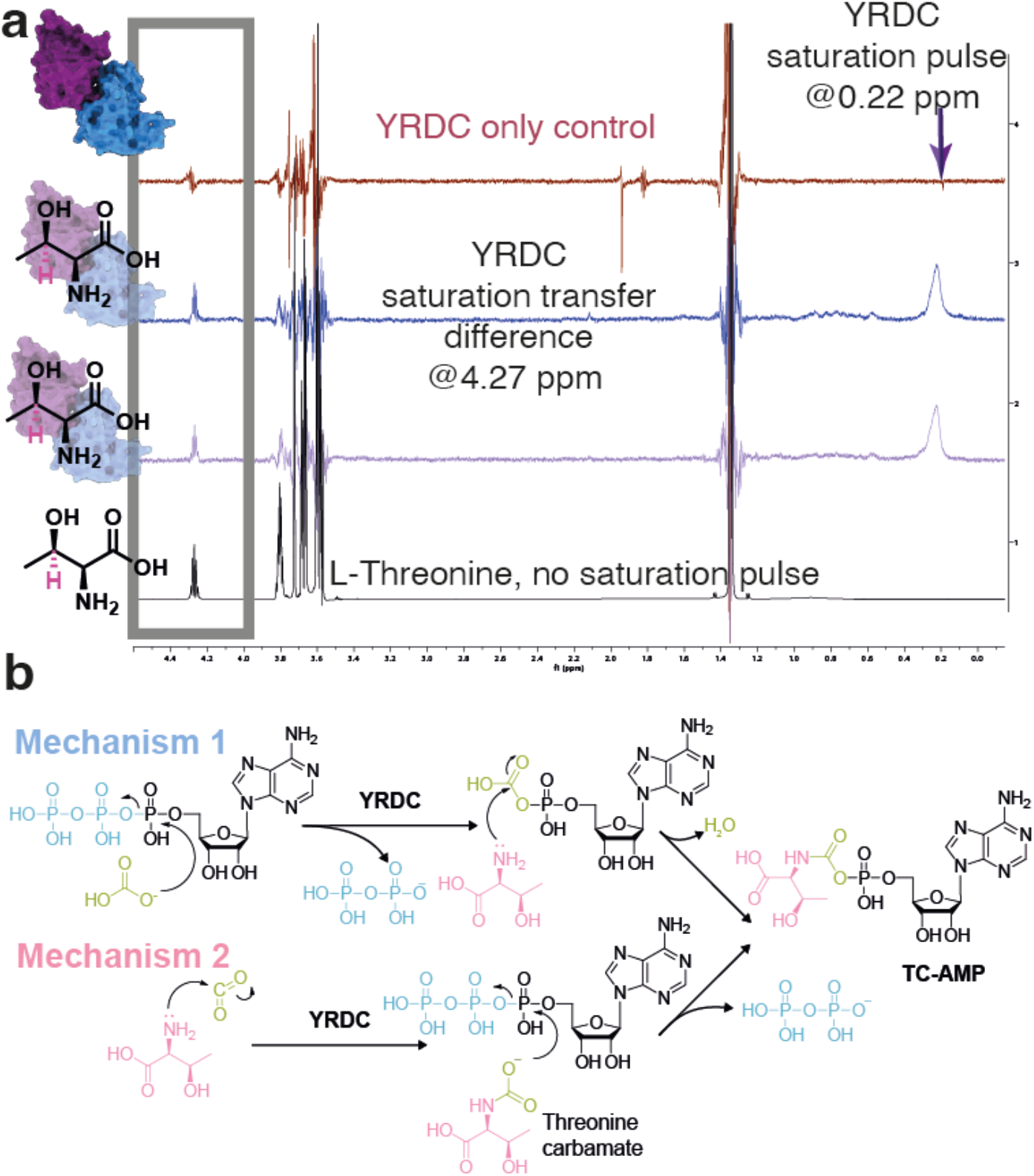
Saturation transfer difference (STD) NMR shows that YRDC binds L-threonine directly, and proposed routes to TC-AMP formation. **Top:** comparison of spectra shown as Black trace: reference off-resonance spectrum for the sample containing both threonine and YRDC with saturation pulse at 40 ppm; purple trace: saturation transfer difference spectrum, 37 °C, saturation pulse at 0.22 ppm, saturation time 0.5 s, recycle delay D1 3s; blue trace: saturation transfer difference spectrum saturation time 1s, further increasing of saturation time does not lead to any substantial improvement of STD effect; maroon trace: control STD experiment on sample containing threonine in protein buffer (flow through from buffer exchange) but no YRDC. Threonine β-H (4.27 ppm – shown in pink and with a grey box highlighting the peak) shows only weak antiphase difference spectrum artefact while the STD experiments carried on the sample containing enzyme (purple and blue curves) show the same peak as inphase multiple that indicates a weak STD effect. **Bottom**: two proposed mechanisms for the YRDC-catalyzed reaction. A quaternary complex must be formed for the reaction to start, but whether this proceeds via direct attack into the alpha-phosphate of ATP to release pyrophosphate (Mechanism 1), or via the formation of a threonine carbamate (Mechanism 2) intermediate remains to be determined.

#### GAMOS-related mutations impact YRDC function, oligomeric state and stability

To gain insight into how GAMOS-associated mutations affect the structure of YRDC, we generated and characterised three previously reported single-point variants: YRDC_A41V_, YRDC_I178T_, and YRDC_ΔL222_ (Figure 6). Of the three mutants, the YRDC_A41V_ variant exhibited the most stable conformation, with a T_m_ of 48.3 °C as determined by DSF, closely matching the value obtained from CD spectroscopy at 47.0°C. In contrast, the YRDC_I178T_ and YRDC_ΔL222_ mutants displayed significantly less stable secondary structures, with T_m_ of 35.9 °C and 44.5 °C, respectively, and CD spectra indicating partial unfolding or misfolding. Both mutants also showed elevated baseline fluorescence at ambient temperature (20 °C), consistent with increased exposure of hydrophobic regions due to structural destabilisation (Supplementary Figure 2). In terms of secondary structure content (Figure 6b), predictions for these three mutants, compared with WT, suggested largely disordered conformations. To evaluate this, CD spectral data were analysed using the SESCA algorithm with the DS-dT basis set^38^. All three GAMOS-associated mutants exhibited significantly higher coil content than WT, which had a total disordered and turn content of approximately 56.9%. Specifically, the coil fractions of YRDC_A41V_, YRDC_I178T_, and YRDC_ΔL222_ in native buffer conditions were estimated at 78.99 ± 10.86 %, 75.91 ± 9.53 %, and 77.95 ± 11.36 %, respectively. These differences in folding between wild type and GAMOS mutants were in agreement with ^1^H NMR of each protein variant (Supplementary Figure 10).

**Figure 6.**
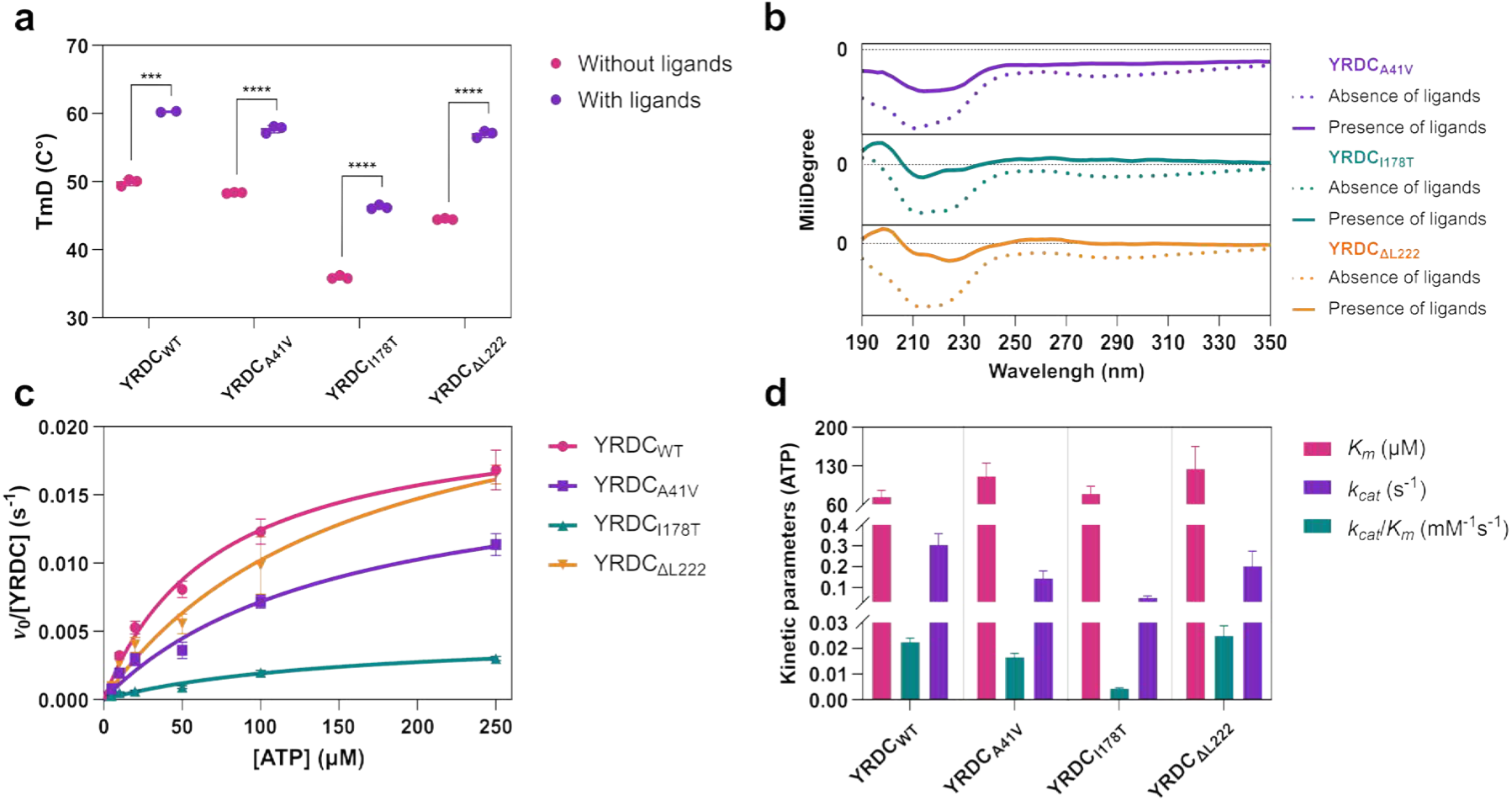
Thermal stability, secondary structure, and catalytic kinetics of YRDC_WT_ and GAMOS-related mutants. **a** Melting temperature values (Tₘ) for YRDC_WT_ and GAMOS-related mutants measured by DSF under two conditions: without ligands (green) and with ligands (pink; 500 µM ATP and 50 mM L-threonine). Each condition was measured in triplicate, and values are shown as mean ± s.d. (n = 3). Unpaired two-sided t-tests with Holm–Šídák correction were used to compare the Tₘ between the two conditions for each enzyme. The significance threshold was P < 0.05; annotations: P < 0.001 (***), P < 0.0001 (****), and not significant (P > 0.05; ns). **b** CD spectra of the three GAMOS-related mutants recorded in the absence (dashed lines) and presence (solid lines) of ligands (500 µM ATP and 50 mM L-threonine) at 25 °C. Data are plotted as millidegrees versus wavelength over 190–350 nm. **c** Michaelis–Menten plots comparing the catalytic behaviour of GAMOS-related mutants with YRDC_WT_. Global fits to the Michaelis–Menten model were performed across three independent replicates, and data are displayed as mean ± s.d. (n = 3) for each substrate concentration. **d** Summary of kinetic parameters for YRDC_WT_ and the three GAMOS-related mutants obtained by varying ATP from 10 µM to 250 µM. The Michaelis– Menten constant (*K*_m_), turnover number (*k_cat_*), and catalytic efficiency (*k_cat_*/*K*_m_) for each protein are highlighted in purple, green, and red, respectively. Plots report mean ± s.d. (n = 3). Abbreviations: T_m_, derivative melting temperatures.

Furthermore, a notable difference from the WT or substrate-binding site mutants was observed: none of the GAMOS mutants showed any further stabilisation of secondary structure upon ligand binding. Their β-sheet and coil content remained unchanged in both ligand-free and ligand-bound conditions (Supplementary table 2). This suggests that these GAMOS-associated variants exist predominantly in a disordered state, with coil content approaching 80%. Furthermore, the extremely low α-helix content observed across all three mutants, along with the high uncertainty in these values (± SD), indicates that stable α-helical structures are virtually absent. Regarding the β-sheet fraction, values fall around 20% in the three mutants, suggesting the presence of this conformation, but it is not clearly predominant and predicted with low confidence. Moreover, a dramatic drop in the CD spectral scaling factor compared with WT, which exhibited a scaling factor near 1, suggests that the signal intensity was diminished, likely due to unfolding, aggregation, or structural heterogeneity. Taken together, these results indicate that the mutants associated with GAMOS exist in a largely disordered conformation, consistent with the thermal instability and partial misfolding observed in the DSF experiments. These findings demonstrate that GAMOS-associated mutations profoundly alter the structural conformation of YRDC, leading to a predominantly disordered and thermally unstable state that does not respond to ligand-induced stabilisation.

To evaluate the functional consequences of GAMOS-associated mutations, the kinetic parameters of the three previously described variants, including YRDC_A41V_, YRDC_I178T_, and YRDC_ΔL222_, were determined using the LC–MS-based method described above, with varying ATP. All three variants showed higher *K*_m_ values clustered near 100 µM (YRDC_A41V_ 110.8 ± 24.2 µM; YRDC_I178T_ 79.3 ± 13.7 µM; YRDC_ΔL222_ 124.6 ± 40.5 µM), higher than wild type and consistent with weaker ATP engagement (raw data on Supplementary Figure 6). With respect to catalytic turnover, the YRDC_A41V_ point mutation (0.016 ± 0.002 s^−1^) did not show a statistically significant difference compared to the WT enzyme, suggesting that the mutation does not impair catalytic rate. Similarly, the YRDC_ΔL222_ mutant exhibited only a slightly elevated turnover rate, with a *k_cat_* value of 0.025 ± 0.004 s⁻¹. In contrast, the YRDC_I178T_ mutation resulted in a marked decrease in catalytic efficiency, with a *k_cat_* value approximately six-fold lower than that of WT, indicating a substantial reduction in enzymatic activity while *K*_m_ stayed close to the wild-type range, pointing to a primary defect in the catalytic step rather than in ATP recognition. The overall catalytic efficiencies for the YRDC_A41V_, YRDC_I178T_, and YRDC_ΔL222_ variants were 0.10 ± 0.04 mM^−1^s^−1^, 0.05 ± 0.01 mM^−1^s^−1^, and 0.20 ± 0.07 mM^−1^s^−1^, respectively. Taken together, these results indicate that two of the three GAMOS-associated mutations (YRDC_A41V_ and YRDC_ΔL222_) do not significantly affect catalytic turnover but do impair substrate binding affinity. In contrast, YRDC_I178T_ slows the turnover rate despite near–wild-type ATP affinity.

#### YRDC is dimeric and GAMOS variants disrupt dimerization

To date, the three-dimensional structure of human YRDC has not been experimentally determined, and the structural basis underlying its enzymatic activity remains poorly understood. Structural modelling of YRDC using AlphaFold suggests that the GAMOS-associated mutations are located at the periphery of the protein, distant from the predicted substrate-binding site. Consistent with this spatial separation, two of the three mutants (YRDC_A41V_ and YRDC_ΔL222_) exhibit catalytic turnover rates (*k_cat_*) comparable to the wild-type enzyme, indicating that their reduced activity is not due to impaired chemical catalysis per se. Instead, combined data from DSF and CD spectroscopy reveal that these distal mutations destabilise the secondary and tertiary structure of YRDC. This raises an interesting question: how can point mutations located far from the active site give rise to severe functional deficits associated with GAMOS? Given their peripheral positions, we hypothesised that YRDC functions as a dimer or higher-order oligomer, and that these mutations impair its ability to adopt or maintain the correct quaternary structure, thereby compromising enzymatic performance. To test this hypothesis, we analysed the oligomeric states of the wild-type protein and GAMOS-associated variants by native mass spectrometry (Figure 7a and b and Supplementary Figure 11), native PAGE (Figure 7c), mass photometry (Figure 7d-g and Supplementary Figure 12), crosslinking mass spectrometry using BS3 (bis(sulfosuccinimidyl)suberate, Figure 7h, Supplementary Figure 13), and size exclusion chromatography (Supplementary Figure 14).

**Figure 7.**
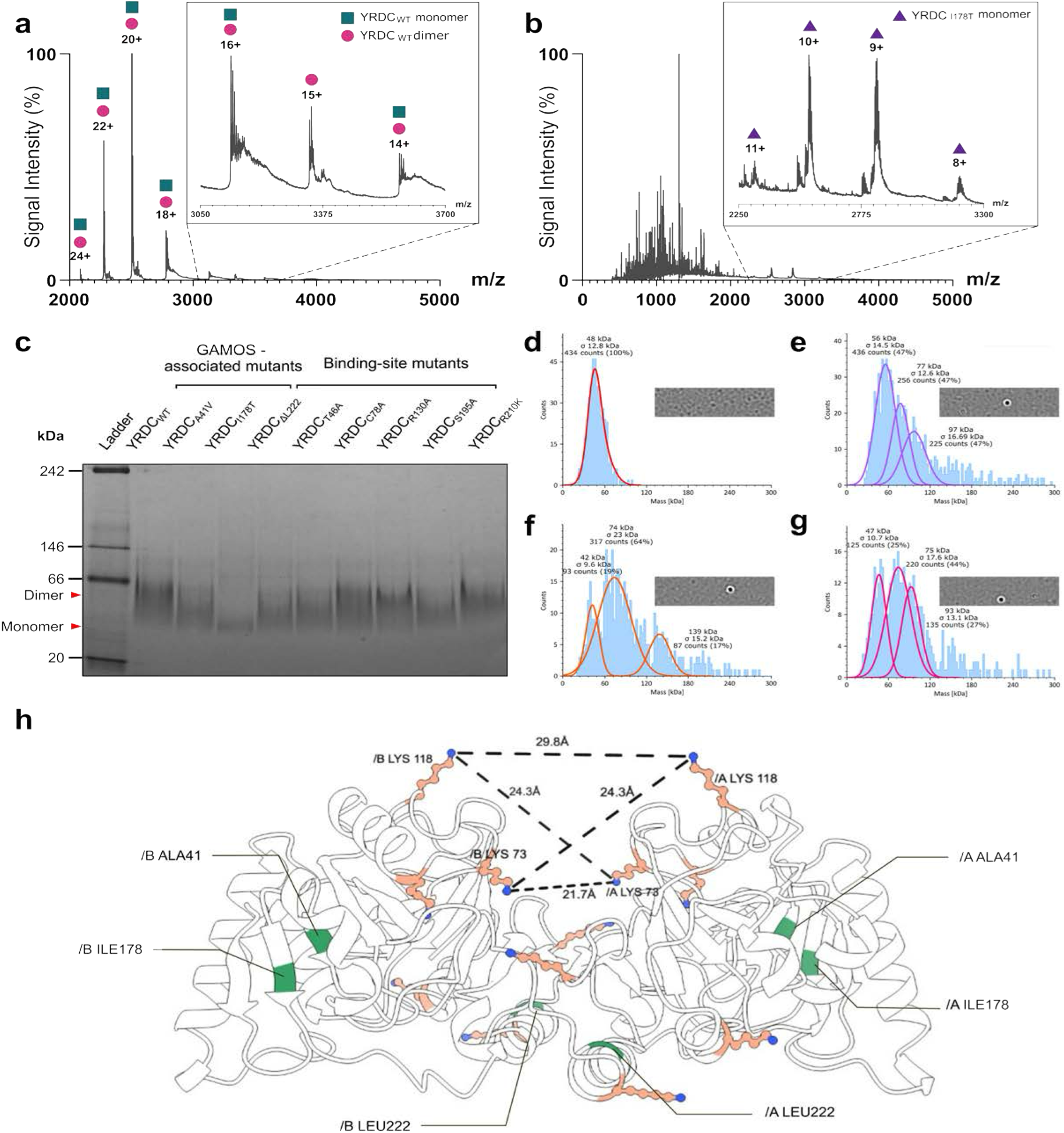
YRDC_WT_ is a dimer in solution. **a** Native mass spectrum of YRDC_WT_ acquired under non-denaturing conditions in 50 mM ammonium acetate (pH 6.8). Green dots mark the monomer charge-state series and orange dots mark the dimer series; numbers above peaks denote the assigned charge state (z⁺) of dimer. Insets display zoomed regions of representative dimer charge. **b** Native mass spectrum of the YRDC_I178T_ variant recorded under identical conditions. Blue dots indicate the monomer charge-state series; numbers denote the corresponding z⁺ values of its monomer. Insets show enlargements of individual charge states used for assignment. **c** Native PAGE analysis of YRDC_WT_ and all mutants examined in this study (binding-site substitutions and GAMOS-associated variants). Gel bands corresponding to monomeric and dimeric assemblies are indicated. **d–g** MP of YRDC_WT_ and its GAMOS-related mutants assemblies in solution. Top panels: representative video frames from 180 s recordings. Bottom panels: mass-distribution histograms obtained with DiscoverMP analysis (Methods). Under these conditions, YRDC_WT_ (**d**) shows a predominant species at 48.0 ± 12.8 kDa, consistent with a dimer, with no higher-order peak detected. By contrast, all three GAMOS-related mutants (YRDC_A41V_, **e**; YRDC_I178T_, **f**; YRDC_ΔL222_, **g**) display, in addition to a reduced dimer population, higher-order oligomers (including trimer and tetramer), as indicated by peaks at integer multiples of the monomer mass within the instrument’s mass tolerance. **h** AlphaFold-predicted dimer of YRDC illustrating the spatial distribution of GAMOS-associated substitutions across the two protomers. Residues in green are mutations associated with GAMOS, and lysine residues depicted in orange. Distances shown depict lysine residues that were crosslinked with BS3, indicating they are in proximity. Supplementary Figure 13 depicts crosslinking and distances for other AlphaFold predicted dimers.

On native page, YRDC_WT_ migrated primarily as a dimer, consistent with an apparent molecular weight of ∼50 kDa, as did most binding-site mutants under the same conditions. In contrast, two of the three GAMOS-associated variants – YRDC_A41V_ and YRDC_ΔL222_ – exhibited a mixture of monomeric and dimeric species, while the YRDC_I178T_ mutant was observed predominantly in the monomeric form (∼25 kDa). We confirmed these findings by native mass spectrometry (Figure 7a and 7b, Supplementary Figure 12). Under native conditions, YRDC_WT_ displayed a dominant high-*m/z* charge-state envelope consistent with a dimer. Deconvolution yielded a neutral mass approximately twice the monomer, around 50 kDa within TOF mass accuracy, alongside a weaker monomer series at lower *m/z*. The theoretical monomer mass, excluding an N-terminal 43-residue segment, was 25,040.9 Da. We observed a ∼76.0 Da increment between the paired monomer (and dimer) populations, most consistent with a single mixed disulfide covalent adduct with β-mercaptoethanol (BME, adduct with theoretical mass +74 of 25,116.9), or glycerol (adduct with theoretical mass +76 of 25,114.9) formed during purification. Native deconvolution returned two monomer populations of 25,040.3 ± 3.3 Da and 25,114.3 ± 6.1 Da, and two dimer populations of 50,082.8 ± 11.6 Da and 50,153.1 ± 12.5 Da (expected as 50081.8 or 50,155.8 for an adduct +74). In contrast, denaturing conditions collapsed the spectrum to a single monomer series whose deconvolved mass matched the calculated protein mass, confirming the dimer assignment in the native spectra. GAMOS-related mutants did not reproduce the WT dimer distribution. YRDC_A41V_ and YRDC_ΔL222_ showed only small dimer signals; most variants were predominantly monomeric and, in some cases, produced additional envelopes that deconvolved to trimeric or higher-order assemblies. YRDC_I178T_ was detected exclusively as a monomer. In native spectra for several mutants, we also observed an additional mass of approximately 475.0 Da together with extensive adducting; these features were absent under denaturing conditions and are consistent with a non-covalent ligand or buffer-associated adduct that remains to be identified. The loss of the WT dimer envelope and the appearance of monomer and alternative oligomers indicate that these mutations destabilise the native dimer interface and redirect self-association toward less stable architectures. Deconvolved intensities showed that the dimer accounts for the majority of the WT signal, whereas mutants shift strongly toward monomer with minor higher-order species. Together, these data support a model in which YRDC forms a dimer as its predominant native assembly and GAMOS-related mutations disrupt this quaternary structure.

Lower molecular weight species including monomers cannot be reliably analysed by single-molecule MP. Instead, we used this approach to assess higher-order assemblies and to validate quaternary states. In reaction buffer at pH 8.0, YRDC_WT_ yielded a single population at 48.0 ± 12.8 kDa, consistent with a dimer, and no higher-order peak was detected. In contrast, YRDC_A41V_ and YRDC_ΔL222_ were predominantly higher-order, with trimer fractions of 28% and 44%, tetramer fractions of 24% and 27%, respectively, and additional low-abundance species above the tetramer range in both cases. The YRDC_I178T_ variant exhibited the lowest dimer fraction (19%) among the mutants. Monomeric species were not observed by MP due to its lower effective detection limit of ∼30 kDa; inference of monomer enrichment therefore relies on the native MS data, which show a dominant WT dimer envelope that collapses to monomer under denaturing conditions. We also performed MP in the presence of ligands (ATP 500 µM and L-threonine 50 mM, Supplementary Figure 13) and observed no appreciable change in the mass distributions, indicating that ligand binding does not measurably alter the oligomeric state under these conditions. Taken together with native MS, these measurements indicate that YRDC forms a dimer as its predominant assembly in solution, whereas GAMOS-related mutations destabilise the dimer and redistribute the population toward non-dimeric oligomers.

To define residues in close proximity due to dimerization, we carried out a crosslinking experiment with BS3. This modifier can crosslink lysine residues within distances between 11.4 Å – 30 Å^40^. Following crosslinking with BS3, samples are run in a denaturing SDS-PAGE, enabling the separation of crosslinked and non-crosslinked species. We observed YRDC dimers crosslinking between Lys118 and Lys73 exclusively on gel bands at ∼50 kDa (Figure 7h, Supplementary Figure 14), demonstrating they are in close proximity and providing experimental evidence for the dimer interface in YRDC_WT_.

## Discussion

Our work reveals key aspects of YRDC catalysis and stability, and how these are linked to mutations that cause disease. Threonine is essential in humans, but not among the most abundant free-amino-acid pools in many mammalian cells^41^, and its availability depends on diet and membrane transport. Shedding light into YRDC’s preference towards threonine, we determined that it strongly stabilizes the human enzyme leading to an increase in the affinity for co-substrates ATP and bicarbonate. DSF shows that several amino acids could bind weakly at low millimolar concentrations, but L-threonine consistently yields the largest thermal shift and leads to a larger shift in the presence of ATP. L-threonine organizes the enzyme into a conformation that not only binds ATP more tightly but also engages bicarbonate more effectively. The order of ligand binding follows directly from those observations as threonine tightens ATP binding by more than one order of magnitude, and bicarbonate binding becomes measurable only when L-threonine is in place. Our data support an ordered mechanism where L-threonine binds first, followed by ATP and bicarbonate.

Two different mechanistic proposals have been put forward for TC-AMP formation. One proposes that threonine reacts with CO₂ or bicarbonate in solution to form a threonine carbamate adduct that is then adenylated by the enzyme^32,42^. Another suggests that bicarbonate is first activated in an ATP-dependent way within the multi-protein bacterial system and that the reaction proceeds through additional activated intermediates, with AMP and ADP seen in bulk assays that track the full pathway^26^. These proposals differ on where the carbamate resides and how many nucleotide activation steps precede the formation of TC-AMP^26^.

Our data for the human mitochondrial YRDC strongly support that carbamoylation occurs on enzyme. Threonine binds directly to YRDC in the absence of bicarbonate and ATP. Bicarbonate binding occurs only after threonine is in place, similarly to ATP binding to the enzyme, which is extremely weak but tightens once threonine binds. ^31^P NMR shows no ADP signal during reaction and only a very low level of AMP at the earliest time points. If a carboxyl-AMP intermediate were obligatorily formed in solution and then aminolyzed before TC-AMP, substantial AMP would be expected prior to TC-AMP formation. The absence of ADP also argues against an additional ATP-dependent activation prior to TC-AMP formation. Together with the NMR STD data, this pattern supports a mechanism in which all three substrates are bound to the enzyme to form a quaternary complex.

This mechanism clarifies previous conflicting reports. The “two-ATP” hypothesis largely came from full-pathway experiments that tracked downstream products by TLC and determined both AMP and ADP^26^. Such readouts integrate multiple steps and side reactions and can conflate the YRDC step with the KEOPS/TsaBD transfer chemistry and with non-enzymatic conversions. Carbamate formation has a strong dependence on bulk pH, becoming negligible as the pH approaches 7.0^39^.

In the structure of *Pa*-Sua5 bound to TC-AMP, threonine participates in a network of hydrogen bonds to residues on the catalytic pocket.^25^ Here we mutated the equivalent residues predicted by structural alignment to interact with TC-AMP – YRDC_T46A_, YRDC_C78A_, YRDC_R130A_, YRDC_S195A_, and YRDC_R210K_. Mutants YRDC_T46A_ and YRDC_R130A_ are poorly folded by DSF and CD and fail to engage ATP in the absence of threonine. They participate on the initial threonine recognition step which is needed for reaction to occur. Other substitutions such as YRDC_S195A_ and YRDC_R210K_ still permit threonine-enabled ATP binding, though with decreased affinity.

Because ATP binds YRDC in the micromolar range, whereas L-threonine and bicarbonate require millimolar concentrations, flux through the pathway is controlled by the availability of bicarbonate and threonine rather than ATP, a conclusion broadly aligned with previous work on the human mitochondrial system demonstrating CO₂/bicarbonate sensitivity of t⁶A formation.^32^ Threonine must be carbamoylated before adenylation is productive, and bicarbonate is often the driver of flux in cells. Lin et al. reported different absolute values for kinetic parameters as their assay depended on the conversion of TC-AMP to a secondary product under mildly alkaline conditions. In comparison, while apparent *K*_m_ values differ, experimental data agree on the qualitative affinity for substrates^43^. Direct measurements of human mitochondrial threonine are limited, but rat liver mitochondria contain threonine at approximately 1–2 mM, higher than serum levels near 0.5 mM^44^. *K*_m_ and *K*_d_ values in the millimolar range are high considering this physiological range, and we cannot discard that the cellular environment and concentration ratios of threonine, other proteins and small molecules might shift these values.

Prior work reported that YRDC_WT_ and the YRDC_A41V_ and YRDC_ΔL222_ mutants are “well folded” based on ^1^H NMR spectra showing amide dispersion and upfield methyl resonances, but we cannot discard that a native-like global fold was still absent^12^. We carried out this experiment with YRDC_WT_, YRDC_A41V_, YRDC_I178T_ and YRDC_ΔL222_ and our findings are in agreement for YRDC_WT_, YRDC_A41V_ and YRDC_ΔL222_, but show that YRDC_I178T_ depicts fewer peaks between 0 and 0.5 ppm, as well as sharper peaks in the 6.5-8.5 ppm region, both indicative of a less well folded protein (Supplementary Figure 10).

Under our conditions, differences in buffer, ligands, temperature, protein concentration, and presence of purification tags affect stability. Far-UV CD for the mutants indicates reduced or redistributed secondary structure relative to WT, consistent with global destabilization or localized disorder rather than complete unfolding. In addition, adding ATP or L-threonine raises the apparent melting temperature but does not increase α-helix or β-sheet content, which means ligands stabilize non-native or heterogeneous states rather than restoring the native fold. This explains how mutations far from the active site still affect activity, potentially shifting the conformational energy landscape and reducing the population of competent states instead of directly impacting the chemical steps in the reaction.

Our data supports that YRDC_WT_ forms a dimer, which is essential for stability and function. We propose that GAMOS-linked distal mutations disrupt dimer interface in distinct ways: YRDC_I178T_ collapses almost entirely to monomer, while YRDC_A41V_ and YRDC_ΔL222_ weaken dimerization and promote aberrant oligomers including trimers and tetramers. More work is required to determine how these mutations lead to these effects, but sequence alignment as well as predicted structural alignment between human YRDC and homologues from mouse, plants, and other highly divergent organisms reveal high conservation on positions equivalent to YRDC_A41V_ and YRDC_ΔL222_ (Supplementary Figure 15). More variation occurs on amino acids at the YRDC_I178_ position, which is replaced by leucine, valine, phenylalanine or tyrosine, all hydrophobic and substantially different from threonine.

For human YRDC, structural defects correlate with functional changes: YRDC_A41V_ and YRDC_ΔL222_ retain near-wild-type turnover but show reduced ATP affinity, whereas YRDC_I178T_ exhibits significant catalytic loss, likely due to failure to maintain the active dimer. Prior Ppi-release assays confirm decreased TC-AMP synthesis for these variants^12^. Dimerization emerges as a critical control point in which GAMOS-related mutants are deficient. Our work complements clinical observations on benefits of dietary therapy with threonine starvation in glioblastoma and provides a starting point to design molecules that modulate YRDC stability and, consequently, activity.

## Methods

### 1. Materials

Common reagents, buffers, and salts used in this research were obtained from Merck and Fisher Scientific and were used without further purification or modification. LC–MS grade ultrapure water and formic acid (> 99%) along with HPLC grade methanol (> 99%) were purchased from Fisher Scientific. PureYield™ Plasmid Miniprep System from Promega was utilised in this study for plasmid purification. All DNA oligonucleotides were manufactured by Integrated DNA Technologies (IDT). Data were analysed using GraphPad Prism version 10.4.1 for Windows (GraphPad Software, United States). All chemical structures and exact mass calculations of the compounds and their corresponding ions were generated using ChemDraw 22 (PerkinElmer, United States). Additionally, Liquid Chromatography-Mass Spectrometry (LC–MS) data were processed using MassLynx V4.2 for Windows (Waters, Massachusetts, USA), while protein stability data were investigated using Protein Thermal Shift™ software (Thermo Fisher Scientific, Massachusetts, United States).

### 2. Cloning, mutagenesis and expression

The DNA sequence encoding the human YRDC enzyme (Uniprot: Q86U90) was truncated by forty-three amino acids at the N-terminal and codon-optimised by Codon Optimization Tool (IDT) for expression in *E. coli*. The construct was designed for insertion into a pET-28a(+) expression plasmid vector, containing an N-terminal His₆-tag with a tobacco etch virus (TEV) protease cleavage site, manufactured by GenScript Biotech. For human YRDC mutants, pET-28a(+)-YRDC(wt)^6H^ was utilised as a template for site-directed mutagenesis using Q5® High-Fidelity DNA Polymerase and the protocol designed by NEBaseChanger (New England Biolabs). PCR products were digested by *Dpn*I, then treated with T4 ligase, and polynucleotide kinase (NEB) for 1 hour at room temperature.

All recombinant plasmids were transformed into *E. coli* DH5α cells (NEB) for amplification and their sequence were verified by Sanger sequencing (Eurofins). Verified constructs were transformed into E. coli BL21(DE3) competent cells (NEB) for protein expression. Transformed cells were grown in lysogeny broth media (LB) supplemented with 100 μg/mL kanamycin at 37 °C with shaking at 180 rpm until reaching an OD_600_ of 0.6 – 0.8. Protein expression was induced with 0.5 mM isopropyl β-d-1-thiogalactopyranoside (IPTG), followed by incubation at 16 °C overnight, while shaking at 180 rpm. Subsequently, cells were harvested by centrifugation (12,000 *g*, 30 min, 4 °C) and stored at −80 °C for further processing.

### 3. Purification of YRDC and its mutants

Bacterial cell pellets were resuspended in lysis buffer containing 100 mM Tris-HCl (pH 8.0), 100 mM KCl, 2 mM β-mercaptoethanol (BME), 0.1 mM EDTA, 10% (v/v) glycerol. To enhance cell lysis, 30 mg of lysozyme and 3 mg of DNase per liter cell culture were added, followed by incubation at 4 °C for 30 minutes. Cells were then lysed using a cell disruptor (Constant Systems) at 30,000 psi at 5 °C, the lysate was separated by centrifugation at 56,000 x *g* for 30 min at 4 °C. The supernatant was filtered with a 0.8 μm membrane filter before loading onto a 5 mL HisTrap™ FF nickel affinity column (Cytiva) which was pre-equilibrated with lysis buffer. To remove unbound proteins the column was washed with 10 column volume of 100% buffer A (100 mM Tris-HCl pH 8.0, 100 mM KCl, 2 mM BME, 0.1 mM EDTA, 10% (v/v) glycerol, 30 mM imidazole), followed by elution of YRDC using buffer B (buffer A with 200 mM imidazole). The eluted protein fractions were then pooled and dialyzed against buffer C (100 mM Tris-HCl pH 8.0, 100 mM KCl, 2 mM BME, and 10% (v/v) glycerol) at 4 °C overnight, with the addition of TEV protease (2 mg/mL at a 1:25 ratio) to remove the N-terminal His₆ tag. Following TEV cleavage, the dialysed sample was loaded onto a second HisTrap™ FF nickel column equilibrated with buffer C. The flowthrough fractions containing cleaved YRDC were collected and analysed by SDS-PAGE. The cleaved protein was further purified by size-exclusion chromatography using a HiLoad™ 26/600 Superdex™ 200 pg column (Cytiva, 28989336), pre-equilibrated with SEC buffer (100 mM Tris-HCl pH 8.0, 100 mM KCl, 10% (v/v) glycerol, and 2 mM BME). The final protein was concentrated to 8-12 mg/mL, using the extinction coefficient at 280 nm (ExPASy ProtParam) to determine concentration. Purified YRDC was flash-frozen in liquid nitrogen and stored at −80 °C for further studies. The molecular mass of the purified protein was confirmed by liquid chromatography electrospray ionization mass spectrometry (LC–ESI–MS) at the Proteomics and Mass Spectrometry Facility (University of St Andrews). All the mutants of YRDC, including YRDC_A41V_, YRDC_T46A_, YRDCC_78A_, YRDCR_130A_, YRDC_I178T_, YRDC_S195A_, YRDC_R210K_, YRDC _ΔL222_, were expressed and purified following the same described purification protocol as the wild-type protein.

### 4. SYPRO orange DSF assay

Reactions were prepared in a MicroAmp™ 96-well optical reaction plate (Fisher Scientific) with a final volume of 20 μL per well, in assay buffer (50 mM Tricine pH 8.5, 10 mM MgCl₂) with 8 μM YRDC, and 5X SYPRO Orange dye (Invitrogen). The concentrations of substrates, including ATP, L-threonine, and NaHCO_3_, were varied based on the reaction type. In most cases wild-type YRDC and its mutants were assessed under various conditions, including enzyme alone, enzyme with ATP concentrations ranging from 10 µM to 500 µM, enzyme with L-threonine concentrations between 0.5 mM and 50 mM, either in the presence or absence of 500 µM ATP, and enzyme with NaHCO_3_ concentrations from 10 mM to 500 mM in the presence of both 500 µM ATP and 25 mM L-threonine. Fluorescence emission was recorded using Applied Biosystems® QuantStudio 1 Real-Time PCR system (λ_ex_ = 520 nm, λ_em_ = 558 nm). Temperature increases were performed over the range of 25–95 °C, with increments of 1 °C/min, while changes in fluorescence intensity were continuously measured.

For binding affinity comparisons, wild-type YRDC and amino acids, including L-threonine, L-cysteine, L-serine, L-alanine, and L-valine, the assay was conducted under the conditions described above, either with or without 500 μM ATP. The amino acid substrates were tested at concentrations ranging from 1 mM to 125 mM. For D-threonine, D-alanine, and D-valine, the assay was conducted under the conditions described above, with 500 μM ATP, each amino acid was tested at concentrations ranging from 1 mM to 125 mM.

Each experiment was conducted in triplicate, and the data are presented as the mean ± standard deviation. Control reactions, prepared without the enzyme, were used to account for background fluorescence and were subtracted from experimental measurements. Melting curves were fitted using a Boltzmann sigmoidal model, and thermal stability (Tₘ) values were determined from the inflection point of the curve. The dissociation constant (K_D_) of the enzyme with each substrate is calculated based on the DSF single binding equation^45^:

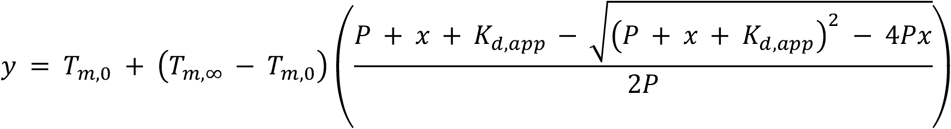

Where *T_m_*_,0_ and *T_m_*_,∞_refer to the melting temperature (*T_m_*) in °C at zero and infinite ligand concentrations, respectively, *P* represents the protein concentration and *K_d_*_,*app*_ is the apparent dissociation constant obtained from ligand-dependent thermal stabilisation.

### 5. LC–ESI–MS analysis of YRDC_WT_ with amino acid analogues

The catalytic activity of YRDC was assessed with alternative amino acid substrates, including L-threonine, L-serine, L-cysteine, L-alanine, and L-valine. Reactions were conducted in a total volume of 200 µL, comprising 5 µM YRDC, 500 µM ATP, 50 mM NaHCO_3_, and 50 mM of the respective amino acid in 50 mM Tricine (pH 8.0), 10 mM MgCl_2_. The mixtures were incubated at 37 °C for 3.5 hours. Following incubation, the enzyme was removed using a 3,000 MWCO centrifugal filter unit (Amicon® Ultra) by centrifugation at 4 °C for 10 minutes. This mild strategy to stop the reaction was used to ensure minimal degradation of TC-AMP and other reaction products, as well as for protein removal prior to LC-MS.

The monoisotopic mass of the reaction products, along with their multiply charged positive/negative ions, was determined using exact mass calculations in ChemDraw. Analyses were performed on an ACQUITY H-Class UPLC coupled to a Waters Xevo G2-XS Q-TOF mass spectrometer using an ACQUITY HSS column (2.1 × 100 mm, 1.8 µm). A flow rate of 300 µL/min at initial conditions of 99% H_2_O + 0.1% Formic Acid with 1% MeOH until 1 min, followed by a linear gradient step to 40% H_2_O + 0.1% Formic Acid with 60% MeOH until 4 min, followed by a step gradient to 1% H_2_O + 0.1% Formic Acid with 99% MeOH until 7 min and then re-equilibrated to 99% H_2_0 + 0.1% Formic Acid with 1% MeOH until 10 min. The column temperature was maintained at 40 °C throughout the run, samples were reconstituted in LC-MS grade water, and the injection volume was 1 µL.

The Q-TOF was operated in negative ESI mode with a capillary voltage of 2.5 kV. The ESI source and desolvation gas temperatures were set at 100 °C and 250 °C, respectively. The cone gas flow was set to 50 L hr^−1^, and the desolvation gas flow was set at 600 L hr^−1^. Acquisition was performed in TOF mode in the range of *m/z* 200 – 800 at 1 scan s^−1^ with target enhancement for each target analyte *m/z*. A lock spray signal was measured, and a mass correction was automatically applied by collecting every 10s, averaging 3 scans of 1s each using Leucine Enkephalin as a standard (*m/z* 554.2615).

### 6. CD spectroscopy

CD spectra were recorded using a MOS-500 CD spectrometer (BioLogic) in a 1 mm path length quartz cuvette (Hellma® Analytics) under continuous nitrogen flushing. CD spectra were collected over a temperature range of 20 °C to 95 °C, with measurements taken at 2.5 °C intervals following a 5-minute equilibration at each temperature. Measurements were conducted over a wavelength range of 190–350 nm with a spectral resolution of 0.2 nm and acquisition period of 0.5 seconds. Protein samples were prepared in 20 mM potassium phosphate buffer (pH 8.0) and 75 mM NaF, with final enzyme concentrations ranging from 12 to 20 μM. The experiments were performed either with or without ligands, including 50 mM L-threonine and 500 μM ATP. The molar ellipticity ([θ], deg cm² dmol⁻¹) was calculated according to the equation below. Data within the transition region were analysed by differentiation or polynomial fitting, and the apparent T_m_ values were defined as the midpoint of the transition, where 50% completion was observed. The results are presented as a replot of two independent measurements, and the acquired data were analysed using GraphPad Prism 10.4.1.

The secondary structure content of the WT protein and its mutants was estimated using the ChiraKit^38^ platform, which applies the either SELCON3 or SESCA algorithm. The raw CD data, expressed in millidegrees (m°), were converted to mean molar ellipticity (Δε) using the formula:

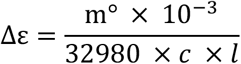

where m° is the measured ellipticity in millidegrees, c is the protein concentration in mol/L, and l is the path length of the cuvette in cm.

### 7. Kinetic analysis of YRDC and its mutants using LC–MS

The kinetic parameters of YRDC and its mutants were determined by measuring its activity against varying substrate concentrations. Due to the instability and short half-life of TC-AMP, the kinetics of the enzyme YRDC and its mutants were assessed by quantifying AMP, which is the byproduct of the reaction. For L-threonine substrate in the YRDC assay, concentrations of 1, 2, 5, 10, 25 and 50 mM were employed, while for ATP, concentrations of 10, 20, 50, 100 and 250 µM were tested. Reactions were initiated by adding 50 nM YRDC to a solution containing 250 µM ATP (for L-threonine kinetics) and 50 mM L-threonine (for ATP kinetics), along with 50 mM Tricine (pH 8.5), 10 mM MgCl_2_, and 50 mM NaHCO_3_. All reactions were carried out at 37 °C and quenched at specific time points within one hour by adding 500 mM EDTA at a 1:10 ratio to the reaction volume. The samples were then pipetted to a 96-well plate for LC–MS analysis.

LC–MS analysis was performed on an ACQUITY™ Arc™ system (Waters™, United States) equipped with a Quaternary Solvent Manager-R, Sampler Manager FTN-R, 2489 UV/Vis Detector, and QDa Detector. Chromatographic separation was achieved using an Acquity Premier HSS T3 (100Å, 1.8 μm, 2.1 × 100 mm) column (Waters™) maintained at 40 °C, while samples were kept at 4 °C throughout the analysis. The mobile phase consisted of buffer A (H_2_O+ 0.1% FA) and buffer B (MeOH), with gradient elution as follows: 0–1.8 min, 99% A; 1.8–7.2 min, 99%–44% A; 7.2–12.6 min, 44%–1% A; and 12.6–18 min, 1%–99% A. The flow rate was 0.4 mL/min, the injection volume was 10 µL, and detection was carried out at 260 nm. Data acquired in single ion monitoring (SIR) mode set to detect AMP and Tc-AMP were integrated and processed by MassLynx V4.2. The MS was operated in positive polarity mode (ESI+). The final AMP concentration was quantified by measuring the area under the curve in ESI+ mode and converting it to concentration using a calibration curve generated under identical experimental conditions. All assays were performed in triplicate, and data are presented as mean ± standard deviation. Substrate saturation curves were generated by varying the substrate concentration while keeping the co-substrate concentration constant. The data were then fitted to kinetic model:

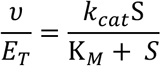

Where *υ* represents the initial reaction rate and *E_T_* denotes the total enzyme concentration. *K_M_* is the apparent Michaelis constant, *k_cat_* is the steady-state turnover number, and *S* corresponds to the concentration of the variable substrate.

### 8. Decomposition of TC-AMP

The reaction was initiated by adding 1 μM of YRDC to a mixture containing 100 mM L-threonine, 200 μM ATP, 50 mM NaHCO_3_, and reaction buffer (50 mM Tricine pH 8.5, 10 mM MgCl_2_). Reactions were performed at 37 °C for 1 minute, then quenched with 500 mM EDTA at a 1:10 ratio. The enzyme was removed from the mixture using a centrifugal filter unit (3,000 MWCO, Amicon®Ultra) and centrifuged at 4,000 g for 5 minutes at 4 °C. A 10 μL aliquot of the reaction mixture was injected to the system to measure TC-AMP every 20 minutes. The resulting TC-AMP concentration was quantified using LC–MS based on the method described above. Experiments were performed in triplicate, and the data are presented as the mean ± standard deviation.

### 9. Native Mass spectrometry for YRDC WT and mutants related to GAMOS

Proteins were buffer exchanged into 50 mM ammonium acetate (pH 6.8) using centrifugal ultrafiltration devices (10 kDa MWCO; [Amicon Ultra-4]) to remove non-volatile salts. For analysis under native conditions, the protein was diluted to 1 µM in 50 mM ammonium acetate. For unfolded (denaturing) conditions, the protein was diluted to 1 µM in 50% (v/v) acetonitrile containing 1% (v/v) formic acid. Samples were introduced by direct infusion at 20 µL min⁻¹ into a Waters Xevo G2-TOF mass spectrometer operated in positive-ion electrospray mode. Typical source parameters were capillary voltage 1.5–3.0 kV, sampling cone 200 V, and desolvation gas 300 L h⁻¹. Spectra were acquired over *m/z* 500–6000. Data processing was performed in MassLynx (Waters), with manual inspection of charge-state distributions to determine zero-charge molecular masses.

### 10. Crosslinking experiments with BS3

Crosslinking reactions (final YRDC 40 μM) were assembled in the exchange buffer at 20 °C with gentle agitation and initiated by addition of freshly prepared BS-3 (bis(sulfosuccinimidyl)suberate-d0, Thermo Scientific) to final concentrations of 0.2 mM or 3 mM. Aliquots were removed at 10, 30, and 60 min and quenched with Tris-HCl to a final concentration of 50 mM. SDS-PAGE loading buffer was added to each sample, and the mixtures were heated at 95 °C for 2 min prior to analysis by SDS-PAGE. Bands corresponding to the YRDC dimer from the 0.2 mM BS-3 condition at 10, 30, and 60 min, and the YRDC monomer from the 0.2 mM BS-3 10-min timepoint were excised and submitted to the Mass Spectrometry Facility, (University of St Andrews) for analysis.

Bands corresponding to expected size were excised and analysed by mass spectrometry (Supplementary Figure 10a). The samples were destained, reduced with dithiorethritol, alkylated with iodoacetamide, and trypsin-digested overnight at 37 °C. The resulting peptides were extracted in 1 % formic acid and evaporated to dryness in a Speedvac. The sample was resuspended in 0.1 % TFA and one quarter of the sample analysed by nano liquid chromatography-tandem mass spectrometry (nLC-MSMS) in trap elute configuration on a Thermo Orbitrap Fusion Lumos mass spectrometer. Peptides were bound to the trap column, and salts washed to waste before the trap switched in line with the PepMap Easyspray 15 cm analytical column, and peptides were eluted with a linear gradient of increasing acetonitrile 0.1% FA over 45 minutes at 300 nL min^−1^. Data were collected from 400 – 1400 *m/z* for MS at a resolution of 120000, data dependant acquisition was used to select peptides for fragmentation with ETD and HCD activation, with MSMS fragmentation spectra collected on the orbitrap at 60000 resolution. The proteins were initially identified by searching against a database of 8005 sequences, including the sequence of YrdC, using Mascot database search engine, v.2.8.1. The cross-linked peptides were identified against the sequence of YrdC, again with Mascot v2.8.1 utilising the crosslinking functionality against the YrdC sequence, with DSS/BS3 crosslinker against K residues, carbamidomethyl on cysteines, and tryptic cleavage with possible missed cleavages. Crosslinking peptide identification was also checked with MeroX 2.0.1.4 software for confirmation.

### 11. Native polyacrylamide (Native–PAGE) Electrophoresis

Protein samples, either SEC-purified or purified via a second nickel-affinity step, were adjusted to a final concentration of 0.5 mg mL^−1^ in detergent-free NativePAGE™ Sample Buffer (4X) (Invitrogen™, Thermo Fisher Scientific), which the final 1X solution contains 50 mM Bis-Tris (pH 7.2, adjusted with 6 N HCl), 50 mM NaCl, 10% (w/v) glycerol, and 0.001% (w/v) Ponceau S. Polymerization behaviour of the wild-type protein and its mutants was assessed by referenced against the marker NativeMark™ Unstained Protein Standard (Invitrogen™, Thermo Fisher Scientific), which contains eight reference bands spanning 20–1,200 kDa.

Prior to loading, pre-chilled cathode buffer (50 mM Bis-Tris, 50 mM Tricine, pH 7.0, 0.02% (w/v) Coomassie G-250) was added to the inner chamber, and anode buffer (50 mM Bis-Tris, 50 mM Tricine, pH 7.0) of the gel electrophoresis apparatus, as per the manufacturer’s instructions. The prepared samples were loaded into each well of NativePAGE™ Bis-Tris Mini Protein Gels (4–16%, 1.0 mm; Invitrogen™, Thermo Fisher Scientific) in a volume of 10 µL. The experiment was conducted at 4 °C, with the initial gel run performed at a constant voltage of 150 V for 60 min, followed by 250 V for 120 min. To enhance visibility, the gel was stained with Bio-Safe™ Coomassie Blue G-250 (Bio-Rad) prior to imaging using ChemiDoc™ MP Imaging System (Bio-Rad, United States).

### 12. Mass photometry for oligomeric-state determination

Measurements were conducted using a OneMP MP instrument (Refeyn Ltd). Microscope slides (70 × 26 mm) and silicone gaskets, designed to hold sample droplets, were cleaned twice with isopropanol followed by Milli-Q water, then air-dried using a stream of pressurized air. All sample preparations were completed immediately prior to measurement. Instrument calibration was performed using NativeMark protein standards. Data acquisition was carried out with the AcquireMP software. Protein stocks were diluted to a final concentration of 100 nM. To establish focus, 8 µL of reaction buffer was initially added to a well, and the optimal focal plane was identified and locked using the autofocus function. For each measurement, 2 µL of protein solution (100 nM) was mixed with 8 µL of buffer. To assess the impact of ligand binding on polymerisation, 2 µL of a solution containing 200 mM L-threonine and 2 mM ATP was introduced to the protein samples. MP videos were recorded over a 180-second period. Data analysis, including generation of mass distribution histograms, was conducted using DiscoverMP software.

### 13. Detection of pyrophosphate (PPi) and by-product AMP by ^31^P-NMR spectroscopy

Samples were prepared in a total volume of 2 mL in reaction buffer (pH 8.0) containing final concentrations of L-threonine (2 mM), NaHCO_3_ (100 mM), ATP (1 mM), and YRDC (1 μM), and incubated for 1 hour at either room temperature or 37 °C. To assess background signals and verify peak assignment, positive controls containing AMP (1 mM) and PPi (1 mM) were prepared in parallel under identical conditions. Following incubation, samples were centrifuged for 5 minutes at 13,000 rpm to remove enzyme, and D₂O was added at a ratio of 1:10 (v/v) for field locking. ^31^P-NMR spectroscopy was performed at room temperature to monitor ATP, AMP, and inorganic PPi using a Bruker AVIII 500 MHz spectrometer equipped with a double-resonance Prodigy BBO CryoProbe. Spectra were acquired in 5 mm Wilmad® precision NMR tubes (Z412007; 7 inches length; rated for 800 MHz) under the following parameters: 128 scans, a relaxation delay of 1.5 s, and 4 dummy scans. PPi, the principal product of the reaction, was used as a reference to validate product detection, whereas AMP, a secondary by-product of ATP turnover, served as an internal marker to monitor nucleotide conversion during catalysis. Spectral data were processed and analysed using MestReNova x64 and NMRium.

### 14. L-threonine-carbamate formation using ^15^N *L*-threonine and ^13^C-NaHCO_3_

NMR experiments were acquired at room temperature on a Bruker AVIII-HD 700 MHz spectrometer equipped with a Prodigy TCI CryoProbe for triple-resonance bio-NMR, using 5 mm Wilmad precision tubes (Z412007; 7 in length; frequency rating 800 MHz). Each sample had a volume of 600 µL and was prepared in D₂O reaction buffer comprising 50 mM Tricine, pH 8.5 and 10 mM MgCl₂, with final concentrations of [¹⁵N]-L-threonine (50 mM; 98% ¹⁵N, Goss Scientific), [¹³C]-NaHCO₃ (50 mM; 99% ¹³C, Goss Scientific), ATP (1 mM), and YRDC (50 µM); mixtures were incubated at 37 °C for the indicated time points prior to acquisition. The following datasets were collected: 1D ¹H (4 scans, relaxation delay 1.0 s, ¹H carrier 5.0 ppm), 1D ¹³C with broadband decoupling (200 scans, relaxation delay 2.0 s, ¹³C carrier 115 ppm), gradient-selected ¹H–¹⁵N HMBC optimized for J = 8 Hz (2 scans, relaxation delay 1.5 s, ¹H carrier 6.2 ppm), and gradient-enhanced ¹H–¹³C HMBC (gHMBC) targeting long-range carbon–proton correlations (2 scans, relaxation delay 1.5 s, ¹H carrier 6.0 ppm, ¹³C decoupling carrier 115 ppm). Spectral data were processed and analysed with MestReNova (x64) and NMRium.

### 15. Direct L-threonine binding demonstrated using STD NMR

NMR experiments to determine the sequence of substrate binding were performed at 37 °C, on a Bruker AVIII-HD 700 MHz spectrometer equipped with a Prodigy TCI CryoProbe for triple-resonance bio-NMR, using 5 mm Wilmad precision tubes (Z412007; 7 in length; frequency rating 800 MHz). The STD experiment was acquired using the STDDIFFESGP parameter set from Bruker TopSpin library^46^. Prior to carrying out experiments, protein was buffer exchanged into 20 mM potassium Phosphate pH 8.5 with 100 mM KCl using a 10 kDa centrifugal ultrafilter (Amicon, Merck Millipore). Each sample had a volume of 600 µL and was prepared with the addition of 10% D₂O to a 20 mM potassium Phosphate pH 8.5 buffer with 100 mM KCl and 10 mM MgCl₂. First, a 1H NMR spectra of YRDC was acquired to determine a suitable protein resonance value to be irradiated during the STD experiment (0.22 ppm – see Supplementary Figure 10 – top panel, as well as Figure 5). Then, a reference off-resonance spectrum for the sample containing both threonine and YRDC but with saturation pulse at 40 ppm was acquired, followed by acquisition of spectra with a saturation pulse at 0.22 ppm, and saturation times of 0.5 s and 1s, each with a recycle delay D1 of 3s. Spectral data were processed and analysed with MestReNova (x64) and NMRium.

## Data Availability

The data necessary to support the findings of this study are available within the main text and supplementary information. Supplementary information file contains all raw data for CD, DSF, kinetic assays. Protein mass spectrometry data were deposited on Figshare https://figshare.com/s/ce133096d5ac8da766a6.

## Acknowledgements

T.N.A.T. was funded by Medical Research Scotland and Nucana plc (PhD-50264-2020), C.M.C. was funded by the BBSRC Pathfinder IAA BB/X511183/1 to the University of St Andrews), S.L.S. and the instrument used for protein mass spectrometry was funded by BB/T017686/1. R.S. is funded by the Wellcome Trust (223816/Z/21/Z). G.M.Z was funded by the Melville Trust PhD Scholarship and Nucana plc. We would like to express our gratitude to the reviewers of this paper, especially on their insights regarding NMR experiments.

## Contributions

Manuscript was written by T.N.A.T., but all authors contributed to the final form. All authors have given approval to the final version of the manuscript. Specific contributions are as follows: T.N.A.T. designed and performed experiments, interpreted data, wrote manuscript; G.M.Z. and A.L.D. contributed to small molecule mass spectrometry, interpreted data, revised manuscript; C.J.H. helped with protein biophysical assays; T.L. helped design, perform and interpret NMR data,; S.S. and S.L.S. performed protein mass spectrometry experiments, analysed data, revised manuscript; R.S. acquired MP data, analysed data, revised manuscript; D.J.H. and C.M.C. co-supervised T.N.A.T., participated in project conception, analysed and interpreted data, revised manuscript.

