## Supporting information for "Dimerization and Threonine-Dependent Stabilization Govern Human YRDC Catalysis"

### Supplementary methods

DNA sequence - YRDC$_{WT}$

GGC GCC CGT CTG TTG CGT TTA CCG GGA TCA GGA GCA GTT CAG GCG GCC
AGT CCA GAA CGT GCT GGG TGG ACT GAA GCG CTG CGC GCT GCG GTC GCC
GAG TTA CGC GCA GGG GCC GTT GTA GCT GTT CCC ACA GAT ACC CTT TAC
GGT TTG GCC TGT GCA GCA AGC TGC TCC GCT GCG TTA CGC GCC GTG TAC
CGT CTG AAG GGC CGC TCA GAA GCG AAA CCT TTG GCC GTA TGT TTG GGC
CGC GTC GCT GAT GTA TAT CGT TAT TGC CGT GTA CGC GTT CCC GAA GGT
CTG TTA AAA GAC TTA TTG CCT GGG CCT GTG ACC TTA GTA ATG GAA CGC
TCG GAG GAG CTT AAT AAA GAC TTG AAC CCT TTC ACG CCA CTG GTA GGT
ATC CGC ATT CCA GAC CAC GCT TTC ATG CAA GAC TTG GCT CAA ATG TTC
GAG GGA CCC TTA GCT TTG ACA AGC GCA AAT TTA TCC TCT CAG GCC AGC
TCG CTT AAC GTC GAA GAG TTT CAG GAT CTG TGG CCA CAG CTG AGT TTG
GTG ATT GAC GGG GGT CAG ATT GGC GAT GGC CAG AGC CCT GAG TGC CGT
TTG GGG TCA ACC GTG GTC GAT CTT TCG GTC CCT GGG AAA TTT GGC ATT
ATC CGT CCC GGA TGC GCC CTT GAG TCA ACG ACG GCC ATC TTA CAA CAG
AAA TAC GGC TTA CTT CCA AGT CAC GCT TCC TAC TTG TGA

#### Protein sequences

| Protein sequence |
| --- |
| The full-length amino acid sequence of the human YRDC protein (UniProt **Q86U90**) is shown below, with the grey colour region indicating the 43 amino acids truncated at the N-terminal.<br><br>5'–<br>MSPARRCRGMRAAVAASVGLSEGPAGSRSGRLFRPPSPAPAAPGARLLRLPGSGAVQAAS<br>PERAG<br>WTEALRAAVAELRAGAVVAVPTDTLYGLACAASCSAALRAVYRLKGRSEAKPLAVCLGR<br>VADVYRYCRVRVPEGLLKDLLPGPVTLVMERSEELNKDLNPFTPLVGIRIPDHAFMQDLA<br>QMFEGPLALTSANLSSQASSLNVEEFQDLWPQLSLVIDGGQIGDGQSPECRLGSTVVDLSV<br>PGKFGIIRPGCALESTTAILQQKYGLLPSHASYL–3' |
| The wild-type amino acid sequence of YRDC (Δ43 amino acid) (Uniprot **Q86U90**) was used in this study, with the grey region representing the region cleaved by TEV protease during protein purification<br><br>5'–<br>HHHHHHDYDIPTTENLYFQGARLLRLPGSGAVQAASPERAGWTEALRAAVAELRAGAVV<br>AVPTDTL<br>YGLACAASCSAALRAVYRLKGRSEAKPLAVCLGRVADVYRYCRVRVPEGLLKDLLPGPV<br>TLVMERSEELNKDLNPFTPLVGIRIPDHAFMQDLAQMFEGPLALTSANLSSQASSLNVEEF<br>QDLWPQLSLVIDGGQIGDGQSPECRLGSTVVDLSVPGKFGIIRPGCALESTTAILQQKYGLL<br>PSHASYL–3' |

### Establishment of an LC-MS assay to determine the activity of YRDC

TC-AMP is an unstable product, which is hydrolysed to AMP and a cyclic threonine intermediate[1, 2]. Therefore, performing assays and monitoring Tc-AMP is technically challenging, and we set out to develop a method to allow AMP quantification as a proxy for Tc-AMP formation.

First, to investigate the catalytic activity of human YRDC and the stability of the TC-AMP product, LC–MS was carried out under defined reaction conditions (50 mM Tricine, pH 8.5, 10 mM $MgCl_2$). The reaction was initiated at 37 °C and allowed to proceed for one minute, yielding a TC-AMP product with a retention time (RT) of approximately 6.5 minutes. Monitoring the same sample at 20-minute intervals revealed a progressive decline in the TC-AMP signal. The resulting degradation profile fit a one-phase exponential decay model, with a calculated half-life of approximately 35 minutes. Therefore, in our standard reaction conditions each time point is quenched with MeOH, solvent is evaporated and samples resuspended in $H_2O$ for injection in the LC-MS. During this sample preparation time no TC-AMP remained in solution.

The formation of the TC-AMP product was further confirmed by liquid chromatography–electrospray ionisation– high resolution mass spectrometry (LC–ESI–HRMS), which detected a negatively charged species $[M–H]^-$ with a measured mass consistent with the molecular formula of TC-AMP (theoretical exact mass: 491.09; observed: 491.0933, Figure 4 and Supplementary Figure 7). A signal corresponding to AMP—a known by-product of TC-AMP degradation—was also identified as a $[M–H]^-$ peak at m/z = 346.0558, and an additional peak corresponding to the monosodium dianion species $[M+Na–2H]^-$ of TC-AMP was observed at m/z = 513.0753. Supporting this result, $^{31}P$ NMR spectroscopy performed under identical reaction conditions detected the characteristic AMP signal, reinforcing the presence of TC-AMP. Importantly, when the reaction was conducted at room temperature rather than at 37 °C, the AMP signal was absent, indicating that catalysis by human YRDC occurs specifically at physiological temperature. These data confirm that human YRDC catalyses the formation of TC-AMP from L-threonine, bicarbonate, and ATP under the specified conditions.

### Supplementary tables

| VARIANTS | | $T_m$ (DSF) (°C) | $T_m$(CD) (°C) |
|---|---|---|---|
| **$YRDC_{WT}$** | No ligands | 49.9 ± 0.5 | 47.0 ± 0.3 |
| | Enzyme·ATP·Thr | 59.4 ± 0.5 | 53.0 ± 0.2 |
| **$YRDC_{T46A}$** | No ligands | 49.4 ± 0.4 | 39.9 ± 0.9 |
| | Enzyme·ATP·Thr | 50.9 ± 0.2 | 53.6 ± 0.1 |
| **$YRDC_{C78A}$** | No ligands | 48.6 ± 0.2 | 45.6 ± 0.3 |
| | Enzyme·ATP·Thr | 58.4 ± 0.1 | 52.8 ± 0.2 |
| **$YRDC_{R130A}$** | No ligands | 50.1 ± 0.1 | 41.2 ± 0.3 |
| | Enzyme·ATP·Thr | 50.6 ± 0.1 | 46.6 ± 0.2 |
| **$YRDC_{S195A}$** | No ligands | 54.2 ± 0.5 | 47.1 ± 0.2 |
| | Enzyme·ATP·Thr | 56.2 ± 0.1 | 52.8 ± 0.1 |
| **$YRDC_{R210K}$** | No ligands | 52.9 ± 0.4 | 46.6 ± 0.3 |
| | Enzyme·ATP·Thr | 55.4 ± 0.4 | 53.3 ± 0.2 |
| **$YRDC_{A41V}$** | No ligands | 48.3 ± 0.1 | 47.0 ± 0.5 |
| | Enzyme·ATP·Thr | 53.5 ± 0.8 | 52.6 ± 0.4 |
| **$YRDC_{I178T}$** | No ligands | 35.9 ± 0.2 | 24.1 ± 4.7 |
| | Enzyme·ATP·Thr | 43.4 ± 0.6 | 41.3 ± 1.8 |
| **$YRDC_{\Delta L222}$** | No ligands | 44.5 ± 0.1 | 37.0 ± 0.6 |
| | Enzyme·ATP·Thr | 51.3 ± 0.6 | 50.9 ± 0.7 |

Supplementary Table 1: Melting temperature comparison | Apparent melting temperatures ($T_m$) of wild-type YRDC and its variants with and without ligands (500 µM ATP and 50 mM L-threonine), measured by DSF and CD. Measurements were performed under identical buffer conditions; DSF data were collected over 25–95 °C, and far-UV CD spectra were acquired from 190–350 nm. Reported values are mean ± s.e.m.

Supplementary Table 2: Equilibrium dissociation constants

**Dissociation constants ($K_D$) from DSF for YRDC and variants across ligand conditions.** $K_D$ values were derived from DSF by varying the concentration of a single substrate while holding the other components as specified for each condition. Data are mean ± s.e.m. (n = 3). Experimental details are provided in Methods. **n.d.** indicates parameters not determined. Some of the error values are high as full binding saturation could not be obtained in the concentration ranges that could be experimentally tested. They should not be interpreted as quantitative values but as an indication that proteins are capable of binding ligands

, albeit with low affinity.

| Variants | Substrate | Conditions | $K_D$ values | Unit |
|---|---|---|---|---|
| $YRDC_{WT}$ | ATP | Buffer | 1998 ± 1064 | µM |
| | | With *L*-threonine | 77.3 ± 29.4 | |
| | L-threonine | Buffer | 31.1 ± 12.1 | mM |
| | | With ATP | 28.4 ± 12.3 | |
| | | With ATP and $NaHCO_3$ | 12.2 ± 8.5 | |
| | $NaHCO_3$ | Buffer | n.d | |
| | | With L-threonine | 38.3 ± 12.9 | |
| | | With ATP and L-threonine | 14.6 ± 5.6 | |
| | AMP | Buffer | 6.1 ± 1.8 | |
| | Pyrophosphate | Buffer | 7.1 ± 0.9 | |
| | L-cysteine | Buffer | 7.1 ± 3.9 | mM |
| | | With ATP | 17.5 ± 7.5 | |
| | L-serine | Buffer | 12.5 ± 12.3 | |
| | | With ATP | 31.2 ± 17.5 | |
| | L-alanine | Buffer | 18.0 ± 19.8 | |
| | | With ATP | 25.7 ± 19.8 | |
| | L-valine | Buffer | 14.8 ± 13.7 | |
| | | With ATP | 10.3 ± 7.1 | |
| $YRDC_{T46A}$ | ATP | With *L*-threonine | n.d | |
| | *L*-threonine | With ATP | n.d | |
| | $NaHCO_3$ | With ATP and *L*-threonine | n.d | |
| $YRDC_{C78A}$ | ATP | Buffer | n.d | |
| | | With *L*-threonine | 133.0 ± 29.7 | µM |
| | *L*-threonine | With ATP | 1.2 ± 0.3 | mM |
| | $NaHCO_3$ | With ATP and *L*-threonine | 3.7 ± 2.0 | |
| $YRDC_{R130A}$ | ATP | With L-threonine | n.d | |
| | *L*-threonine | With ATP | n.d | |
| | $NaHCO_3$ | With ATP and *L*-threonine | n.d | |
| $YRDC_{S195A}$ | ATP | With L-threonine | 48.9 ± 37.8 | µM |
| | *L*-threonine | With ATP | 17.2 ± 9.8 | mM |
| | $NaHCO_3$ | With ATP and *L*-threonine | n.d | |
| $YRDC_{R210K}$ | ATP | With L-threonine | 147.5 ± 111.8 | µM |
| | *L*-threonine | With ATP | 3.4 ± 1.8 | mM |
| | $NaHCO_3$ | With ATP and *L*-threonine | 53.9 ± 28.3 | |

| | | | | |
|---|---|---|---|---|
| $YRDC_{A41V}$ | ATP | Buffer | n.d | |
| | *L*-threonine | Buffer | 124.3 ± 74.6 | mM |
| | | With ATP | 0.3 ± 0.1 | |
| | $NaHCO_3$ | With ATP and *L*-threonine | 6.5 ± 2.0 | |
| $YRDC_{I178T}$ | ATP | With *L*-threonine | n.d. | |
| | *L*-threonine | Buffer | n.d. | mM |
| | | With ATP | 0.4 ± 0.1 | |
| | $NaHCO_3$ | With ATP and *L*-threonine | 8.3 ± 4.2 | |
| $YRDC_{\Delta L222}$ | ATP | With *L*-threonine | n.d | |
| | *L*-threonine | Buffer | 89.6 ± 44.9 | mM |
| | | With ATP | 2.1 ± 0.7 | |
| | $NaHCO_3$ | With ATP and *L*-threonine | 48.0 ± 9.9 | |

### Supplementary Table 3: Secondary structure elements determined by Circular dichroism

*The rationale for choosing different algorithms to fit data is as follows:*

SELCON3 for functional mutants as our samples were likely soluble globular proteins, and concentration/pathlength are accurate. Algorithm fits spectrum as a linear combination of spectra from a reference set of proteins with known structures. SESCA for GAMOS mutants as concentration of folded protein is uncertain (SESCA's Bayesian fit estimates a scaling factor). Furthermore, proteins are disordered/flexible. This analysis allows uncertainty estimates for the secondary structure fractions to report alongside point estimates.

| | ***SELCON***[1] ***algorithm*** | $YRDC_{WT}$ | $YRDC_{T46A}$ | $YRDC_{C78A}$ | $YRDC_{R130A}$ | $YRDC_{S195A}$ | $YRDC_{R210K}$ |
|---|---|---|---|---|---|---|---|
| *α-helix* | No ligands | 8.5 | 17.7 | 14.3 | 15.5 | 15.6 | 14.4 |
| | Enzyme·ATP·Thr | 9.6 | 2.6 | 7.8 | 2.6 | 2.6 | 5.9 |
| *β-sheet* | No ligands | 33.6 | 24.7 | 29.2 | 27.6 | 26.6 | 27.9 |
| | Enzyme·ATP·Thr | 37.1 | 43.0 | 35.9 | 43.0 | 43.0 | 33.7 |
| *turns* | No ligands | 14.6 | 14.7 | 11.6 | 13.7 | 12.4 | 13.4 |
| | Enzyme·ATP·Thr | 13.2 | 20.2 | 15.7 | 20.2 | 20.2 | 16.1 |
| *disordered* | No ligands | 42.3 | 44.6 | 43.0 | 42.0 | 45.1 | 42.9 |
| | Enzyme·ATP·Thr | 39.3 | 34.2 | 40.0 | 34.2 | 34.2 | 40.7 |

| **VARIANTS - SESCA**[1] **ALGORITHM** | | **α -helix** | **β-sheet** | **coil (disordered)** | **Scaling factor** |
|---|---|---|---|---|---|
| **$YRDC_{WT}$** | No ligands | 8.6 ± 5.0 | 23.7 ± 6.0 | 67.7 ± 7.3 | 1.2 ± 0.2 |
| | Enzyme·ATP·Thr | 11.7 ± 5.4 | 29.0 ± 6.7 | 59.3 ± 5.7 | 1.0 ± 0.2 |
| **$YRDC_{A41V}$** | No ligands | 4.9 ± 6.9 | 16.0 ± 11.6 | 79.0 ± 10.9 | 0.3 ± 0.1 |
| | Enzyme·ATP·Thr | 5.2 ± 7.0 | 17.5 ± 12.2 | 77.2 ± 10.8 | 0.4 ± 0.2 |
| **$YRDC_{I178T}$** | No ligands | 5.3 ± 6.3 | 18.8 ± 7.2 | 75.9 ± 9.5 | 1.1 ± 0.2 |
| | Enzyme·ATP·Thr | 5.2 ± 7.5 | 16.5 ± 13.2 | 78.3 ± 11.9 | 0.5 ± 0.3 |
| **$YRDC_{\Delta L222}$** | No ligands | 7.5 ± 7.1 | 14.6 ± 12.9 | 78.0 ± 11.4 | 0.5 ± 0.2 |
| | Enzyme·ATP·Thr | 5.3 ± 6.1 | 12.2 ± 8.9 | 82.5 ± 10.9 | 1.0 ± 0.2 |

*Secondary-structure composition of YRDCWT and GAMOS-related point mutants in the absence or presence of ligands, determined by circular dichroism. Far-UV/near-UV CD spectra (190–350 nm) were recorded at 20 °C in 20 mM sodium phosphate buffer (pH 8.0) containing 75 mM NaF. Measurements were performed without ligands and with ligands: ATP (500 µM) and L-threonine (50 mM). Spectra are reported as mean molar extinction (Δε; L•mol-1•cm-1•chromophores-1) versus wavelength. Deconvolution was performed on the ChiraKit platform using the SELCON (top) or SESCA (bottom) algorithm with the DS-dT base set. Values are presented as percentage composition (mean for SELCON and mean ± S.D. for SESCA ).*

### Supplementary Table 4: Kinetic parameters for $YRDC_{WT}$

| Substrates | $k_{cat}$ ($s^{-1}$) | $K_m$ (mM) | $k_{cat}/K_m$ ($mM^{-1}$ $s^{-1}$) |
|---|---|---|---|
| ATP | 0.022 ± 0.002 | 0.07± 0.01 | 0.30 ± 0.05 |
| *L*-threonine | 0.038 ± 0.002 | 7.8 ± 1.1 | 0.003 ± 0.001 |
| $NaHCO_3$ | 0.017 ± 0.002 | 3.3 ± 1.1 | 0.005 ± 0.002 |

***Kinetic parameters of human $YRDC_{WT}$ for ATP, L-threonine, and bicarbonate measured by LC–MS.*** *Initial rates were fit to the Michaelis–Menten equation to obtain $k_{cat}$, $K_m$, and $k_{cat}/K_m$. AMP was quantified as a surrogate for TC-AMP formation. Values are mean ± s.e.m. from n=3 independent experiments at 37°C. Exact substrate ranges, enzyme and ion concentrations, and LC–MS settings are provided in Methods.*

### Supplementary Table 5: Kinetic parameters for $YRDC_{WT}$ and mutants

| Variants | $k_{cat}$ ($s^{-1}$) | $K_{m\text{-}ATP}$ (μM) | $k_{cat}/K_{m\text{-}ATP}$ ($mM^{-1}\ s^{-1}$) | *Fold decrease from WT* |
|---|---|---|---|---|
| YRDC Wild type | 0.022 ± 0.002 | 73.6 ± 12.0 | 0.3 ± 0.05 | - |
| $YRDC_{T46A}$ | 0.02 ± 0.001 | 230.7 ± 217.6 | 0.09 ± 0.08 | 3 |
| $YRDC_{C78A}$ | 0.01 ± 0.001 | 69.8 ± 20.7 | 0.1 ± 0.05 | 3 |
| $YRDC_{R130A}$ | n.d | n.d | n.d | - |
| $YRDC_{S195A}$ | 0.005 ± 0.0009 | 711.8 ± 175.5 | 0.007 ± 0.002 | 43 |
| $YRDC_{R210K}$ | 0.005 ± 0.003 | 737.5 ± 521.2 | 0.007 ± 0.006 | 43 |
| $YRDC_{A41V}$ | 0.016 ± 0.002 | 110.8 ± 24.2 | 0.10 ± 0.04 | 3 |
| $YRDC_{I178T}$ | 0.004 ± 0.0003 | 79.3 ± 13.7 | 0.05 ± 0.01 | 6 |
| $YRDC_{\Delta L222}$ | 0.025 ± 0.004 | 124.6 ± 40.5 | 0.20 ± 0.07 | 2 |

***Kinetic parameters of YRDC WT and its variants (binding-site and GAMOS-related) with ATP as the varied substrate.*** *Initial rates were fit to the Michaelis–Menten equation to obtain $k_{cat}$, $K_m$, and $k_{cat}/K_m$ . AMP was quantified by LC–MS as a surrogate for TC-AMP formation. L-threonine and bicarbonate were held at saturating concentrations. Values are mean ± s.e.m. (n = 3).* ***n.d.*** *indicates parameters not determined within the tested ATP range. Experimental conditions are detailed in Methods.*

Supplementary Table 6: Expected and observed m/z values for different analytes.

| Molecule | Theoretical *m/z* | Experimental *m/z* | Ion |
|---|---|---|---|
| **TC-AMP** | 491.0959 | 491.0933 | [M-H]- |
| | 513.0771 | 513.0753 | [M+Na-2H]- |
| **AC-AMP** | 461.0839 | 461.0828 | [M-H]- |
| | 483.0652 | 483.0647 | [M+Na-2H]- |
| **CC-AMP** | 493.0536 | 493.0548 | [M-H]- |
| | 515.0363 | 515.0368 | [M+Na-2H]- |
| **SC-AMP** | 477.0802 | 477.0777 | [M-H]- |
| | 499.0600 | 499.0596 | [M+Na-2H]- |
| **VC-AMP** | 489.1148 | 489.1141 | [M-H]- |
| | 511.0985 | 511.0960 | [M+Na-2H]- |

**Exact mass and observed *m/z* values for TC-AMP and aminoacylcarbamoyl-AMP analogues measured by LC–ESI(−)–MS**. Reported ions including $[M-H]^-$ and $[M+Na-2H]^-$. Calculated monoisotopic neutral masses (Da) were obtained with ChemDraw; corresponding adduct *m/z* values were derived accordingly. **Abbreviations:** TC-AMP, threonylcarbamoyl-AMP; AC-AMP, alanylcarbamoyl-AMP; CC-AMP, cysteinylcarbamoyl-AMP; SC-AMP, serylcarbamoyl-AMP; VC-AMP, valylcarbamoyl-AMP.

### Supplementary Figures

#### Supplementary Figure 1: SDS–PAGE analysis of recombinant YRDC wild type and GAMOS-associated variants during purification.

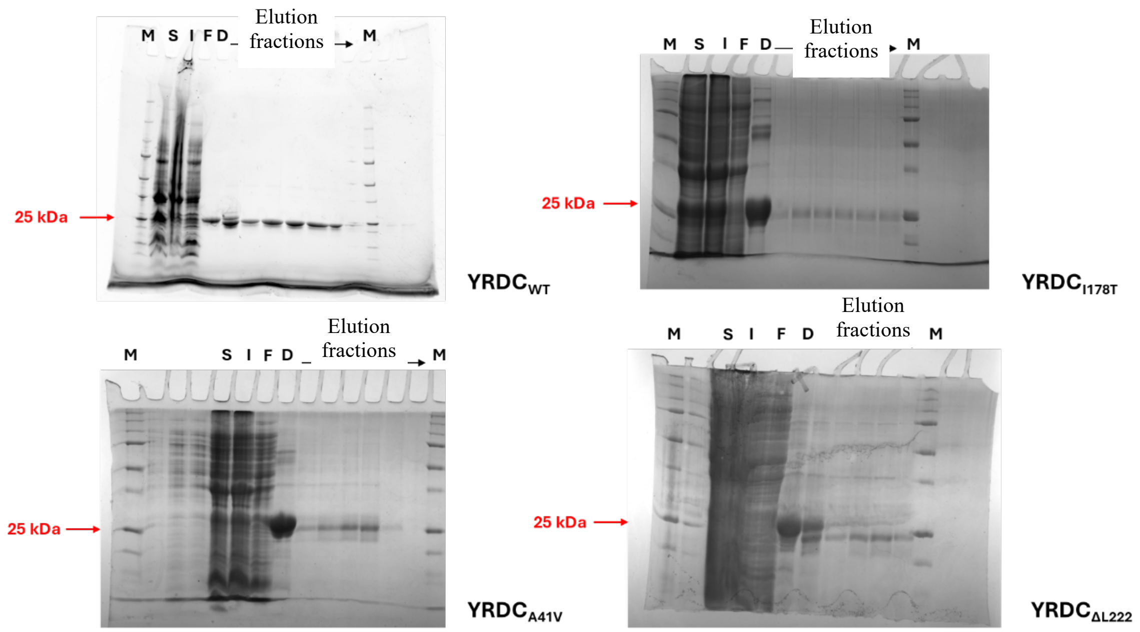

Coomassie-stained 10–20% Tris–glycine SDS–PAGE gels showing purification of recombinant YRDC$_{WT}$, YRDC$_{A41V}$, YRDC$_{I178T}$, YRDC$_{\Delta L222}$. Lane assignments are: M, molecular-mass marker (Unstained Protein Standard, Broad Range, 10–200 kDa); S, soluble fraction; I, insoluble fraction; F, flow-through; D, dialysed sample; and Elutions, sequential elution fractions. Red arrows indicate the YRDC band at approximately 25 kDa.

Supplementary Figure 2: DSF for all proteins with and without ligands

Supplementary Figure 2: Differential Scanning Fluorimetry (DSF) for all protein variants in the absence and presence of ligands as indicated.

<table>
<tr><td colspan="2">Thermal stability of YRDC and variants as a function of substrate concentration (DSF/SYPRO Orange). For each substrate condition, the left panel plots the derivative melting temperature ($T_m$) versus substrate concentration for $YRDC_{WT}$ and all mutants; points show mean ± s.e.m. (n = 3). The right panel shows the corresponding fluorescence traces (raw or normalized RFU vs temperature, 20–95 °C). One substrate was varied per experiment while other components were held as specified for that condition. SYPRO Orange was used as the reporter dye; samples were heated from 20 to 95 °C at a constant ramp rate. $T_m$ values were obtained from the first derivative of the melting curves. Full experimental details are provided in Methods.</td></tr>
<tr><td colspan="2">YRDC WT</td></tr>
<tr><td>
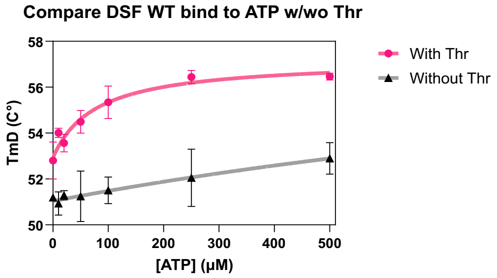

$K_d$ for binding without Threonine >500 µM, too high to accurately determine, estimated 1997.6 ± 10648.4.

$K_d$ with threonine present = 77.3 ± 29.4 µM</td>
<td>ATP had little or slight effect on the stability of the L-threonine–YRDC complex, as evidenced by an increase in the $T_m$ value from 55 °C with 50 mM L-threonine alone to 59 °C with 25 mM L-threonine in an ATP-saturated medium.

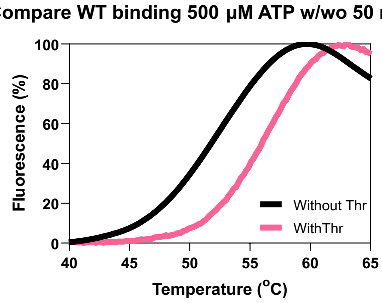
</td></tr>
<tr><td>
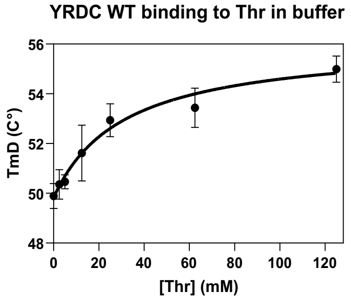

$K_d$ for threonine binding to apo protein = 31.1 ± 12.1 mM</td>
<td>
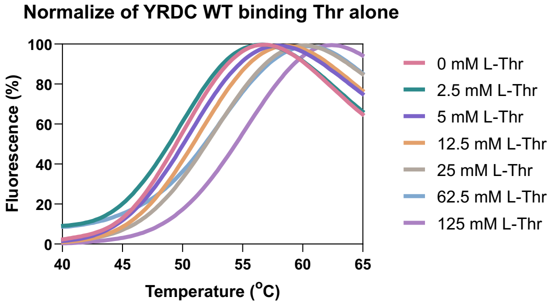
</td></tr>
<tr><td>YRDC WT binding to Thr in TAPS pH 8.5</td>
<td>
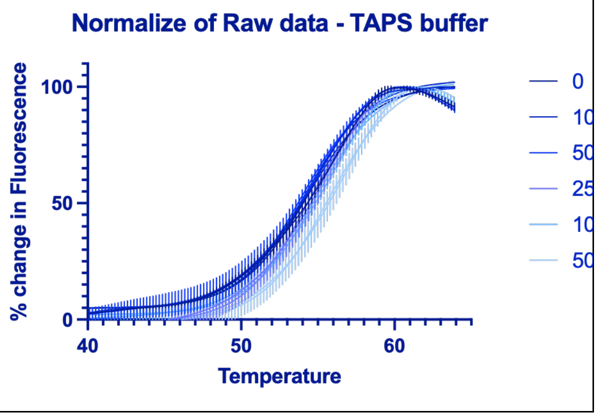
</td></tr>
</table>

### Supplementary Figure 2: DSF for all proteins with and without ligands

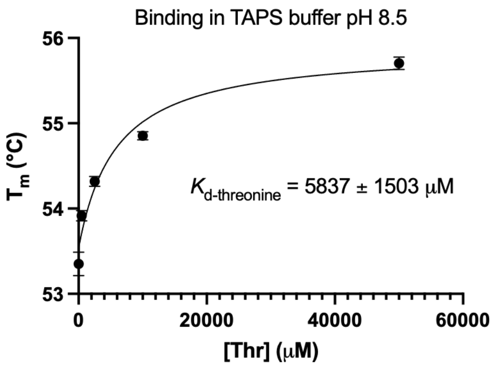

0

0

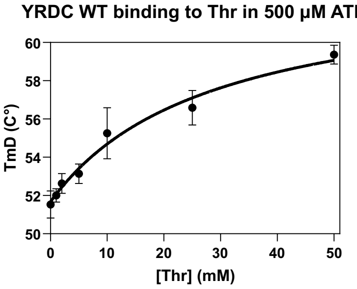

$K_d$ for threonine binding in the presence of ATP = 28.4 ± 12.3 mM

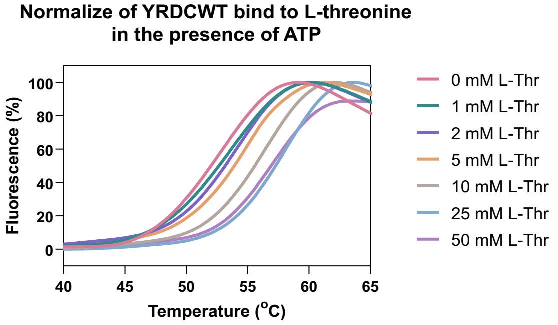

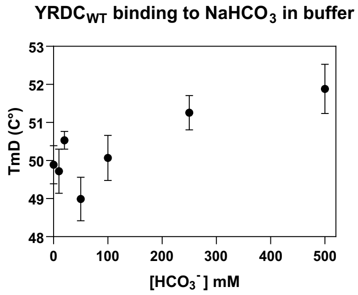

Trend is shown but $K_d$ for binding NaHCO3 in the absence of other ligands was >500 µM, too high to accurately determine.

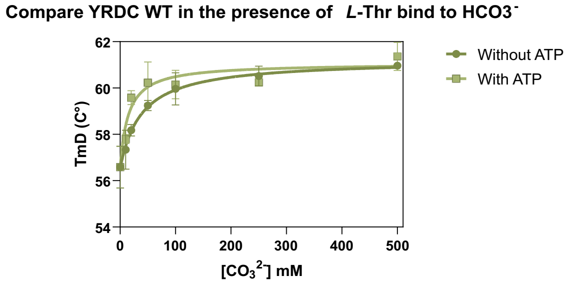

$K_d$ for binding with threonine = 38.3 ± 12.9 mM, $K_d$ with threonine and ATP present = 14.6 ± 5.6 mM.

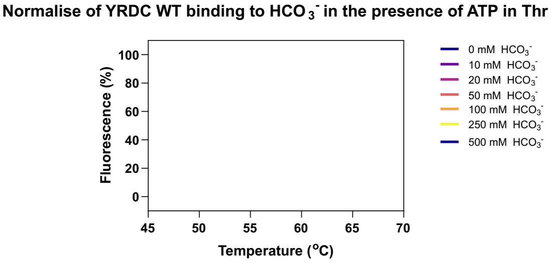

$T_m$ with threonine + ATP = 55.1 ± 1.1

$T_m$ with threonine + ATP + $HCO^{3-}$ = 61.1 ± 0.2

### Supplementary Figure 2: DSF for all proteins with and without ligands

YRDC T46A

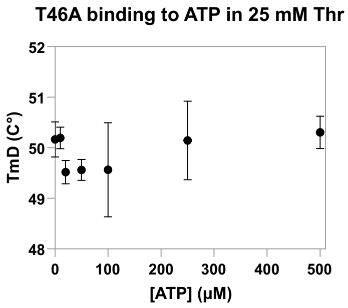

$K_d$ too high to accurately determine.

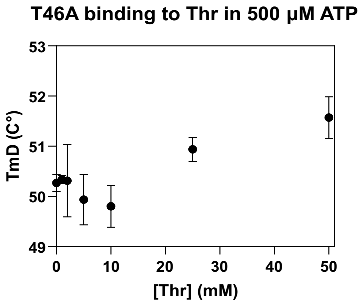

$K_d$ too high to accurately determine.

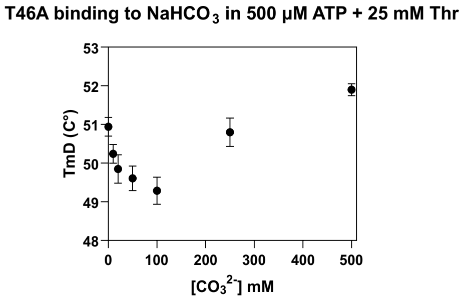

$K_d$ too high to accurately determine.

YRDC C78A

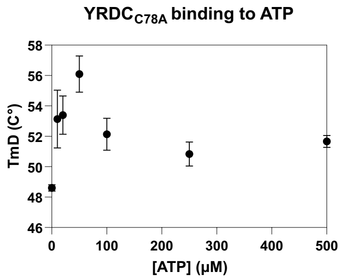

$K_d$ undetermined

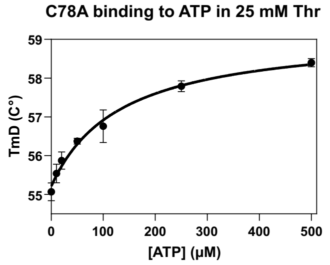

$K_d$ with threonine present = 133.0 ± 29.7 μM, undetermined in the absence of threonine.

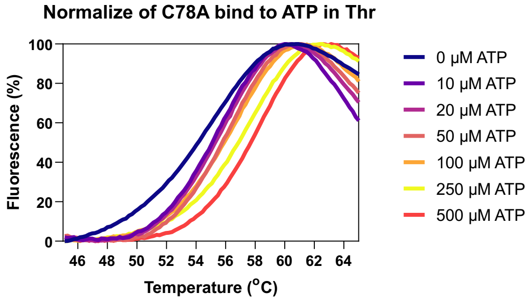

$T_m$ with threonine = 53.9 ± 0.1
$T_m$ with threonine + ATP = 57.6 ± 0.2

### Supplementary Figure 2: DSF for all proteins with and without ligands

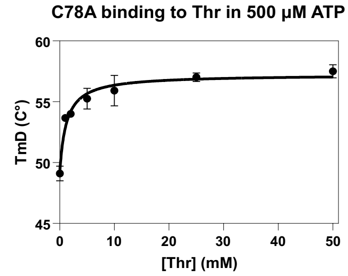

$K_d$ with ATP present = 1.2 ± 0.3 mM

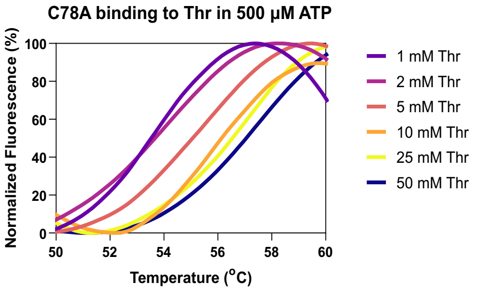

$T_m$ with ATP = 53.7 ± 0.2

$T_m$ with threonine + ATP = 57.3 ± 1.1

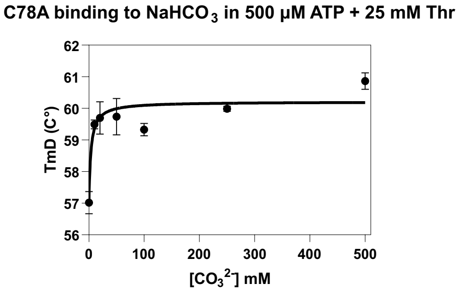

$K_d$ with ATP and threonine present = 3.7 ± 2.0 mM

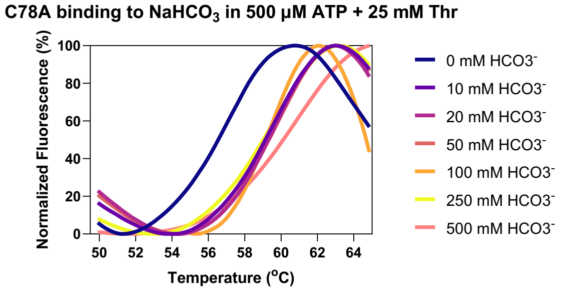

$T_m$ with ATP + threonine = 56.1 ± 0.3

$T_m$ with threonine + ATP + $HCO^{3-}$= 60.4 ± 0.1

#### YRDC R130A

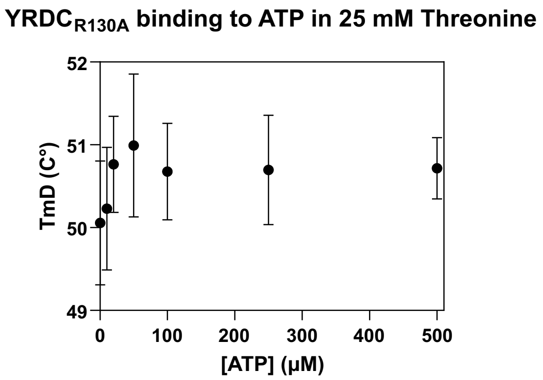

$K_d$ undetermined

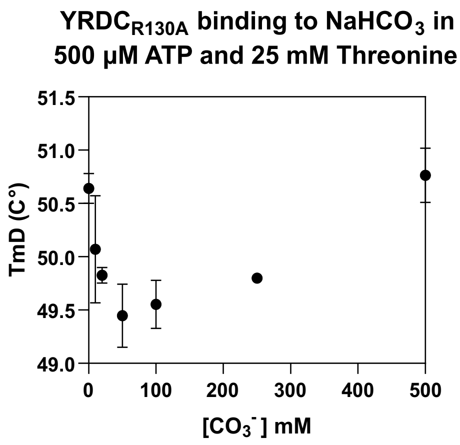

$K_d$ undetermined

#### YRDC S195A

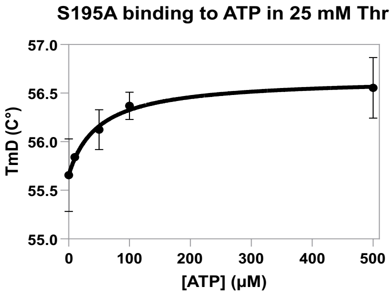

$K_d$ with threonine present =48.9 ± 37.8 μM

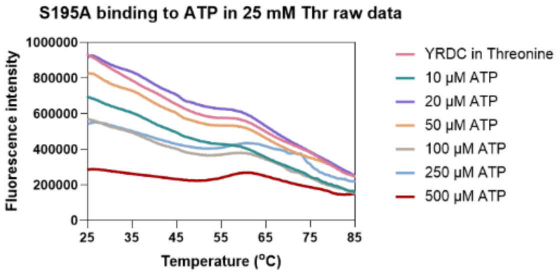

### Supplementary Figure 2: DSF for all proteins with and without ligands

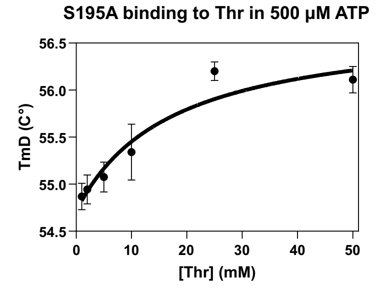

$K_d$ with ATP present =17.2 ± 9.8 mM

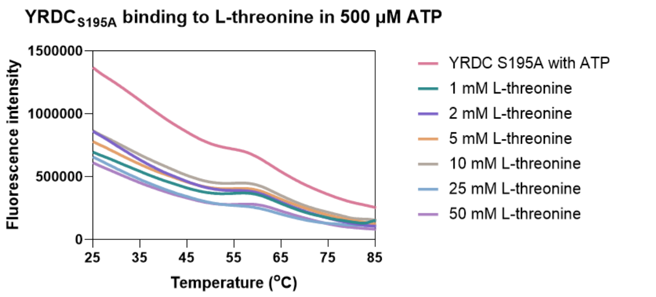

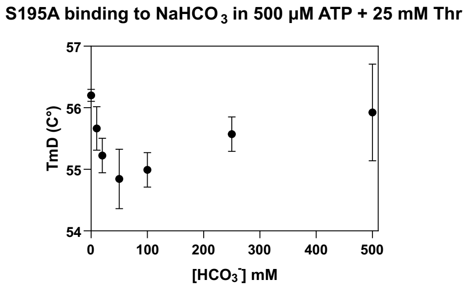

$K_d$ undetermined

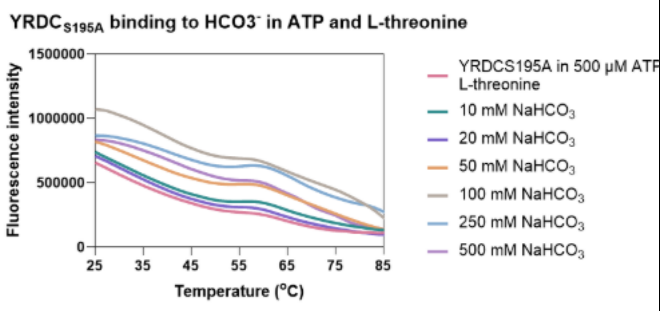

YRDC R210K

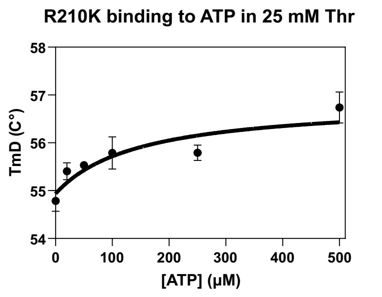

$K_d$ with threonine present =147.5 ± 111.8 μM

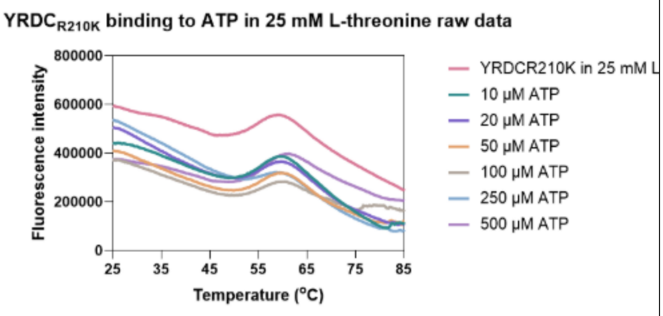

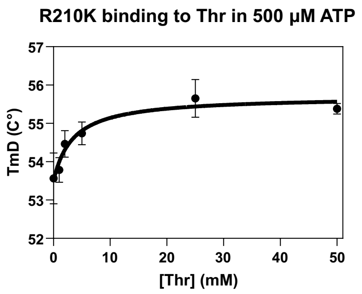

$K_d$ with ATP present =3.4 ± 1.8 mM

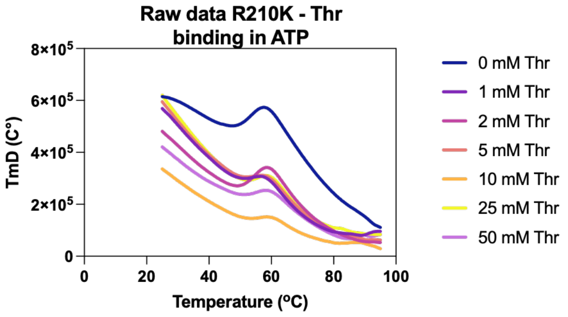

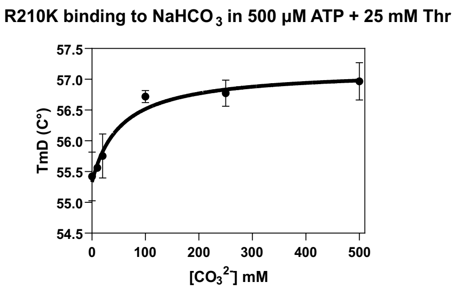

$K_d$ with threonine and ATP present =53.9 ± 28.3 mM

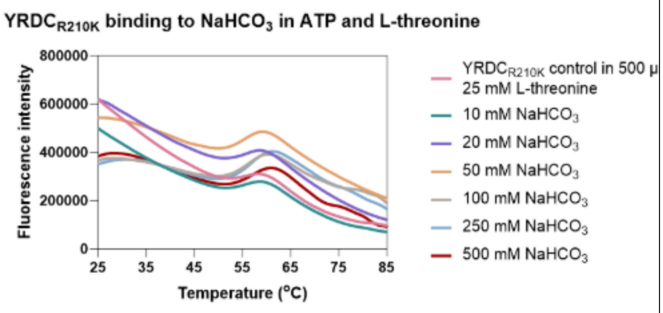

YRDC A41V

### Supplementary Figure 2: DSF for all proteins with and without ligands

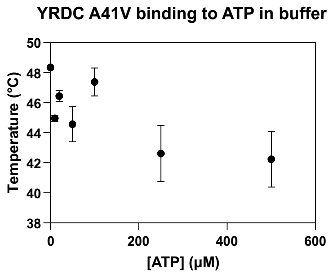

$K_d$ undetermined

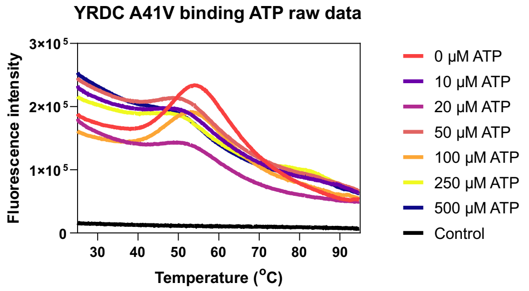

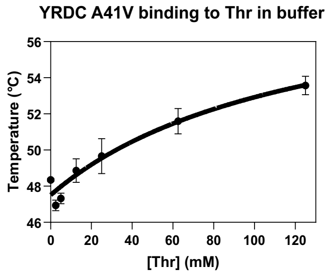

$K_d$ for binding threonine = 124.3 ± 74.6 mM

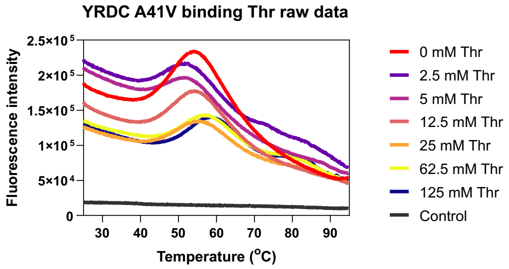

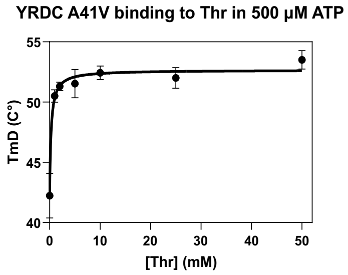

$K_d$ with ATP present = 0.3 ± 0.1 mM

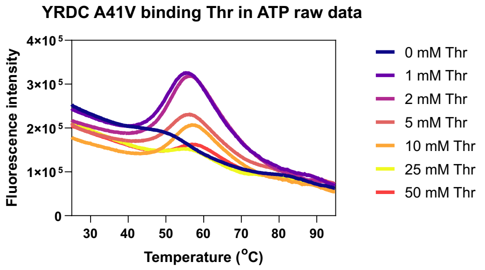

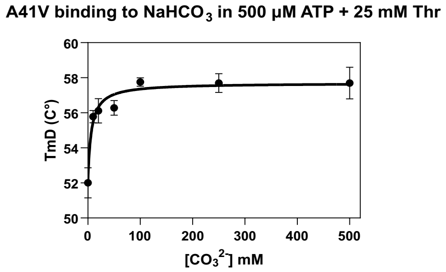

$K_d$ with threonine and ATP present = 6.5 ± 2.0

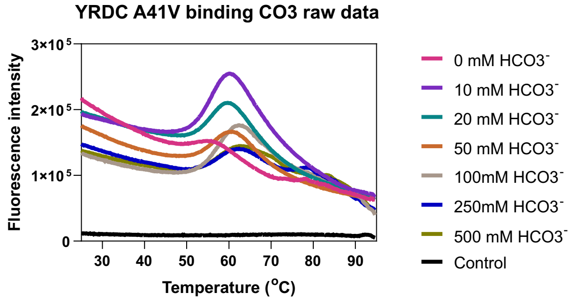

#### YRDC I178T

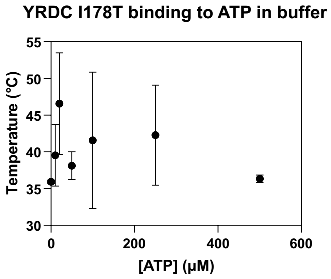

$K_d$ undetermined

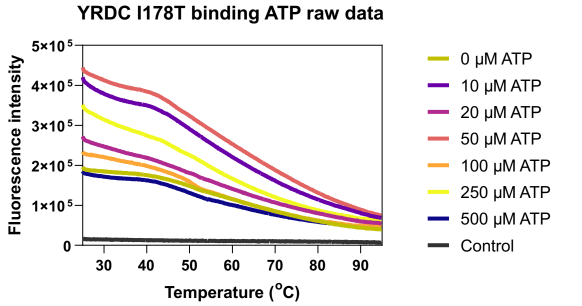

### Supplementary Figure 2: DSF for all proteins with and without ligands

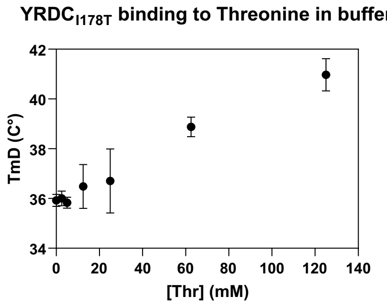

$K_d$ for binding without Threonine >500 µM, too high to accurately determine, trend of increase shown.

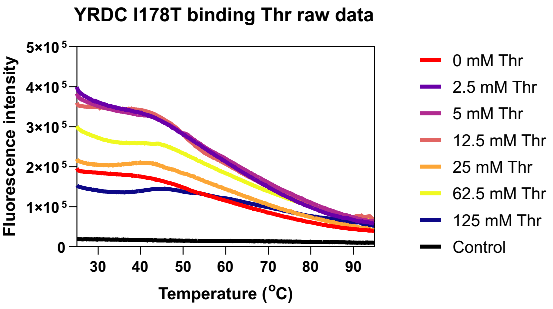

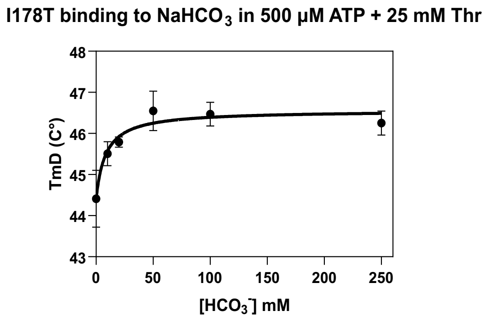

$K_d$ with ATP and threonine present = 8.3 ± 4.2 (mM)

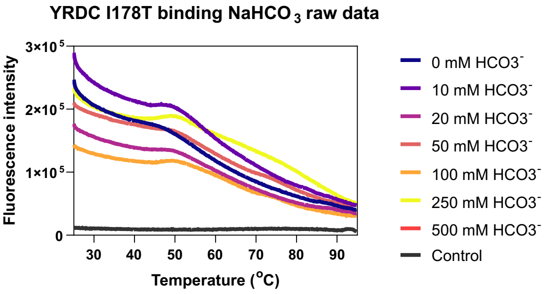

#### YRDC ΔL222

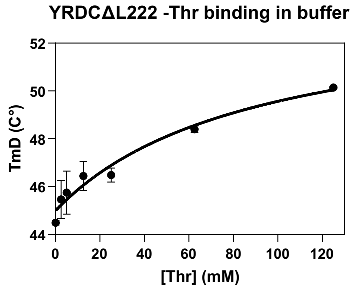

$K_d$ for binding Threonine, too high to accurately determine, estimated value of 89.6 ± 44.9 mM

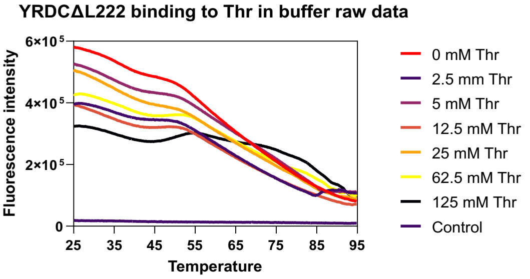

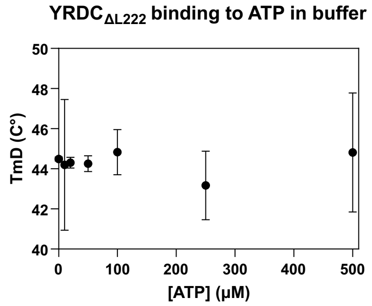

$K_d$ undetermined

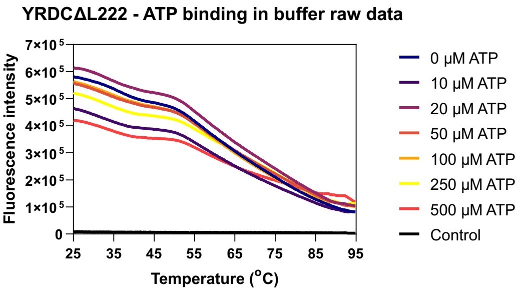

### Supplementary Figure 2: DSF for all proteins with and without ligands

| | |
|---|---|
| 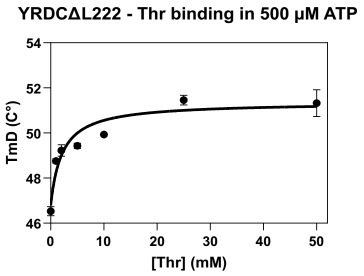<br><br>$K_d$ for binding threonine = 2.1 ± 0.7 mM | 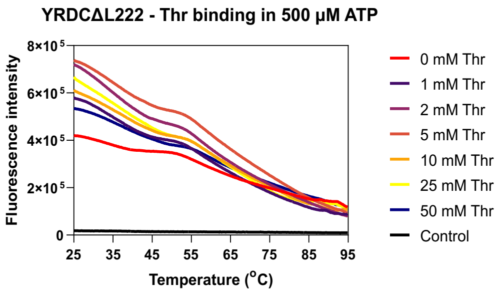<br> |

Supplementary Figure 3: PPi and AMP binding monitored by DSF.

**Binding of reaction products to YRDCWT by DSF/SYPRO Orange.**
Left: Derivative melting temperature ($T_m$) plotted against ligand concentration for pyrophosphate (PPi) and AMP. Points are mean ± s.e.m. (n = 3). Right: Corresponding fluorescence traces (normalized RFU vs temperature). $T_m$ values were obtained from the first derivative of melting curves. Replots shown and data fit to a binding hyperbola to yield pyrophosphate and AMP, both metabolic by-products of the reaction and potential inhibitors, with $K_{D\text{-}PPi}$ = 7.1 ± 0.9 mM and $K_{D\text{-}AMP}$ = 6.1 ± 1.8 mM.

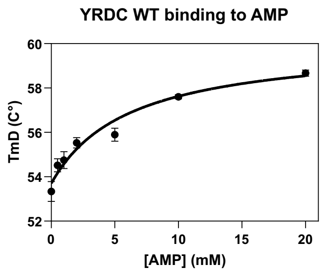

$K_d$ for binding AMP = 6.1 ± 1.8 mM

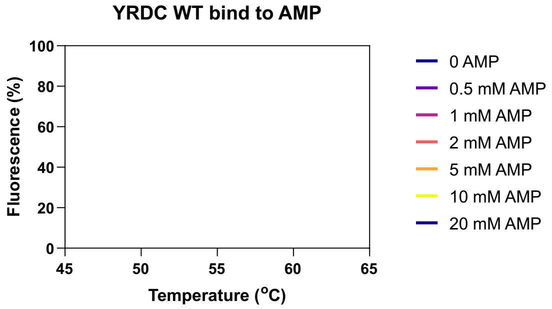

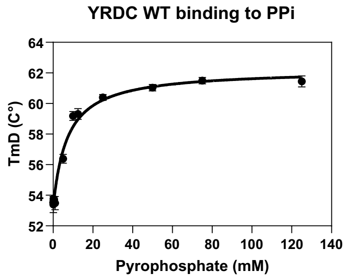

$K_d$ for binding PPi = 7.1 ± 0.9 mM

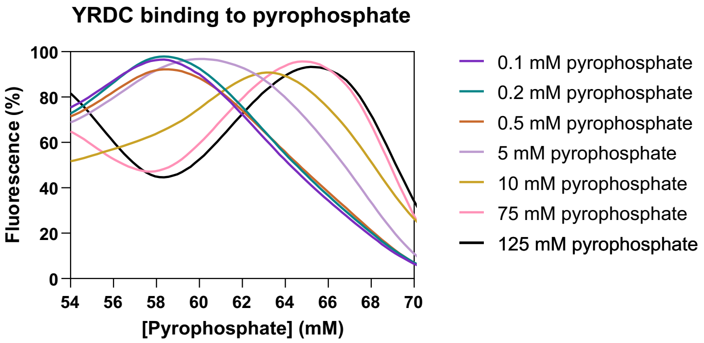

Supplementary Figure 4

Supplementary Figure 4: Circular dichroism spectra (CD) for all protein variants in the absence and presence of ligands as indicated.

Circular dichroism spectra (CD) for all protein variants in the absence and presence of ligands as indicated. Left: Far-UV CD spectra (mdeg vs wavelength, nm) for wild-type (WT) YRDC and all mutants recorded in 20 mM potassium phosphate (pH 8.0), 75 mM NaF (buffer) and in buffer + 500 µM ATP + 50 mM L-threonine (ligands). Right, thermal unfolding traces (mdeg vs temperature) used to extract melting temperatures ($T_m$) fitting procedures are described in Methods.

**YRDC WT**

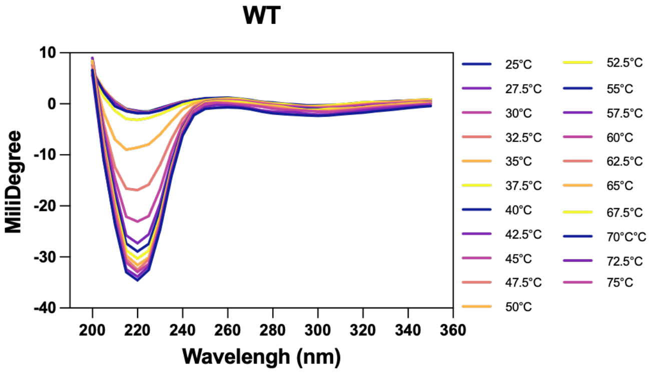

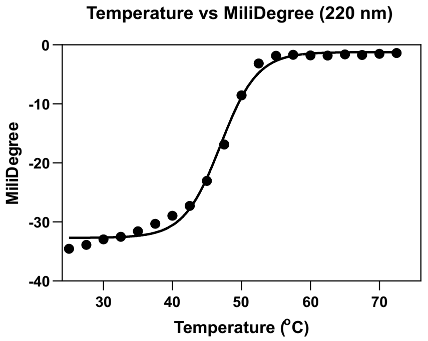

No ligands
$T_m = 47.0 \pm 0.3$

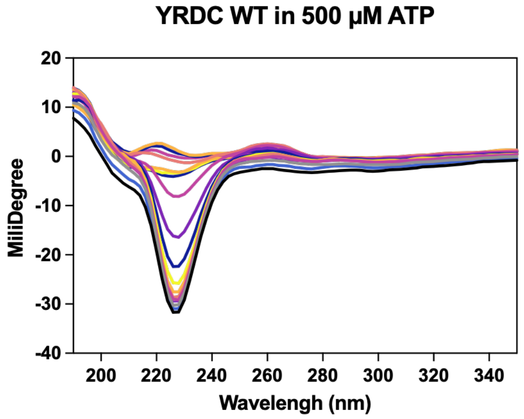

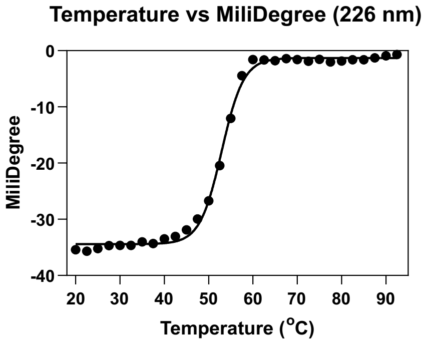

+ 500 µM ATP and 5 mM *L*-threonine
$T_m = 53.0 \pm 0.2$

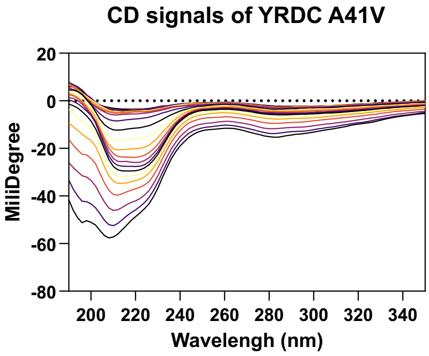

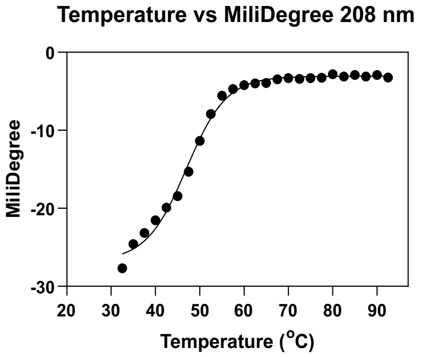

No ligands $T_m = 47.0 \pm 0.5$

### Supplementary Figure 4

**CD signals of YRDC A41V in 500 μM ATP and 5 mM Thr**

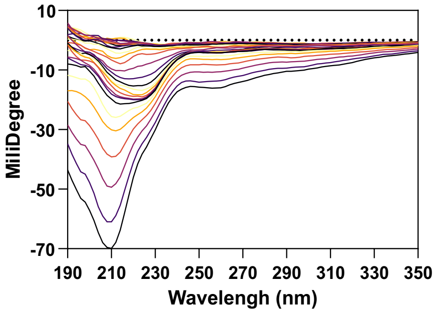

**Temperature vs MiliDegree**

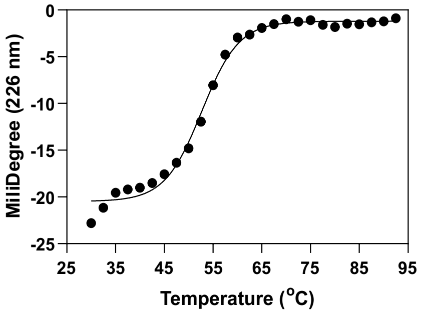

+ 500 μM ATP and 5 mM *L*-threonine
$T_m$ = 52.6 ± 0.4

YRDC T46A in the absence and presence of 500 μM ATP and 5 mM *L*-threonine

**CD signals of YRDC T46A**

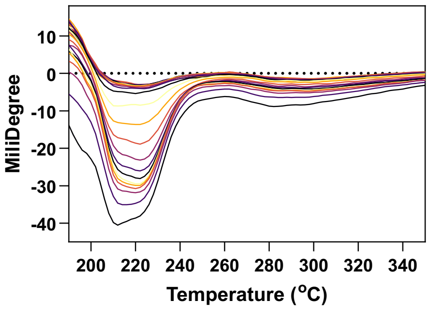

**Temperature vs MiliDegree 212 nm**

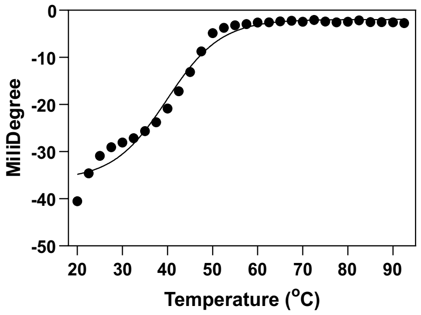

No ligands $T_m$ = 39.9 ± 0.9

**CD signals of T46A in 500 μM ATP and 5 mM Thr**

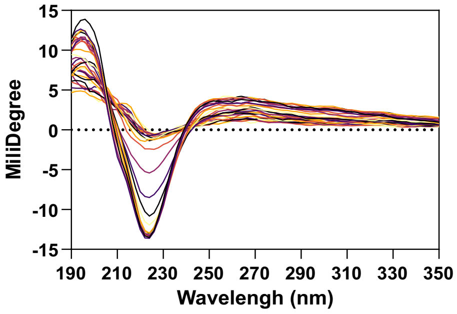

**Temperature vs MiliDegree**

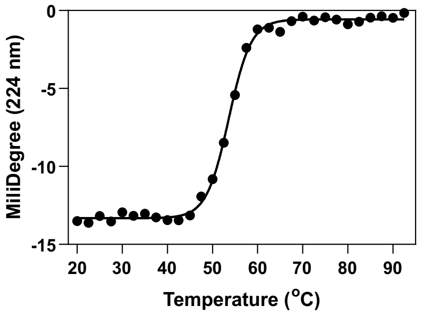

+ 500 μM ATP and 5 mM *L*-threonine
$T_m$ = 53.6 ± 0.1

YRDC C78A in the absence and presence of 500 μM ATP and 5 mM *L*-threonine

**CD signals of C78A**

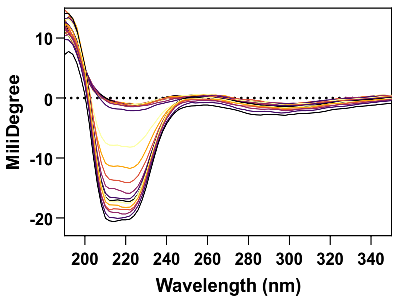

**Temperature vs MiliDegree (214nm)**

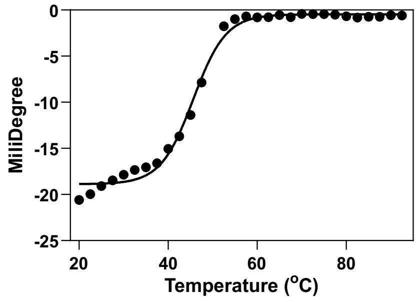

No ligands $T_m$ = 45.6 ± 0.3

### Supplementary Figure 4

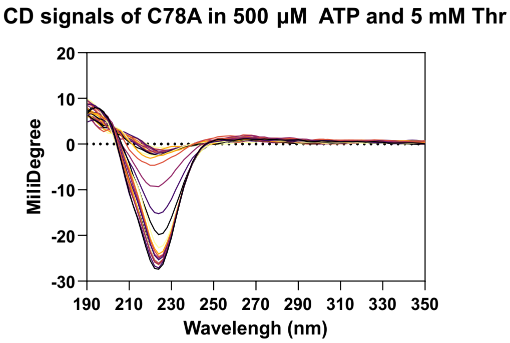

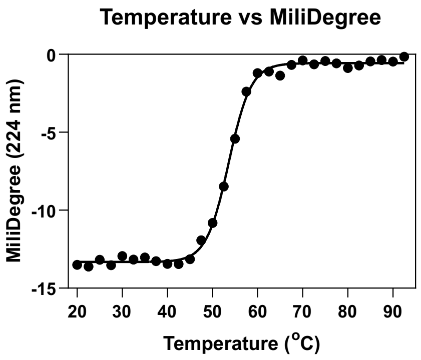

+ 500 µM ATP and 5 mM *L*-threonine
$T_m$ = 52.8 ± 0.2

YRDC R130A in the absence and presence of 500 µM ATP and 5 mM *L*-threonine

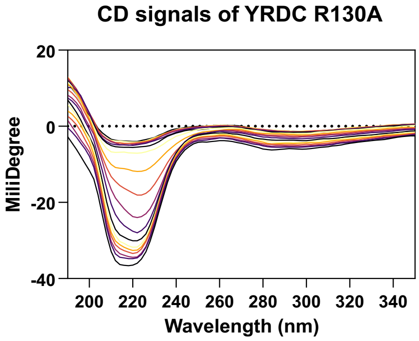

No ligands $T_m$ = 41.2 ± 0.3

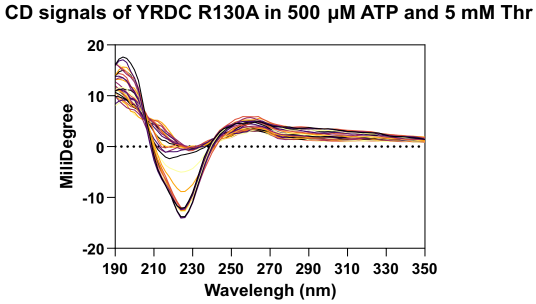

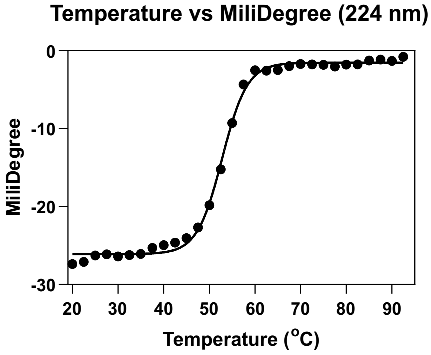

+ 500 µM ATP and 5 mM *L*-threonine
$T_m$ = 46.6 ± 0.2

YRDC I178T in the absence and presence of 500 µM ATP and 5 mM L-threonine

### Supplementary Figure 4

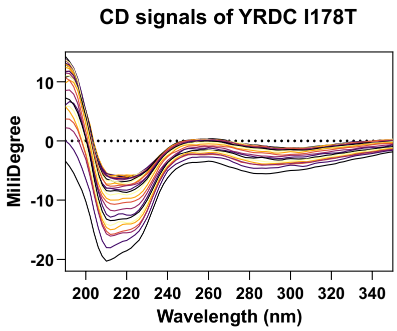

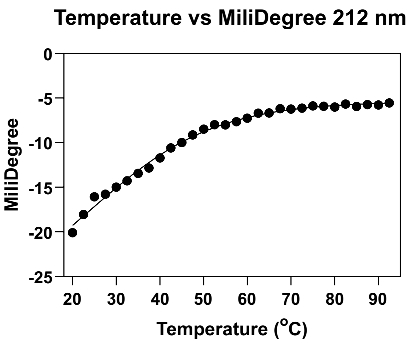

No ligands $T_m = 24.1 \pm 4.7$

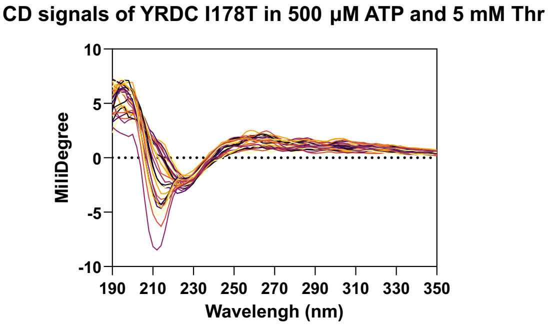

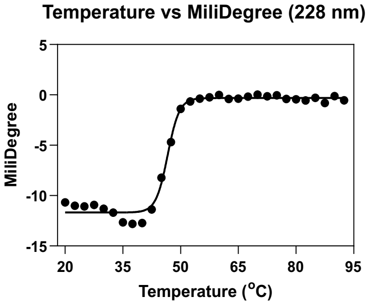

+ 500 μM ATP and 5 mM *L*-threonine
$T_m = 41.3 \pm 1.8$

YRDC S195A in the absence and presence of 500 μM ATP and 5 mM L-threonine

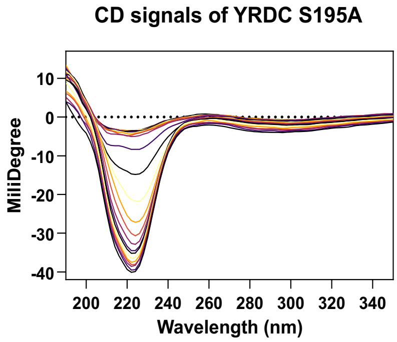

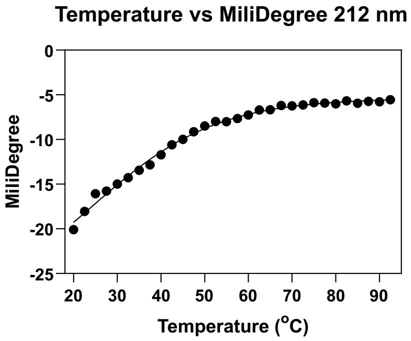

No ligands $T_m = 47.1 \pm 0.2$

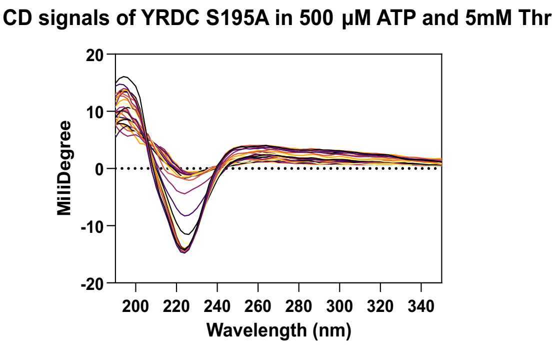

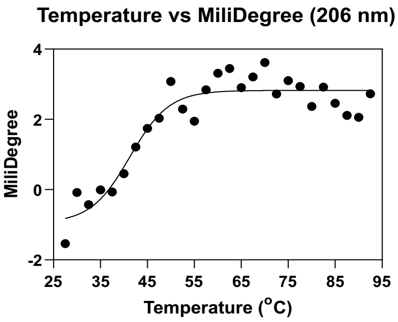

+ 500 μM ATP and 5 mM *L*-threonine
$T_m = 52.8 \pm 0.1$

YRDC R210K in the absence and presence of 500 μM ATP and 5 mM L-threonine

### Supplementary Figure 4

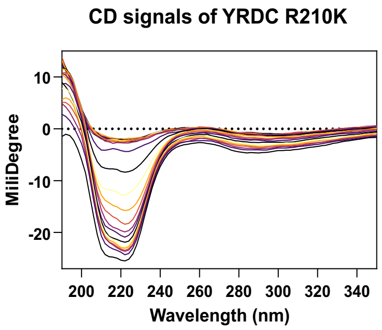

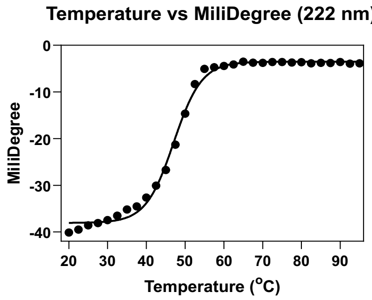

No ligands $T_m$ = 46.6 ± 0.3

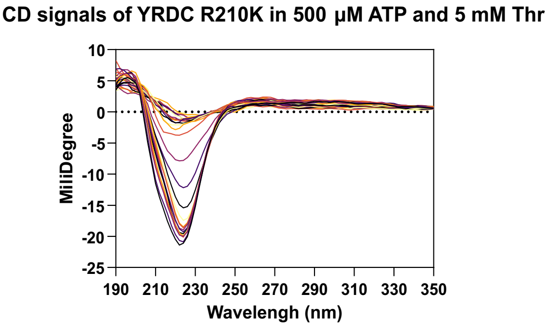

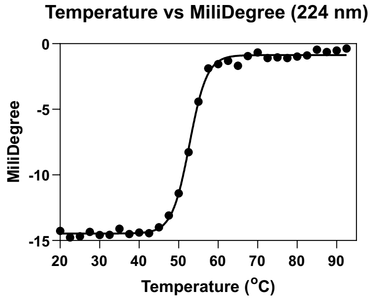

+ 500 μM ATP and 5 mM *L*-threonine
$T_m$ = 53.3 ± 0.2

YRDC ΔL222 in the absence and presence of 500 μM ATP and 5 mM L-threonineYRDC

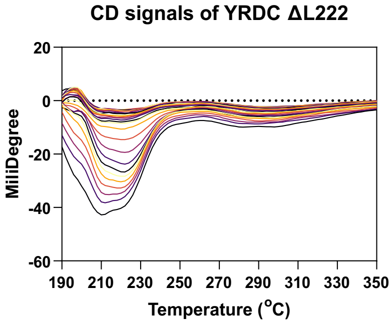

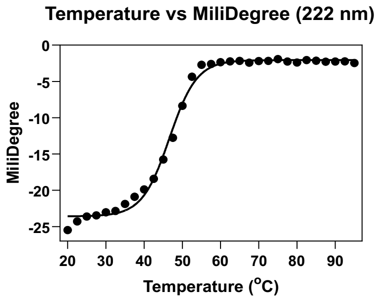

No ligands $T_m$ = 37.0 ± 0.6

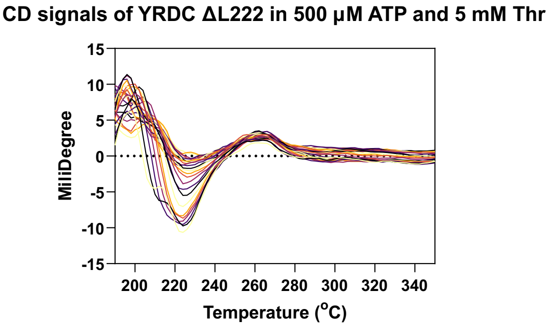

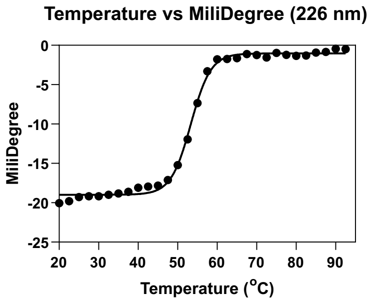

+ 500 μM ATP and 5 mM *L*-threonine
$T_m$ =50.9 ± 0.7

Supplementary Figure 5: Detection of AMP and pyrophosphate (PPi) in the YRDC reaction mixture by $^{31}$P NMR.

Stacked spectra show the complete reaction (ATP + *L*-threonine + $NaHCO_3$) at 37 °C and at room temperature, alongside AMP, PPi, and ATP standards and two controls lacking either L-threonine or $NaHCO_3$. At 37 °C, clear PPi and AMP peaks appear, and ADP is not detected. The absence of PPi/AMP at room temperature and their appearance (PPi) at 37 °C indicate temperature-dependent catalysis by YRDC, consistent with formation of TC-AMP with PPi as the paired product and AMP generated during this step. Both negative controls show only the ATP pattern, indicating that AMP does not arise from nonspecific ATP hydrolysis under these conditions.

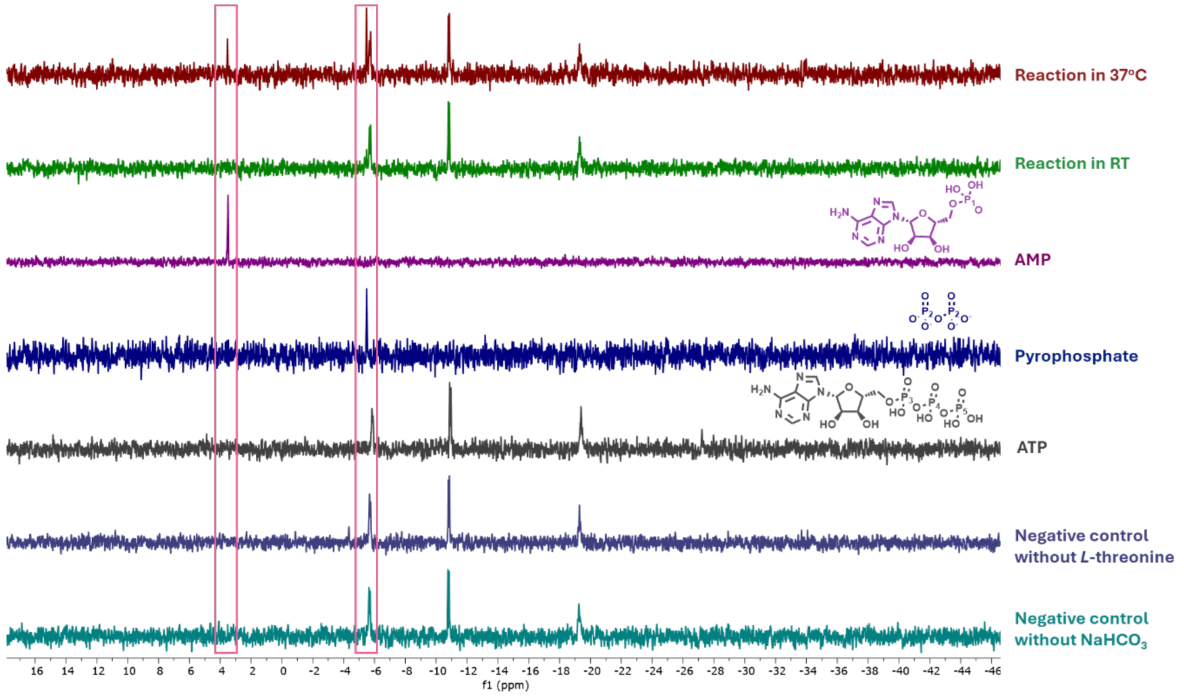

### Supplementary Figure 6: Michaelis Menten curves for all protein variants

Supplementary Figure 6: Michaelis Menten curves for all protein variants

**Michaelis Menten curves for all protein variants with varying ATP.** Left: Representative progress curves for WT and all mutants recorded at increasing [ATP]; AMP formation was quantified by LC–MS and used as a readout of TC-AMP synthesis. Initial rates were obtained from the linear regime of each trace (fit windows defined in Methods). Right: Corresponding Michaelis–Menten plots with single-site fits yielding $k_{cat}$, $K_{\mathrm{m}}$, and $k_{cat}/\mathrm{K_m}$. *L*-threonine and bicarbonate were held at saturating concentrations; only ATP was varied. Points show mean ± s.e.m. (n = 3). Full conditions are provided in Methods.

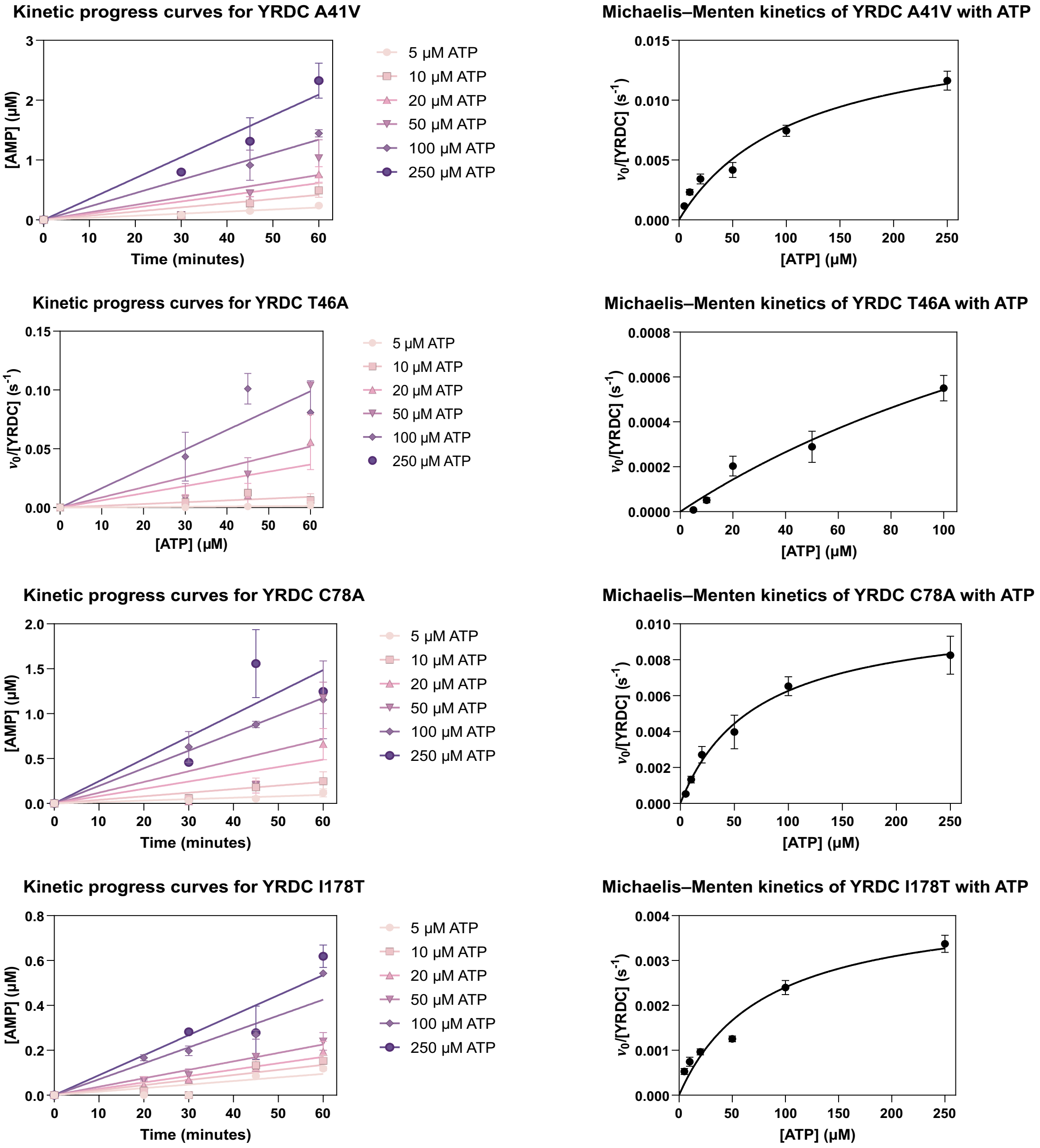

### Supplementary Figure 6: Michaelis Menten curves for all protein variants

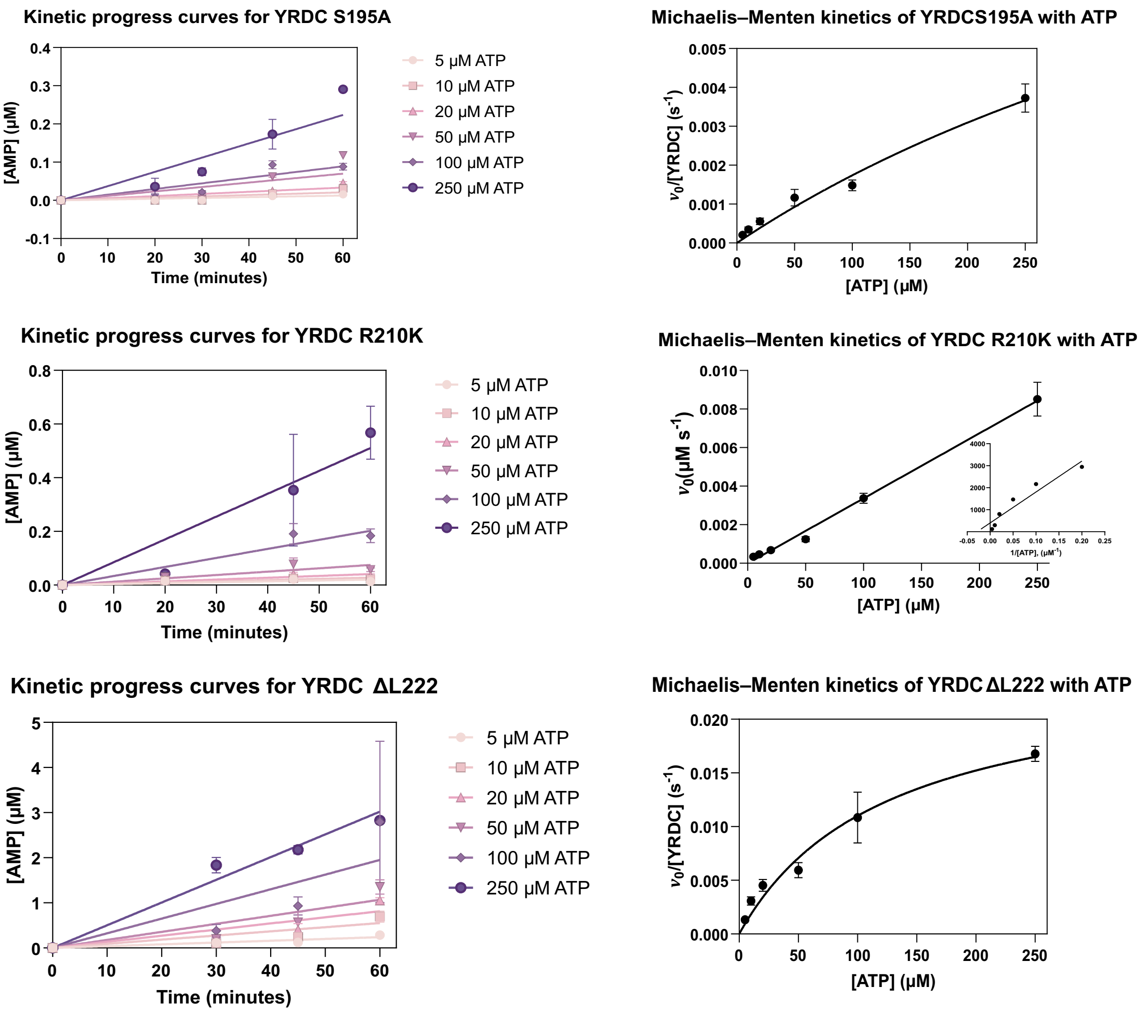

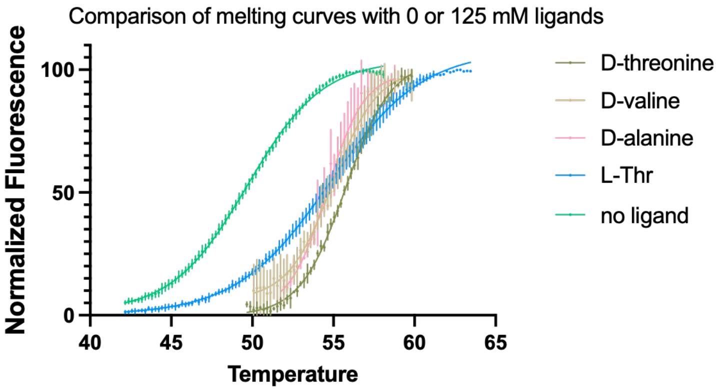

| Ligand | $T_m$ |
|---|---|
| D-threonine | 55.7 ±0.1 |
| D-valine | 54.9 ±0.1 |
| D-alanine | 54.6 ±0.1 |
| L-Threonine | 54.8 ±0.1 |
| No ligand | 49.7 ±0.1 |

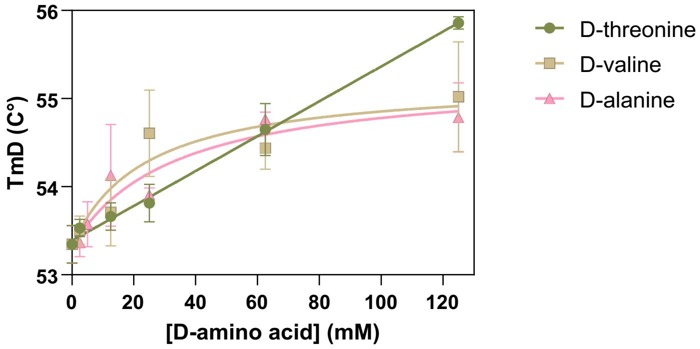

| Ligand | $T_m$ |
|---|---|
| D-threonine | ND |
| D-valine | 23.7 ± 18.3 |
| D-alanine | 33.0 ± 19.2 |

Supplementary Figure 7: YRDC shows weak binding and poor stabilization by D-amino acids. DSF experiments were carried out with 8 mM YRDC and varying concentrations of D-threonine, D-valine and D-alanine. Top shows a comparison between no ligands, and 125 mM D-threonine, D-valine and D-alanine, with fitted values for Tm showing all are capable of stabilizing the enzyme. Bottom shows the replot of Tm values in function of ligand concentration, showing a linear trend for threonine, and hyperbolic curves for other ligands, albeit with large error values for individual measurements (table on the right).

Supplementary Figure 8: LC-HRMS for reaction products with other amino acid substrates.

**LC–HRMS detection of AMP and amino-acid carbamoyl-AMP adducts formed by YRDC.** Shown are extracted-ion chromatograms and accompanying MS[1] spectra for reactions containing ATP + $NaHCO_3$ + one amino acid (L-threonine, L-serine, L-alanine, L-valine, or L-cysteine). Peaks are assigned to the expected carbamoyl-AMP $[M–H]^-$ and $[M+Na–2H]^-$ species.

AMP release measured for WT YRDC converting ATP, *L*-threonine, and $NaHCO_3$ to TC-AMP.

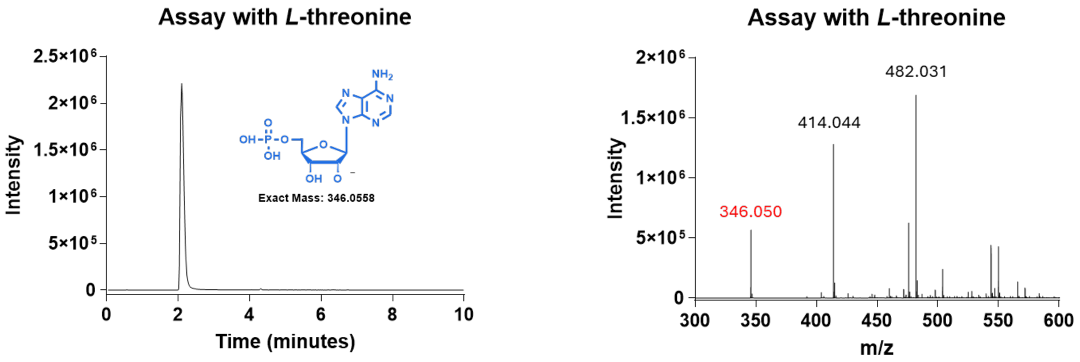

AMP release measured for WT YRDC converting ATP, *L*-cysteine, and $NaHCO_3$ to TC-AMP.

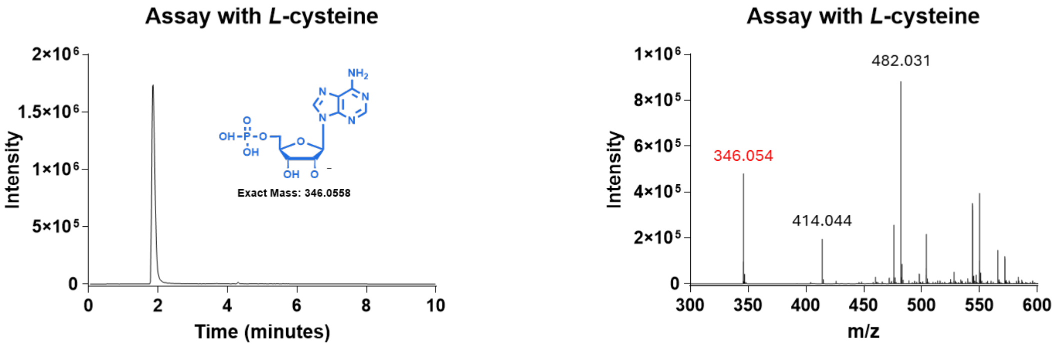

Signal of alanylcarbamoyl -AMP

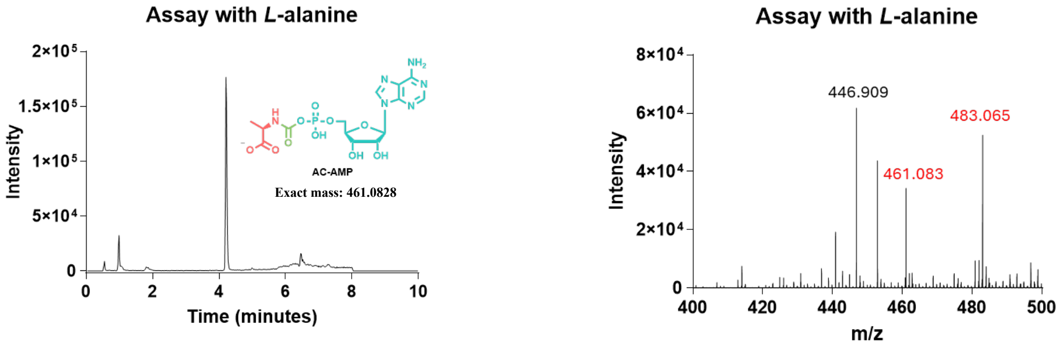

AMP release measured for WT YRDC converting ATP, *L*-alanine, and $NaHCO_3$ to TC-AMP.

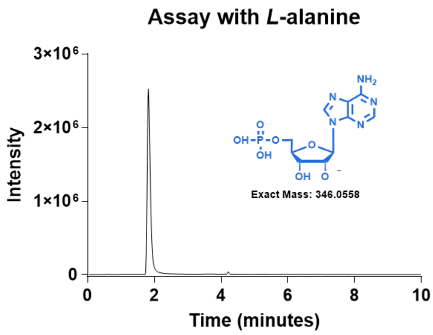

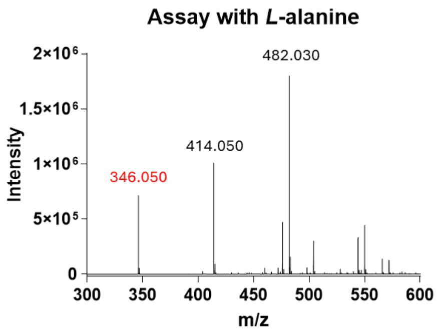

Signal of serylcarbamoyl -AMP

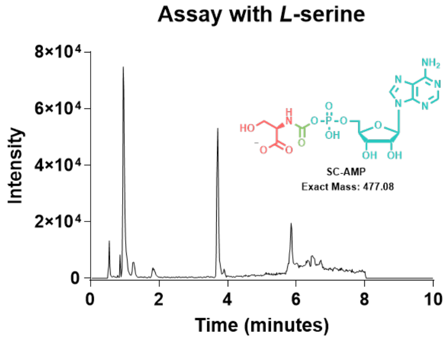

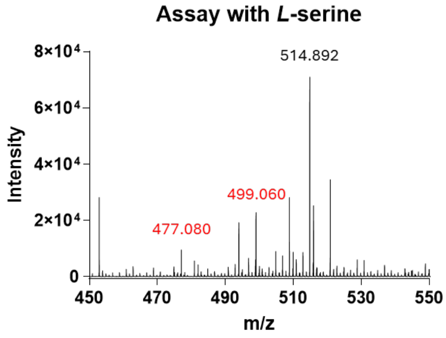

AMP release measured for WT YRDC converting ATP, *L*-serine, and $NaHCO_3$ to TC-AMP.

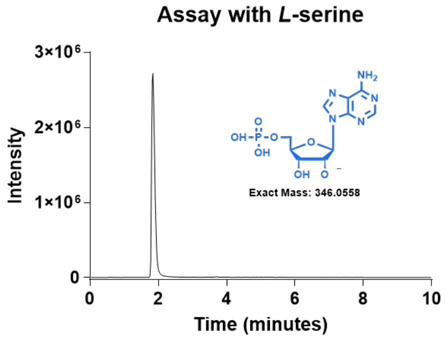

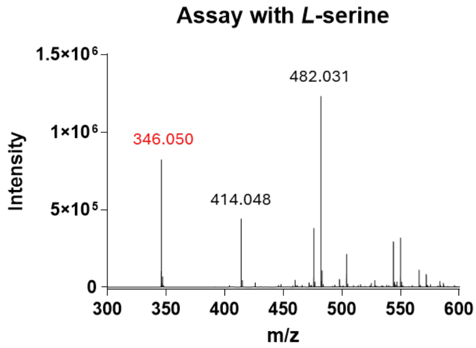

Signal of valylcarbamoyl-AMP

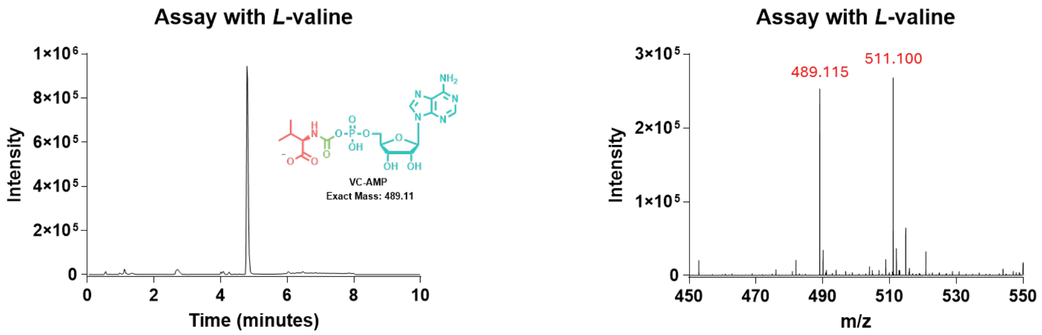

AMP release measured for WT YRDC converting ATP, *L*-valine, and $NaHCO_3$ to TC-AMP.

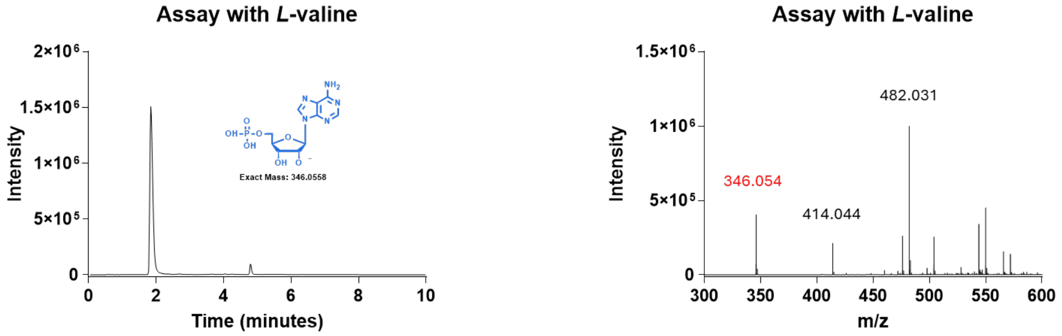

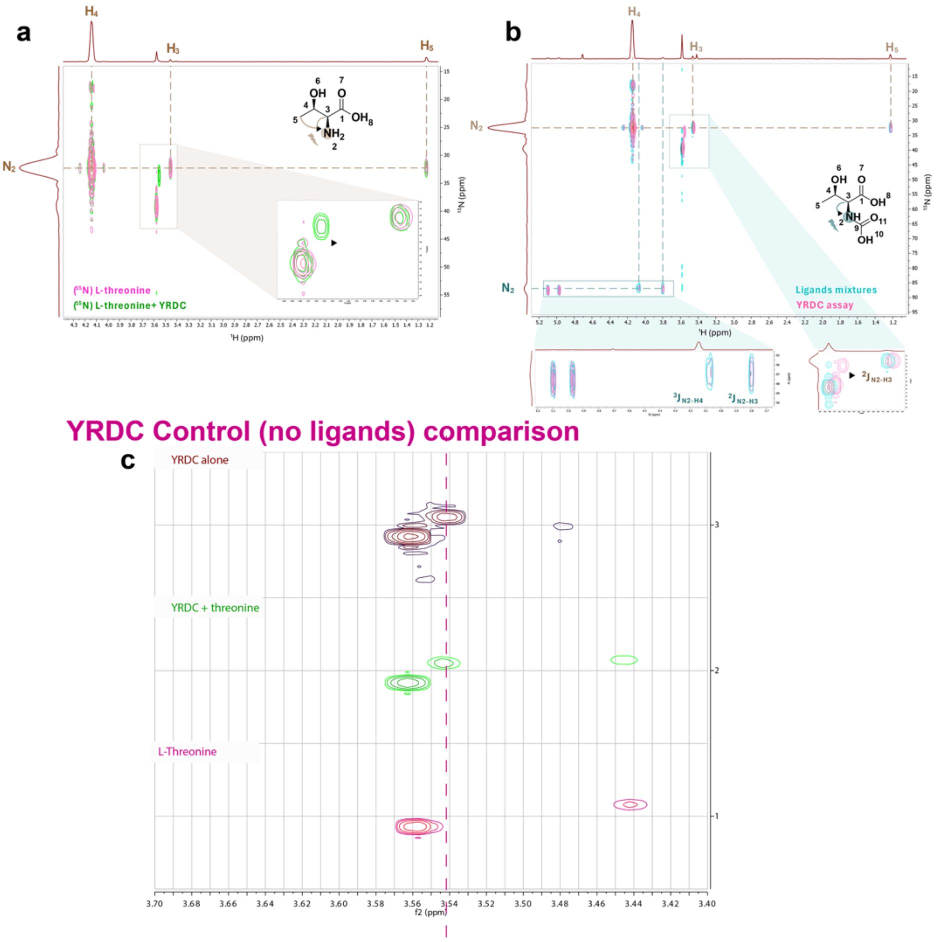

Supplementary Figure 9: Threonine–carbamate adduct forms to the same extent in the presence and absence of YRDC.

a) [$^{15}$N]-L-threonine showed $^{1}$H resonances at δ 1.22 (H-5), δ 3.43 (H-3), and δ 4.13 (H-4) correlating to δN ≈ 31.94 ppm. Presence of peak with δ 3.55, later attributed to tricine buffer which interacts with enzyme (panel c)

b) With all substrates present ([$^{15}$N]-L-threonine, [$^{13}$C] $NaHCO_3$, ATP; 37 °C, 2 h – cyan traces), the $^{1}$H–$^{15}$N HMBC spectrum contained multiple features diagnostic of a threonine–carbamate. Long-range correlations were observed from H-3 (δ 3.80) and H-4 (δ 4.07) into an amide-region nitrogen at δN 86.93 ppm, and the $^{1}$H–$^{13}$C HMBC showed H-3 (δ 3.80) coupling to a carbonyl at δC 164.11 ppm, consistent with a carbamate carbonyl. Two additional $^{1}$H–$^{15}$N cross-peaks were detected at the same nitrogen chemical shift (δN 86.93 ppm) with $^{1}$H signals at δ 4.97 and δ 5.10. These features are explained by isotopomers that arise in $D_2O$: the carbamate exists predominantly as a deuterated amide species and to a lesser extent as a protonated amide species. The long-range H-3 and H-4 correlations at δ 3.80 and δ 4.07 originate from the major deuterated amide isotopomer ($^{2}J/^{3}J_{N\text{-}H}$), whereas the pair of stronger cross-peaks near δ 5.0 correspond to one-bond $^{1}J_{N\text{-}H}$ correlations from the minor protonated amide isotopomer. One-bond correlations are intrinsically more intense than long-range correlations, which accounts for their appearance despite the lower population of the protonated amide isotopomer. Pink traces: [$^{15}$N]-L-threonine, [$^{13}$C] $NaHCO_3$, ATP; 37 °C, in the presence of YRDC for 2h. No changes in the relative abundance for the threonine–carbamate adduct can be observed. No chemical shift differences are observed and sample is identical to what was observed in the absence of enzyme.

c) YRDC in the absence of ligands (top) shows a peak corresponding to the bicarbonate buffer (δ 3.55 ppm), which is a remnant of purification despite buffer exchange prior to this experiment.

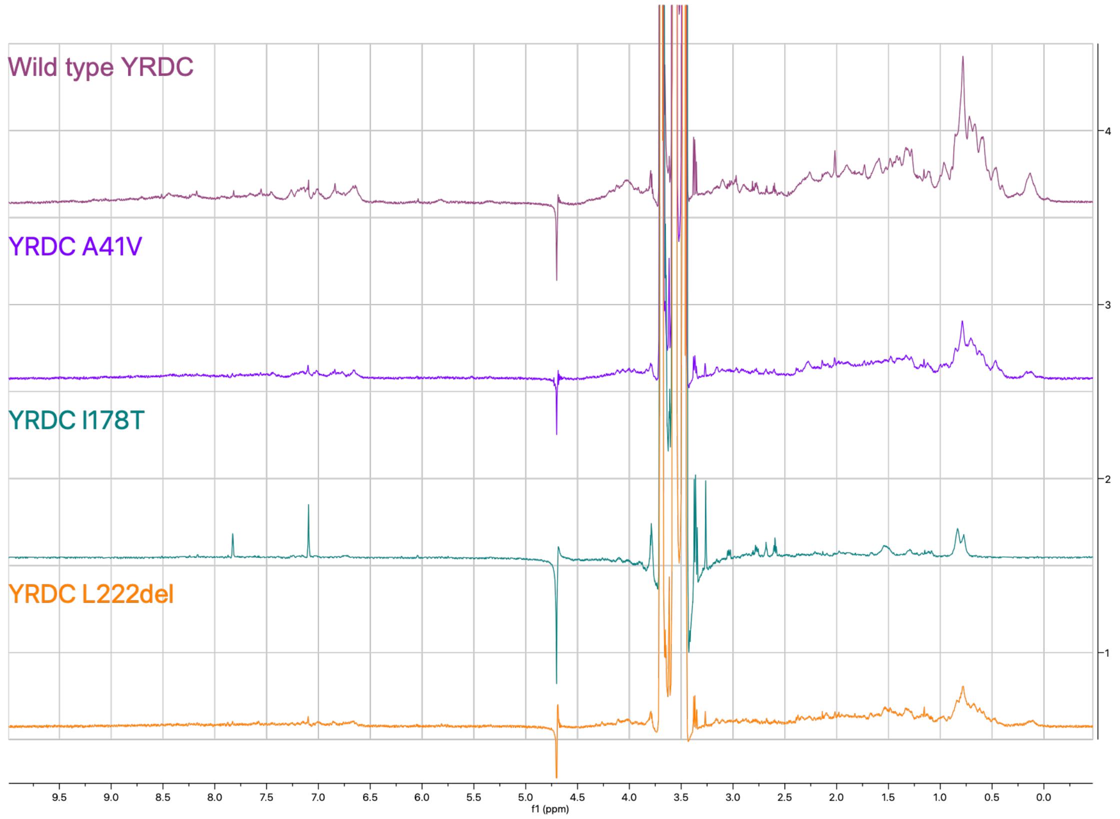

Supplementary Figure 10: $^{1}H$ NMR of WT and GAMOS mutants. Each protein variant was buffer exchanged into 20 mM potassium phosphate pH 8.5 with 100 mM KCl. Spectra were acquired with 10% $D_2O$, to reveal a similar degree of folding when comparing WT and A41V variants. The L222 deletion and I178T mutants show significantly less well-folded protein, with decrease of peaks between 0-1 ppm, characteristic of solvent-protected CH groups, as well as sharper and fewer peaks between 6.5-9 ppm, which is characteristic of amide groups which are solvent exposed.

Supplementary Figure 11: Native Mass spectrum of YRDCA41V and YRDCΔL222

**Native Mass spectrum of YRDC$_{A41V}$ and YRDC$_{\Delta L222}$.** Spectra were recorded in 50 mM ammonium acetate (pH 6.8). Colored dots mark assigned charge-state series; numbers above peaks indicate charge state ($z^+$). Top, A41V. Light green denotes monomer, green dimer, and purple trimer. A41V redistributes signal away from the WT-like dimer into monomer and higher-order species. Insets show zoomed regions of representative dimer and trimer envelopes. Bottom, ΔL222. Pink denotes monomer and purple dimer. For both variants, deconvolution yields neutral masses consistent with the sequence-derived monomer and its multiples within instrument accuracy, supporting assignment of the observed oligomers.

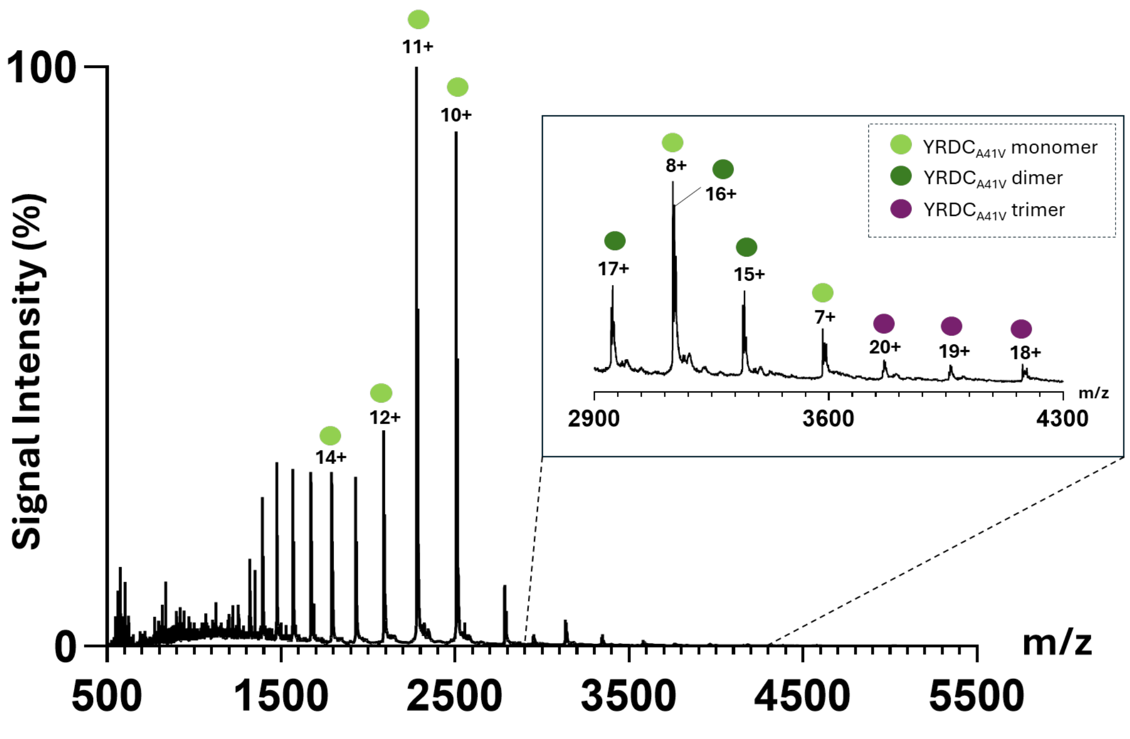

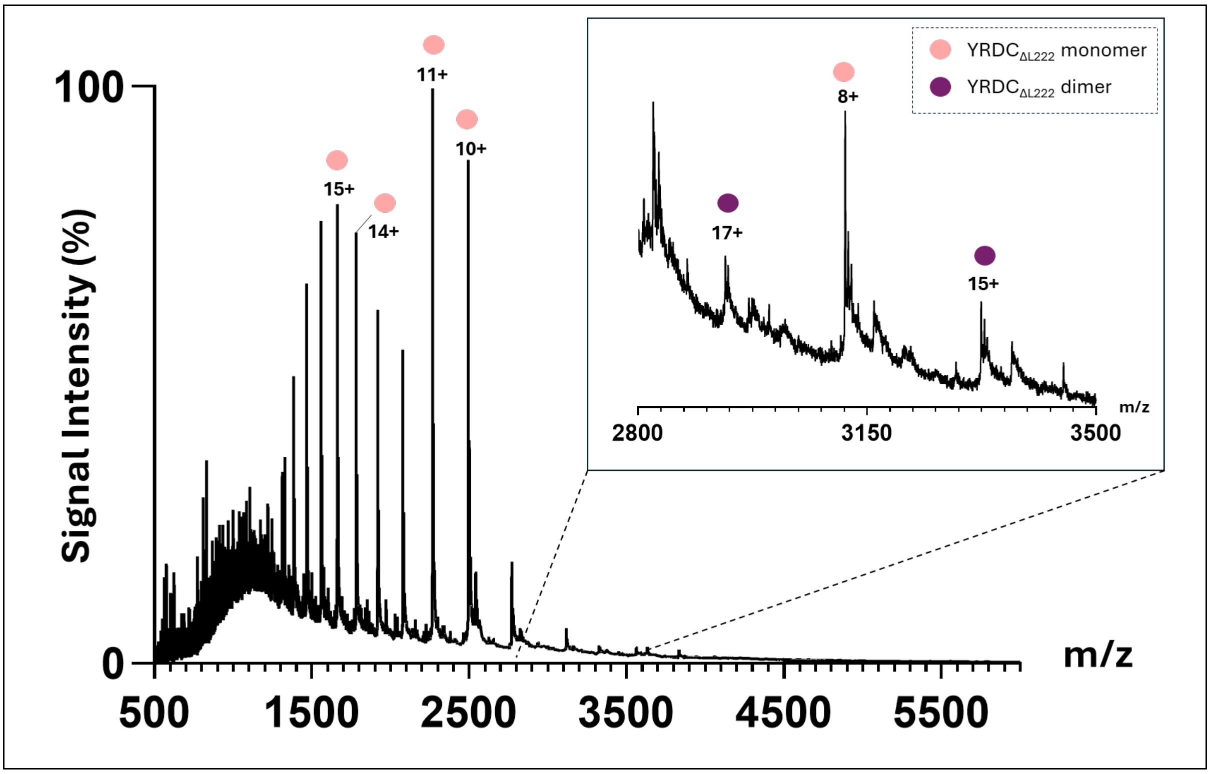

100
0
Signal Intensity (%)
500
1500
2500
3500
4500
5500
m/z
15+
14+
11+
10+
YRDC$_{\Delta L222}$ monomer
YRDC$_{\Delta L222}$ dimer
17+
8+
15+
2800
3150
3500
m/z

### Supplementary Figure 12: Mass photometry of YRDC and GAMOS variants in the presence of ligands.

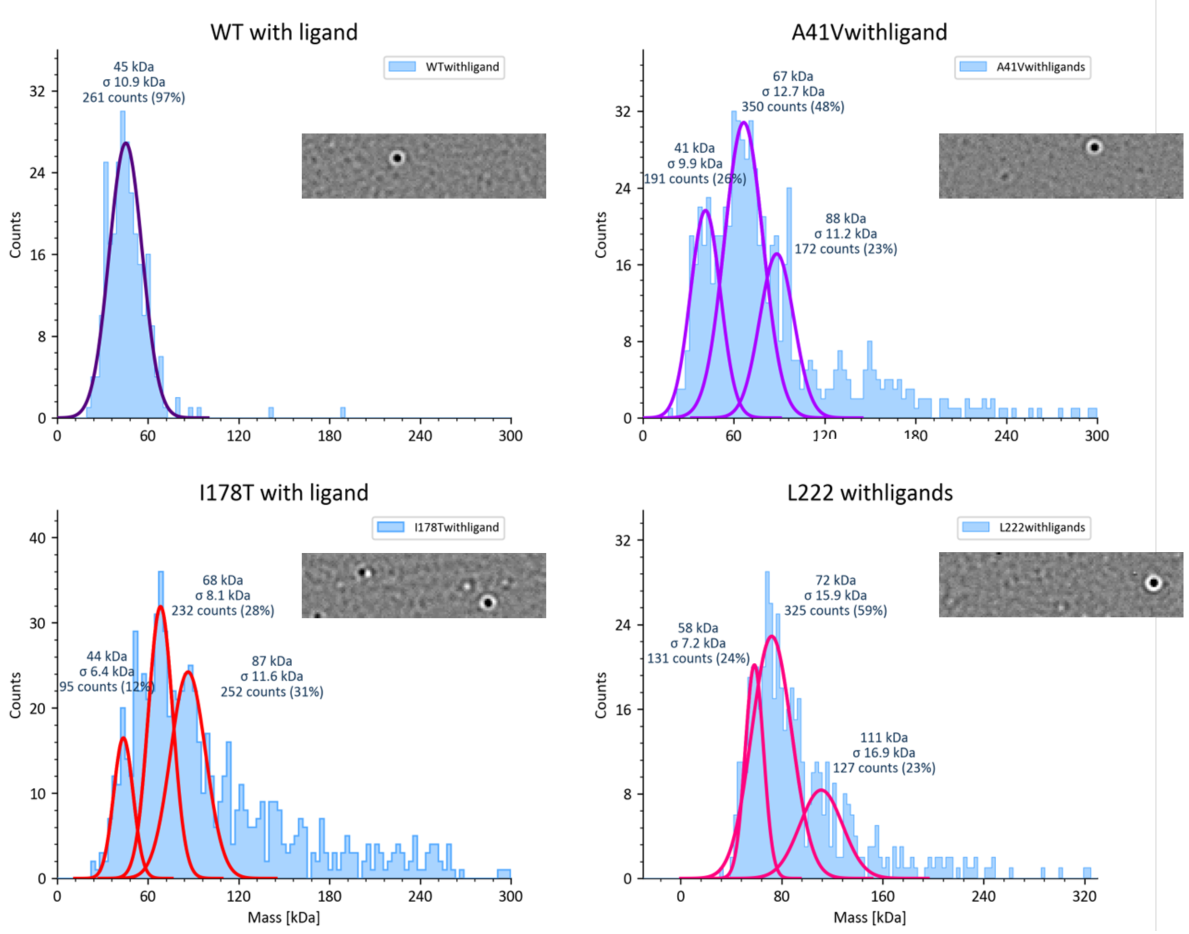

Supplementary Figure 12: Mass photometry of YRDC and GAMOS variants in the presence of ligands.
Representative single frames from 180-s movies (insets) and corresponding mass distributions (histograms) are shown for YRDCWT, YRDCA41V, YRDCI178T, and YRDCΔL222 recorded in reaction buffer with ATP (500 µM) and L-threonine (50 mM) at room temperature. Under these conditions, WT is dominated by a single species at 45 ± 10.9 kDa (≈ dimer), with no detectable higher-order peak. YRDCA41V resolves into multiple populations near ~41, ~67, and ~88 kDa, consistent with redistribution from the native dimer toward higher oligomers. YRDCI178T shows the lowest dimer fraction, with broad populations around ~44, ~68, and ~87 kDa. YRDCΔL222 is enriched in larger assemblies, with peaks around ~58, ~72, and ~111 kDa. Together, these measurements indicate that the WT enzyme is predominantly dimeric, whereas GAMOS-linked variants shift the assembly toward non-dimeric species under identical solution conditions.

Supplementary Figure 13: Crosslinking with BS3 to probe dimer interface

a) SDS-PAGE after crosslinking with BS$^3$. Bands analysed by mass spectrometry are highlighted with a red box.

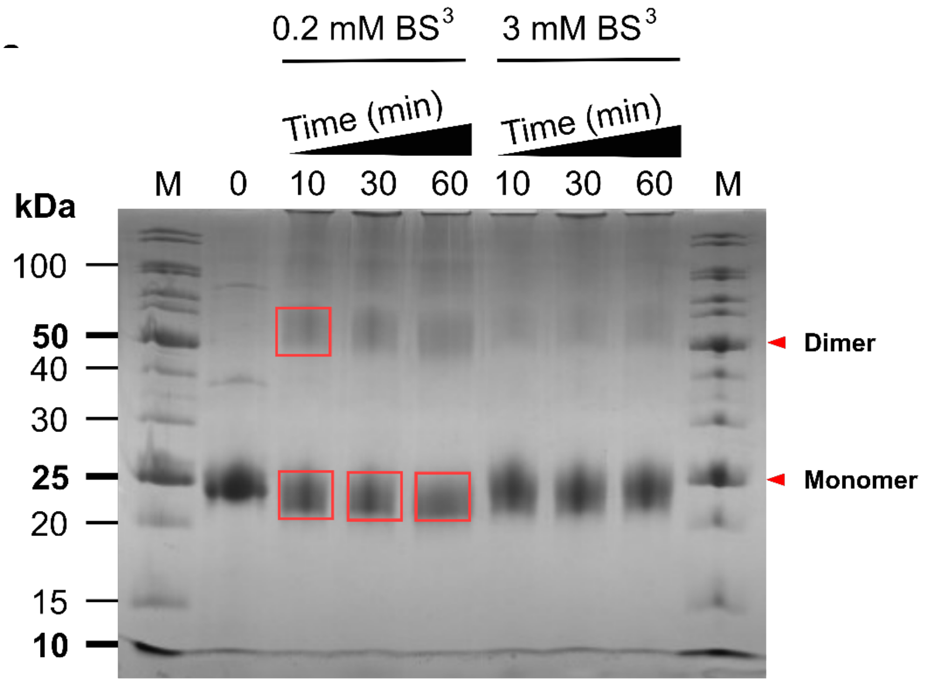

b) Band corresponding to dimer: peptides identified to be crosslinked. Crosslinking with BS3 showing less confidence Alphafold dimer predictions and locations of detected crosslinks. Only one of the predicted dimer interfaces is in agreement with crosslinking data (top, model 2). Calculated distances between crosslinked residues are shown.

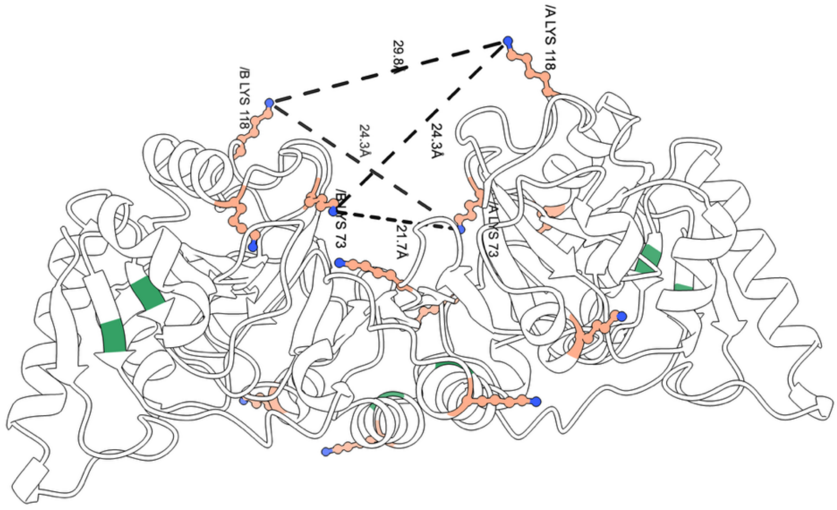

Alpha fold model 2 – top agreement with crosslinking data.

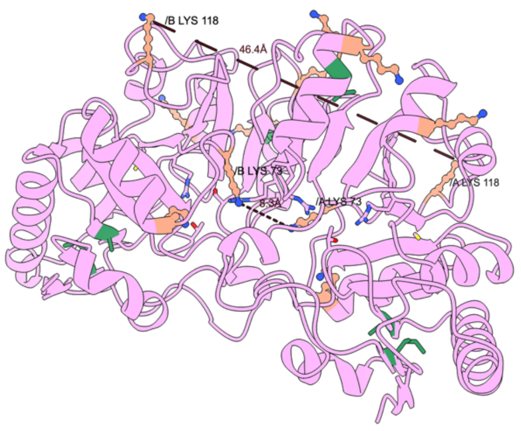

Alpha fold Model 3
(r.m.s.d. compared to Model 2 = 0.397 Å)

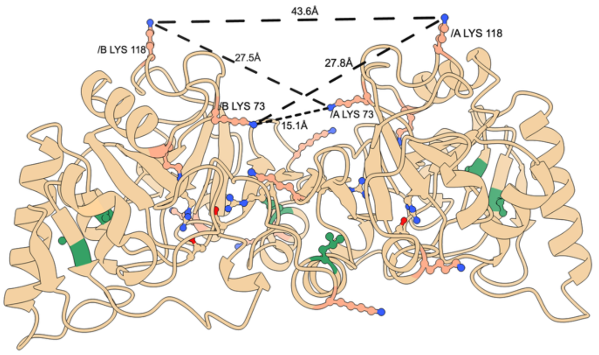

Alpha fold Model 1
(r.m.s.d. compared to Model 2 = 0.273 Å)

c) Summary of identified crosslinks.

| Index | Number of 1st Residue | Number of 2nd Residue | |
|---|---|---|---|
| 1 | 73 | 67 | Also on monomer |
| 2 | 118 | 67 | Also on monomer |
| 3 | 73 | 73 | |
| 4 | 99 | 67 | |
| 5 | 205 | 73 | |
| 6 | 118 | 73 | |
| 7 | 99 | 73 | |
| 8 | 118 | 118 | |

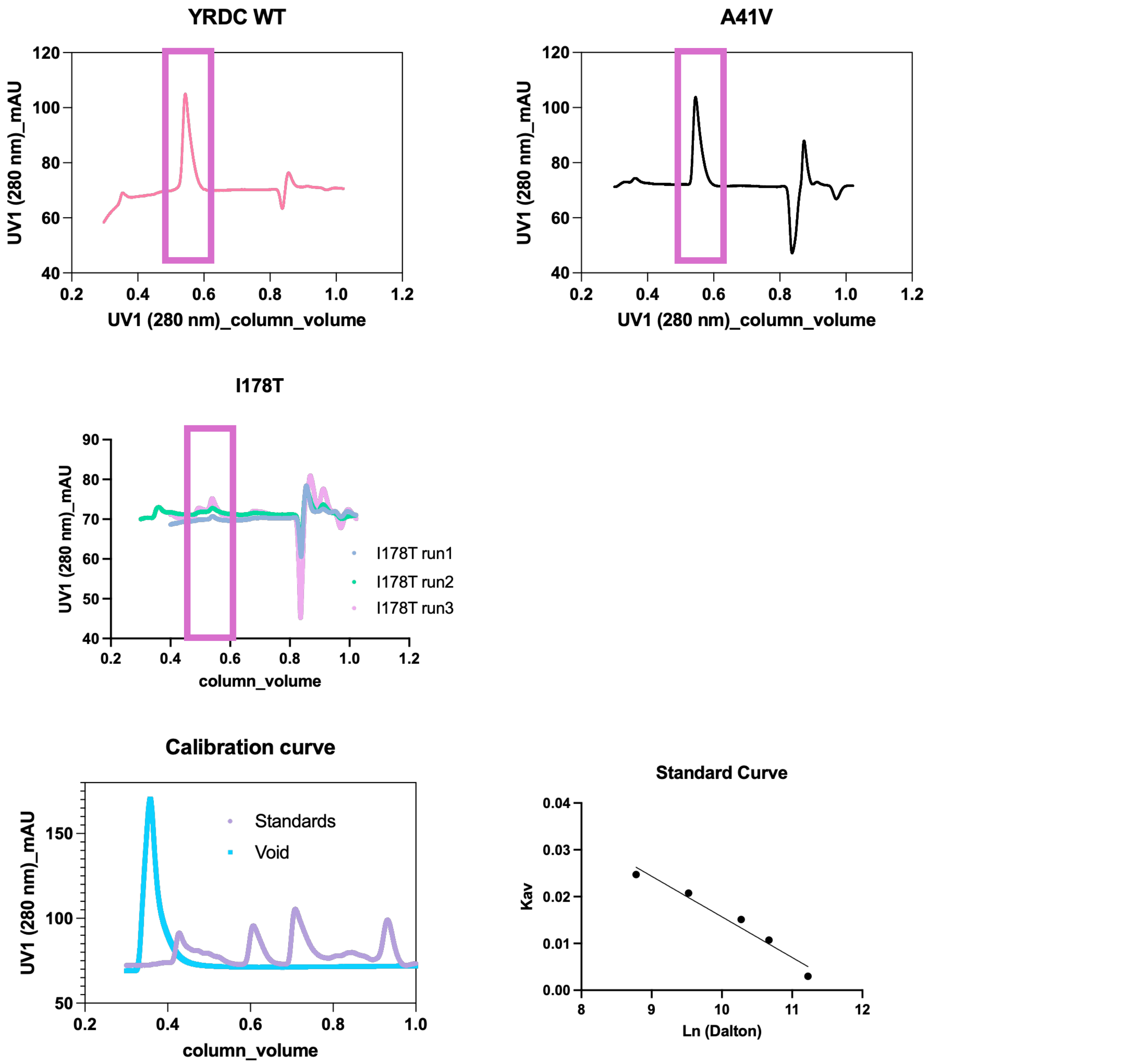

Supplementary Figure 14: Size exclusion chromatography comparing WT and GAMOS variants. Molecular weight was estimated using standards (bottom left), and applying a calibration curve to convert elution volumes into molecular weights (bottom right). Elution volumes corresponding to dimeric species are shown in a purple box. It was not possible to ascertain the molecular weight of I178T and L222 could not be concentrated enough to perform this experiment.

### Supplementary Figure 15: Sequence and structural alignment between human YRDC and homologues

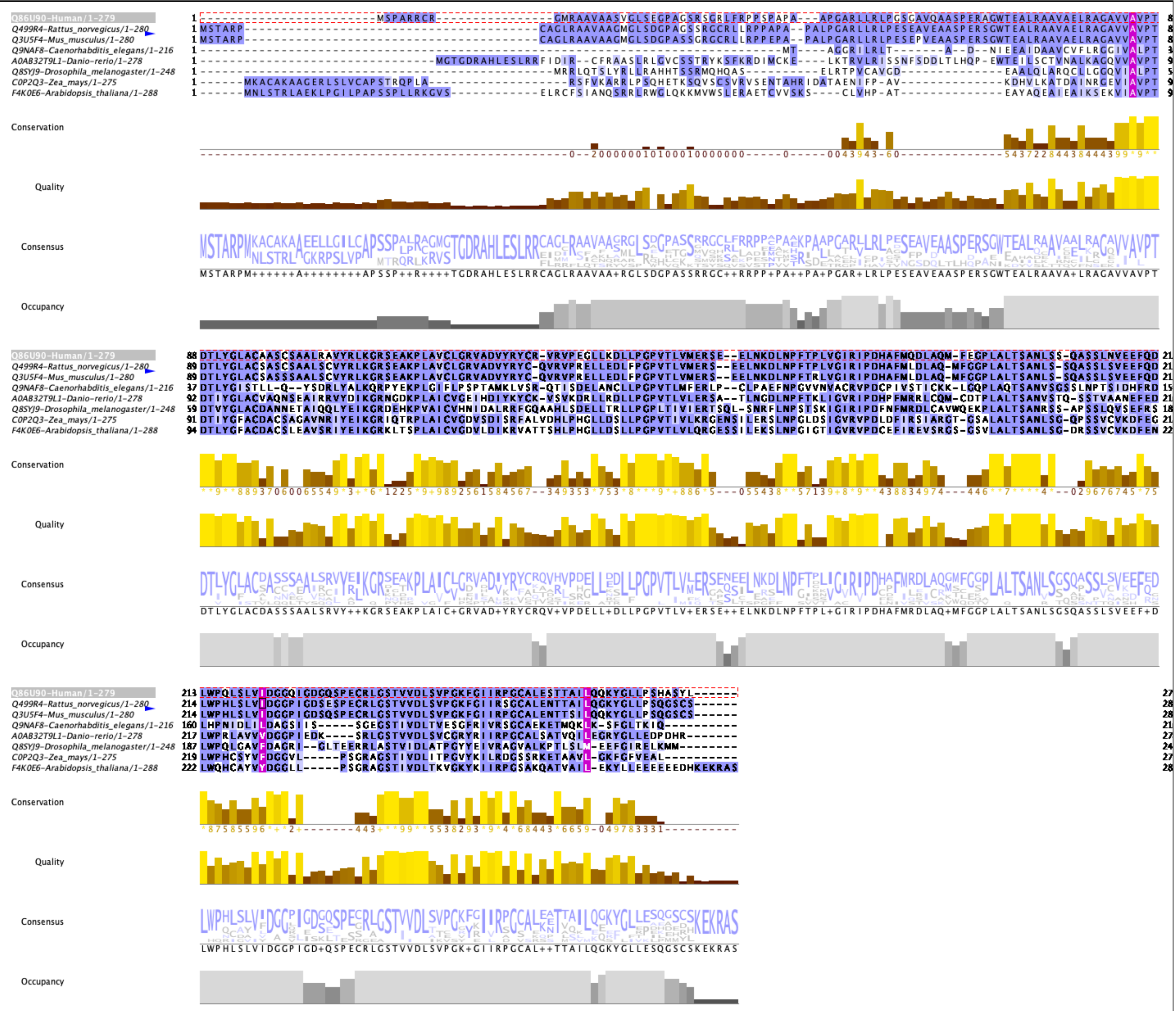

a) Sequence alignment carried out using Clustal omega[3], and visualized with Jalview[4]. Sequences coloured by BLOSUM32 score, and residues marked in pink correspond to equivalent positions to the human enzyme on positions A41, I178 and L222.

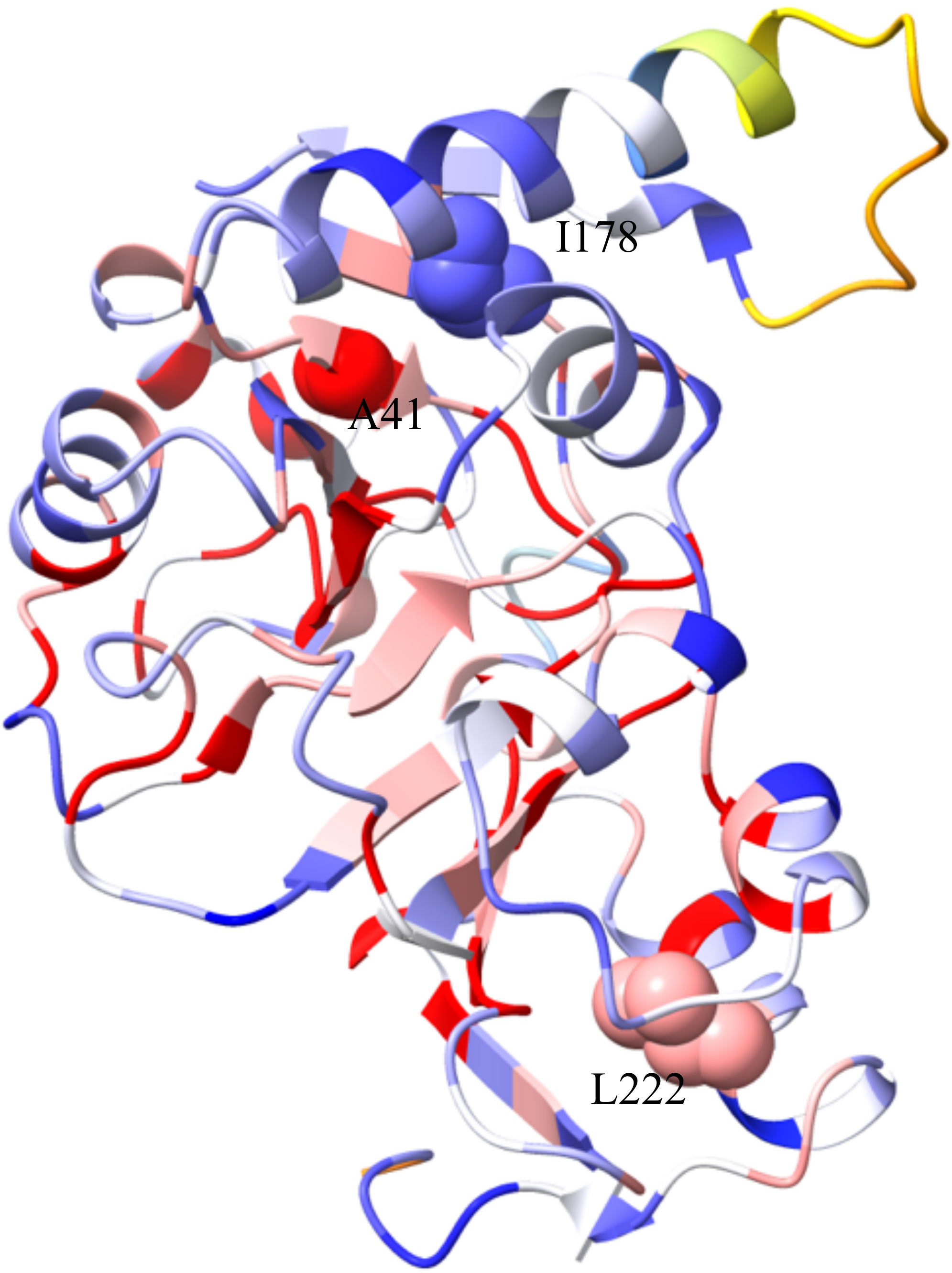

b) Human YRDC coloured by sequence conservation using the alignment above. Highly conserved residues are shown red, highly variable in blue. Yellow regions are gaps between different aligned sequences. GAMOS related positions A41, I178 and L222 are shown as spheres.
